# Canal number and configuration are predictors of external root morphology

**DOI:** 10.1101/2022.04.19.488788

**Authors:** Jason J. Gellis

## Abstract

Within tooth roots canals can vary in shape and configuration, and it is not uncommon for a single root to contain multiple canals. Externally, root morphology also varies, though the range of variation, and its relation to canals remains little explored. This investigation of modern human post-canine teeth uses data from computerized tomography scans of a global sample of 945 modern humans to identify the most frequent phenotypes of root and canal morphologies, and investigate how canal number, shape, and configuration relate to external root morphology. Results (1) include descriptions and definitions of root and canal morphologies, counts, and configurations; (2) indicate that certain canal counts, morphologies, and configurations can predict external morphologies; and (3) that this pattern varies in individual teeth and roots in the maxilla and mandible.

## Introduction

Radiographs, Cone Beam Computed Tomography (CT), and micro CT (µCT) technology have revealed, in ever increasing detail, that the complexity of the root canal system does not always correspond with external tooth root morphology, and that teeth can have multiple canals and canal configurations within a single root (Abbott, 1984; Ahmed et al., 2017; Gellis & Foley, 2021; Kupczik et al., 2019; Kupczik & Hublin, 2010; Moore et al., 2015, 2016)(Figure 1). A recent study has shown that canal number can predict root number, and that the relationship between canal and root number differs between tooth types (i.e., molars and premolars), within and between the maxilla and mandible (Gellis & Foley, 2022). This study investigates the relationship and variability between root canal configuration and external root morphology of fully developed, adult post-canine teeth in a global sample of modern humans (n = 945) from several osteological collections. Specifically, (1) what is the relationship between canal number and configuration to external root morphology; (2) does this relationship vary by tooth; and (3) can root canal number and configuration predict external root morphology?

**Figure 1:**
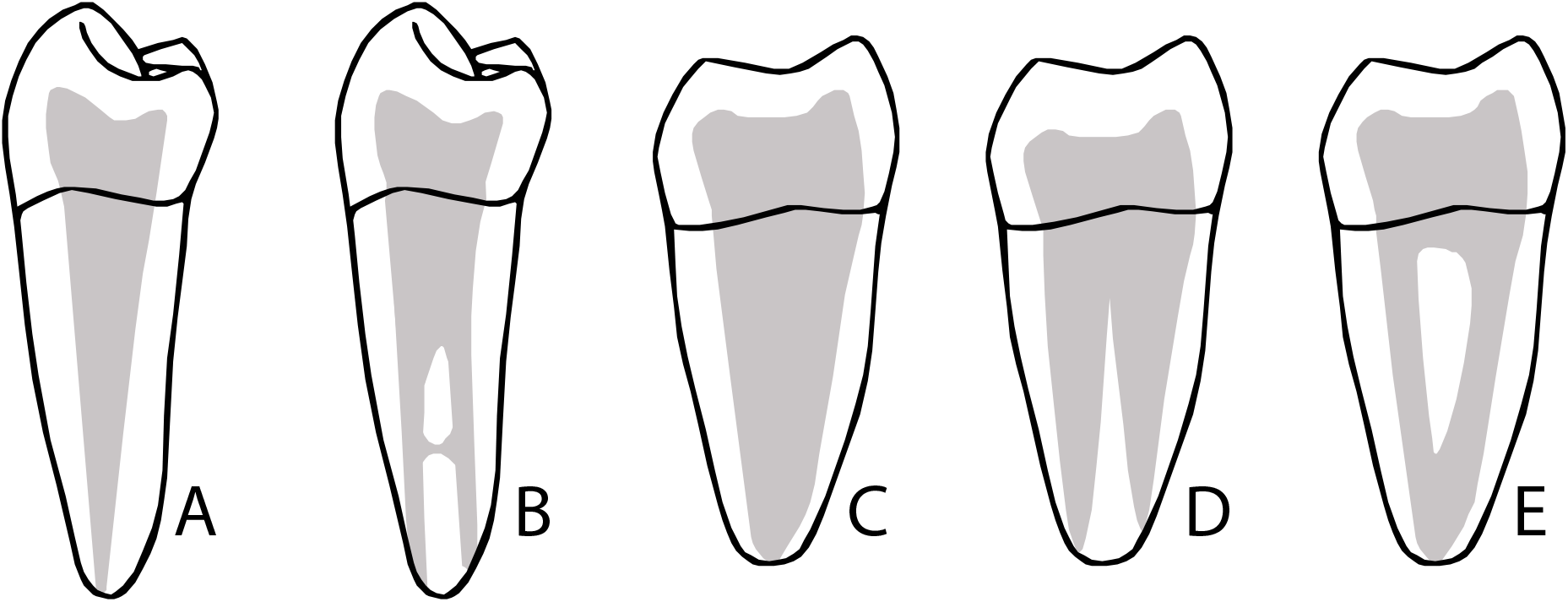
Left - Canal number and morphology does not always follow external number and form. Grey area represents pulp chambers and various canal configurations. A and B are single-rooted mandibular premolars (distal view); C, D, and E are two-rooted mandibular molars (mesial view).

### Development of tooth roots

Tooth root development can be split into two phases: the eruptive and penetrative (Figure 2). The eruptive phase commences when roots begin to develop and ends when the tooth crown is in occlusion. The penetrative phase begins after the tooth crown is in occlusion and ends when the apices of the tooth root complete formation. Both phases can be seen on the root surface — during the eruptive phase, the root surface is smooth while the surface formed during the penetrative phase is rough (Kovacs, 1971). Further, the proportion of smooth to rough surface appears vary among species, with the smooth surface decreasing from carnivore to herbivore (Kovacs, 1971). Amongst modern humans, the proportion is generally two-thirds smooth to one-third rough (Kovacs, 1971).

**Figure 2:**
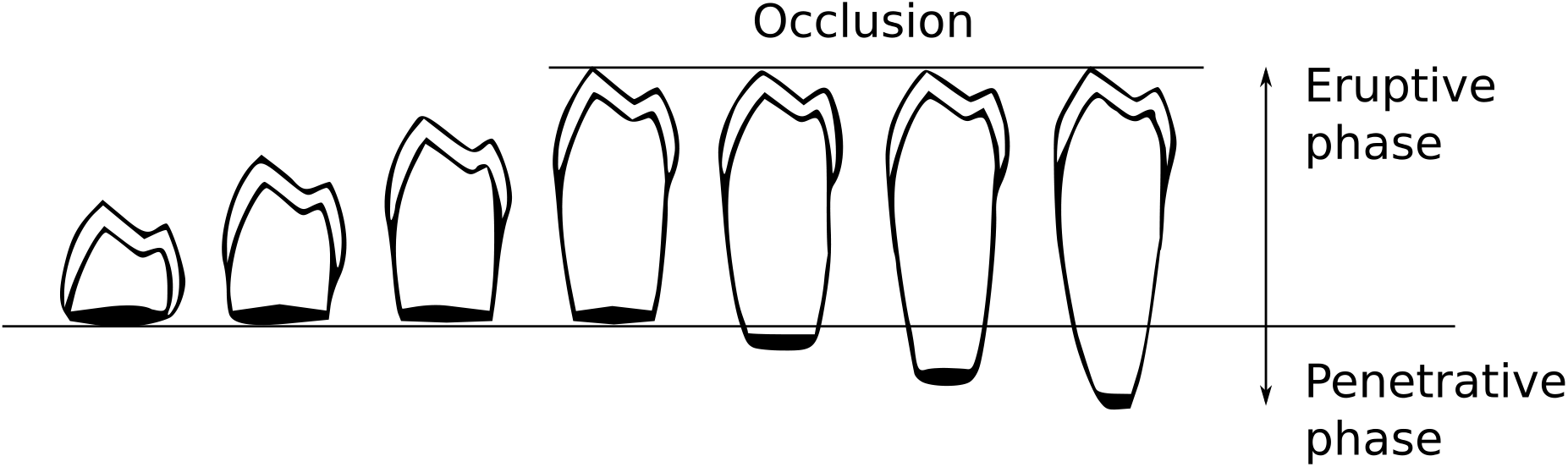
Eruptive and Penetrative phases of tooth root development. Figure modified from Kovacs (1971).

Upon completion of the tooth crown the cervical loop forms a double layer of cells known as Hertwig’s epithelial root sheath (HERS). It is HERS that determines length, curvature, and thickness of roots (Luder, 2015). HERS extends around the dental papilla covering all but the basal portion where it forms an epithelial diaphragm over the apical foramen of the developing root. As the dental papilla expands HERS encases it to form the architecture of the root. Inside the root sheath, ameloblasts induce odontoblasts in the dental papilla to form dentin, which forms the bulk of the root. Simultaneously, mesenchymal cells in the dental papilla differentiate into cementoblasts and secrete cementoid to the external surface of the root sheath. The secreted cementoid gradually matures into a smooth, calcified cementum. During this process the epithelial diaphragm remains in a stationary position relative to the inferior and superior border of the mandible and maxilla respectively. Once the tooth is in occlusion the epithelial diaphragm no longer remains stationary. Instead, it begins to extend towards the base of the alveolar socket and the apices of the root begin to close.

### Teeth with more than one root

As in single rooted teeth, multi-rooted teeth have eruptive and penetrative phases, and extension of HERS, dentin and cementum formation all follow the same basic developmental and physiological processes. During the eruptive phase, as the epithelial diaphragm begins narrowing, ‘tongue like’ extensions called inter-radicular processes (IRP) caused by differential growth rates in HERS begin to divide the primary apical foramen (Figure 3). As these IRPs of the epithelial diaphragm contact opposing IRPs, they fuse and divide the epithelial diaphragm into multiple new openings. The epithelial diaphragm surrounding the opening to each root continues to form at an equal rate of growth. Deviations in the process potentially lead to variations in root morphology (e.g., Taurodontism, supernumerary roots, and pyramidal-shaped roots). The rate at which HERS narrows also determines the length of the root – if HERS narrows rapidly, the root will be shorter; if HERS narrows slowly then the root will be longer (Jafarzadeh et al., 2008).

**Figure 3:**
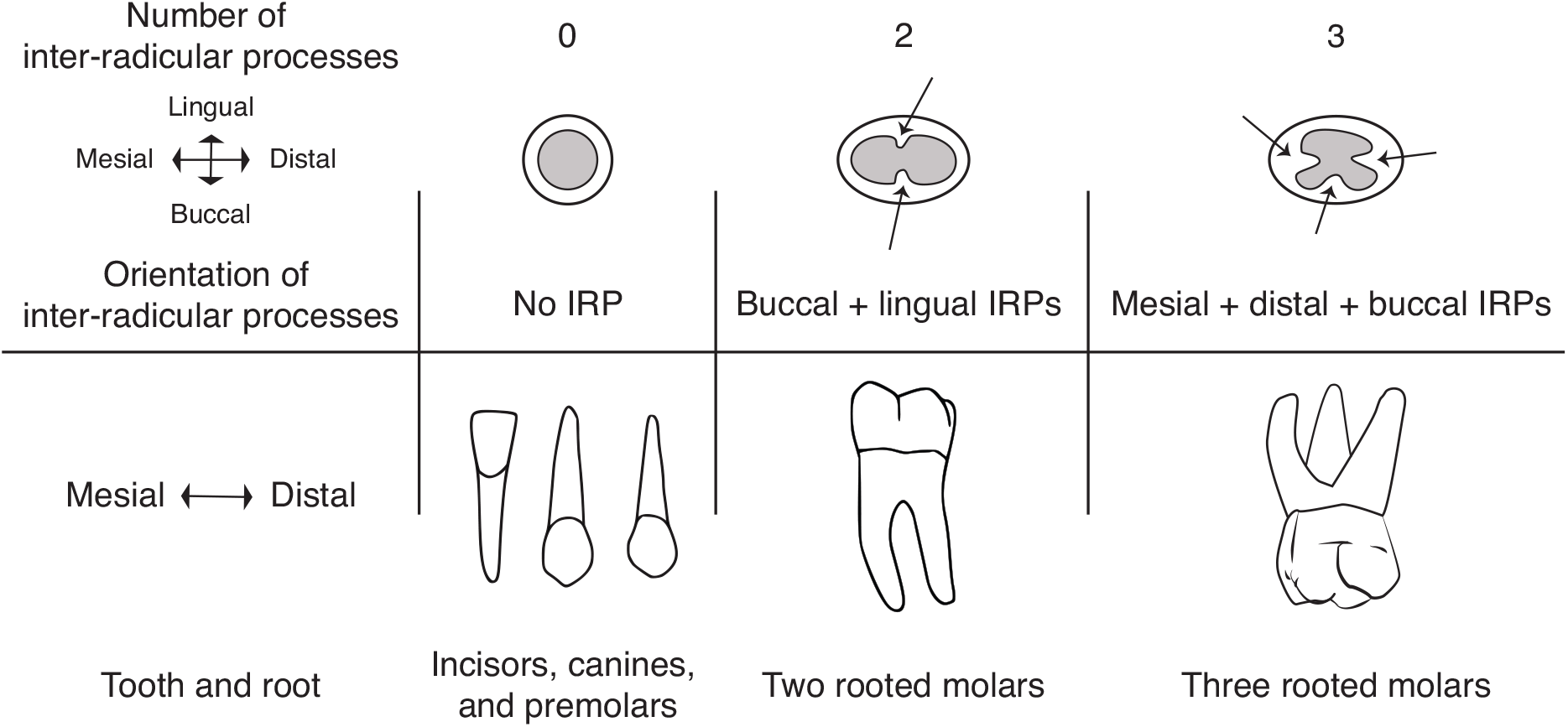
**Top**: Sites on the apical foramen of the developing tooth crown. The location of inter-radicular process (arrows) divides the apical foramen determines the orientation of each tooth root/tooth roots. For example, in a tooth with mesial, distal roots, and buccal roots (top right) three inter-radicular processes arise from the mesial, buccal, and distal borders of the apical foramen, forming mesial, buccal, and distal secondary apical foramina upon fusion. Grey = apical foramina of the developing tooth crown. **Bottom:** Fully developed roots of different tooth types. From right to left: single rooted teeth (no IRP), two rooted teeth in which two opposing IRPs fused, and three rooted teeth in which three opposing IRPs fused.

Externally, the roots of non-human primate and fossil species appear to be variable in morphology (Table 1 & Figure 4). For example, root morphologies described as ‘plate-like’ and ‘dumb-bell’ shaped, have been described in great apes, cercopithecoids, and Plio-Pleistocene hominins (Kullmer et al., 2011; Kupczik et al., 2019; Robinson, 1956). However, until recently the full range of morphological variants has been under-explored and with contradictory or complimentary definitions (Gellis & Foley, 2021). The anthropological literature has generally emphasized root number and rare or infrequent morphologies such as Tomes’ root, Taurodont molars and/or C-shaped molars. First described in *Homo neanderthalensis* molars (Keith, 1913), mandibular post-canine tooth roots and pulp canals are sometimes C-shaped. This type of configuration consists of a root canal system in a 180-degree arc and is most common in 2nd mandibular molars (Fan, Cheung, Fan, Gutmann, & Bian, 2004; Fan, Cheung, Fan, Gutmann, & Fan, 2004; Fernandes et al., 2014). Also common in Neanderthals, taurodont molars occur when the HERS fails to invaginate at the proper horizontal level. As a result, the external shape of the root is enlarged, and the floor of the pulp chamber is displaced apically of the cemento-enamel junction. Mandibular premolars with a prominent mesial developmental groove of varying depth have been classified as Tomes’ roots (1889). This last feature is found in modern humans (Scott and Turner, 1997) and fossil members of Homo from the Chinese Middle Pleistocene and European Early Pleistocene (Prado-Simón et al., 2012; Xing et al., 2018).

**Table 1:**
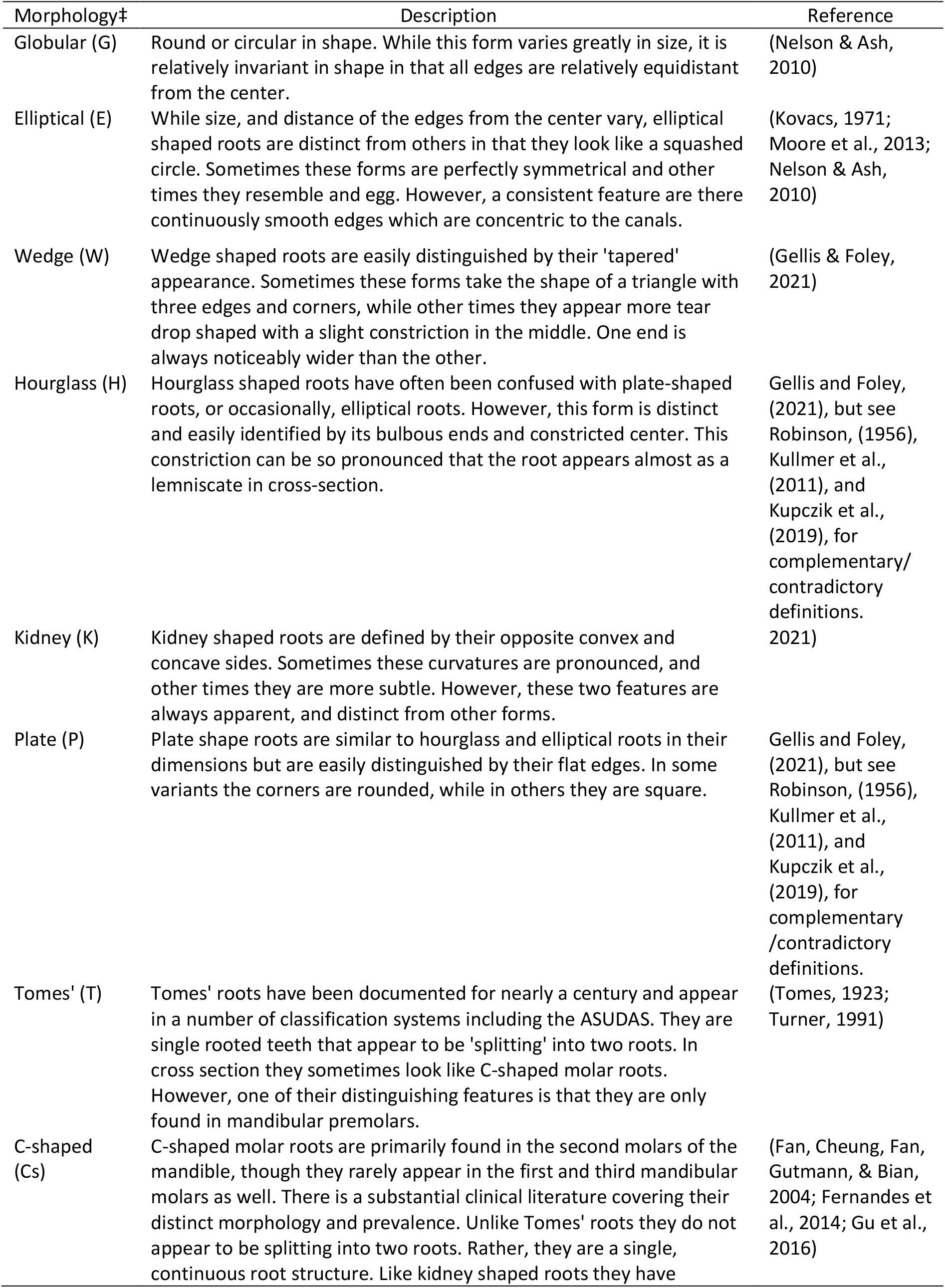
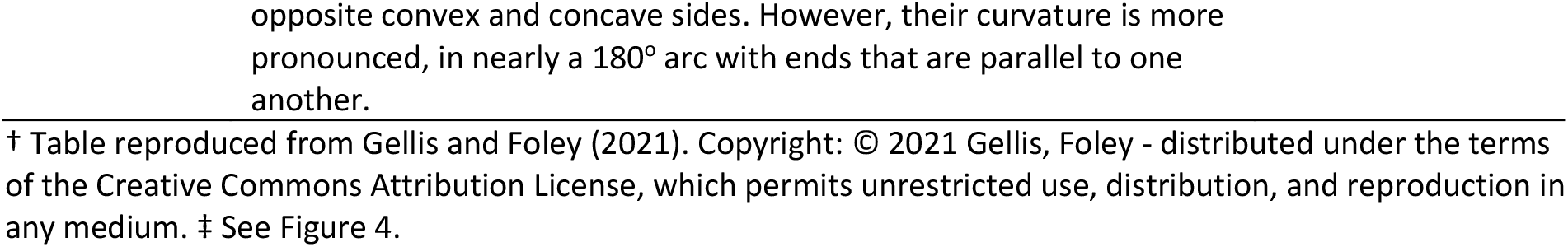
Description of external tooth root morphologies at the midpoint†

| Morphology‡ | Description | Reference |
| --- | --- | --- |
| Globular (G) | Round or circular in shape. While this form varies greatly in size, it is relatively invariant in shape in that all edges are relatively equidistant from the center. | (Nelson & Ash, 2010) |
| Elliptical (E) | While size, and distance of the edges from the center vary, elliptical shaped roots are distinct from others in that they look like a squashed circle. Sometimes these forms are perfectly symmetrical and other times they resemble an egg. However, a consistent feature are there continuously smooth edges which are concentric to the canals. | (Kovacs, 1971; Moore et al., 2013; Nelson & Ash, 2010) |
| Wedge (W) | Wedge shaped roots are easily distinguished by their 'tapered' appearance. Sometimes these forms take the shape of a triangle with three edges and corners, while other times they appear more tear drop shaped with a slight constriction in the middle. One end is always noticeably wider than the other. | (Gellis & Foley, 2021) |
| Hourglass (H) | Hourglass shaped roots have often been confused with plate-shaped roots, or occasionally, elliptical roots. However, this form is distinct and easily identified by its bulbous ends and constricted center. This constriction can be so pronounced that the root appears almost as a lemniscate in cross-section. | Gellis and Foley, (2021), but see Robinson, (1956), Kullmer et al., (2011), and Kupczik et al., (2019), for complementary/contradictory definitions. |
| Kidney (K) | Kidney shaped roots are defined by their opposite convex and concave sides. Sometimes these curvatures are pronounced, and other times they are more subtle. However, these two features are always apparent, and distinct from other forms. | 2021) |
| Plate (P) | Plate shape roots are similar to hourglass and elliptical roots in their dimensions but are easily distinguished by their flat edges. In some variants the corners are rounded, while in others they are square. | Gellis and Foley, (2021), but see Robinson, (1956), Kullmer et al., (2011), and Kupczik et al., (2019), for complementary/contradictory definitions. |
| Tomes' (T) | Tomes' roots have been documented for nearly a century and appear in a number of classification systems including the ASUDAS. They are single rooted teeth that appear to be 'splitting' into two roots. In cross section they sometimes look like C-shaped molar roots. However, one of their distinguishing features is that they are only found in mandibular premolars. | (Tomes, 1923; Turner, 1991) |
| C-shaped (Cs) | C-shaped molar roots are primarily found in the second molars of the mandible, though they rarely appear in the first and third mandibular molars as well. There is a substantial clinical literature covering their distinct morphology and prevalence. Unlike Tomes' roots they do not appear to be splitting into two roots. Rather, they are a single, continuous root structure. Like kidney shaped roots they have | (Fan, Cheung, Fan, Gutmann, & Bian, 2004; Fernandes et al., 2014; Gu et al., 2016) |

**Figure 4:**
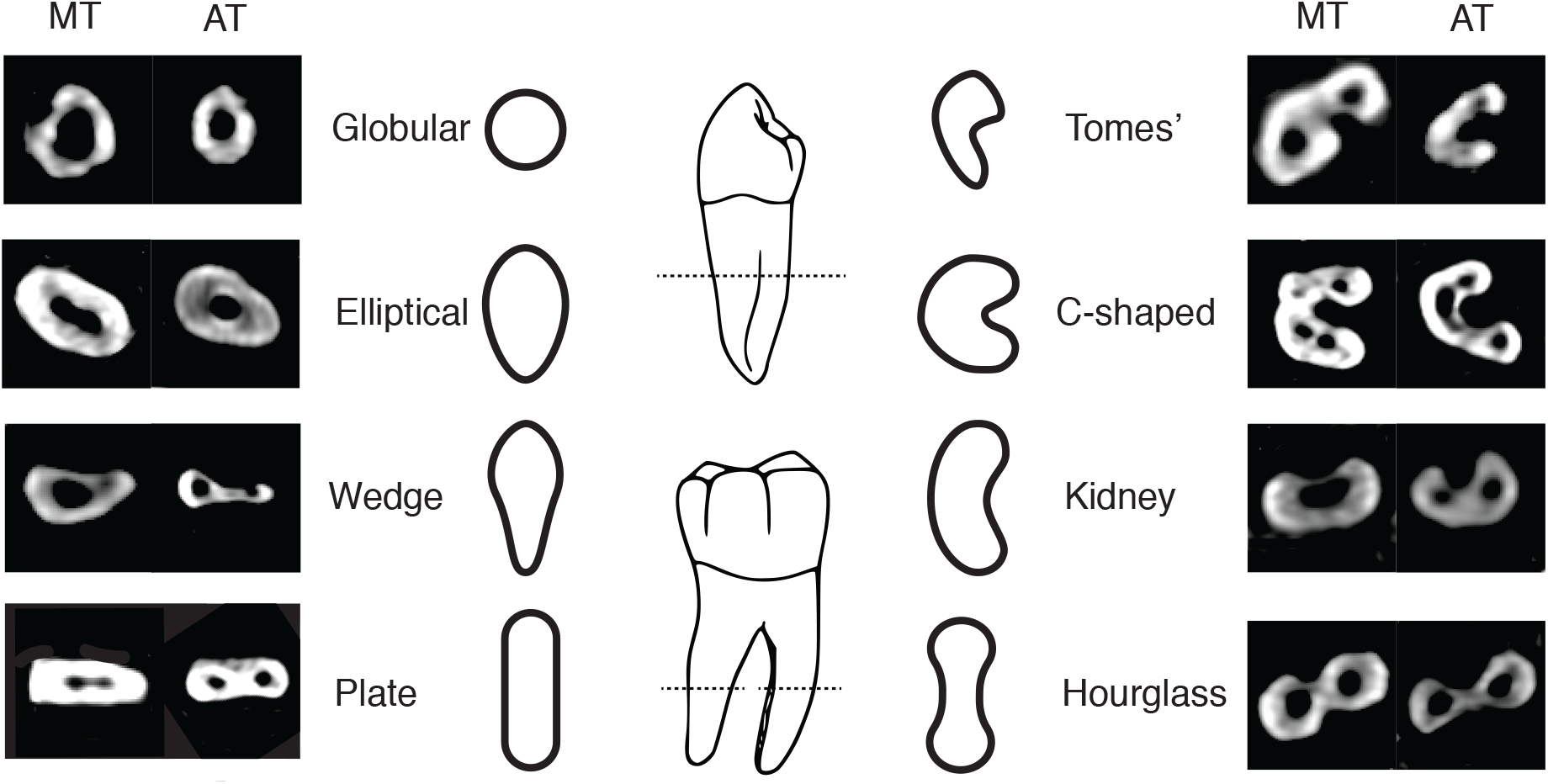
Left and right columns = axial CT slices showing external root morphologies at the middle third (MT) and apical third (AT). Centre illustrations = root morphologies at center of root/s. Figure reproduced from Gellis and Foley (2021). Copyright: © 2021 Gellis, Foley - distributed under the terms of the Creative Commons Attribution License, which permits unrestricted use, distribution, and reproduction in any medium.

Similar to the variation found in tooth cusp morphology (Turner and Nichol, 1991), these morphologies exist as distinctive anatomical variants (Figure 4). However, what defines the most frequent phenotype (MFP) - root and canal number, external root morphology and internal canal configuration - is unknown for humans. Thus, it is possible that there is an untapped wealth of useful morphological features in tooth roots.

The biological basis for tooth root variation stems from developmental changes during root growth (Shields, 2005; Wright, 2007). This variation is established early in the roots’ development and is, in part, influenced by the size of the tooth germ. A small increase in tooth germ size will lead to an expansion in cell count and reproduction in the HERS (Shields, 2005). Tooth germ size also effects molecular signaling governing the development of interradicular processes which can lead to differing degrees of fusion and bifurcation (ibid). While canal number has been shown to predict root number (Gellis & Foley, 2022), explanations for external tooth root morphology recorded in the literature (Table 1) are almost non-existent.

### Canal variation

It is easy to conceptualize canals as round holes which taper towards the roots’ apex, mirroring the external morphology of the root. In reality, the number and shape of canals does not always covary with number of roots, and many teeth have multiple canals within a single root. Canals can be round, oval shaped, or one of several isthmus configurations; and multiple canals, when found in a single root, can join and separate in unpredictable places between the cemento-enamel junction (CEJ) and root apices (Figure 5). Several clinicians have developed typologies to classify root canal variation (Weine, 1969; Vertucci and Gegauff, 1979; Hsu and Kim, 1997; Fan et al., 2004a; Vertucci, 2005). The variation between systems is in part due to technological restrictions (e.g., radiography versus CT and µCT) and the appearance of accessory and lateral canals which extend from the pulp to the periodontal tissues surrounding the teeth; as some practitioners choose to include accessory canals in their typologies, while others focus on canal structures that extend from the pulp chamber and exit foramina in the apices of the root.

**Figure 5:**
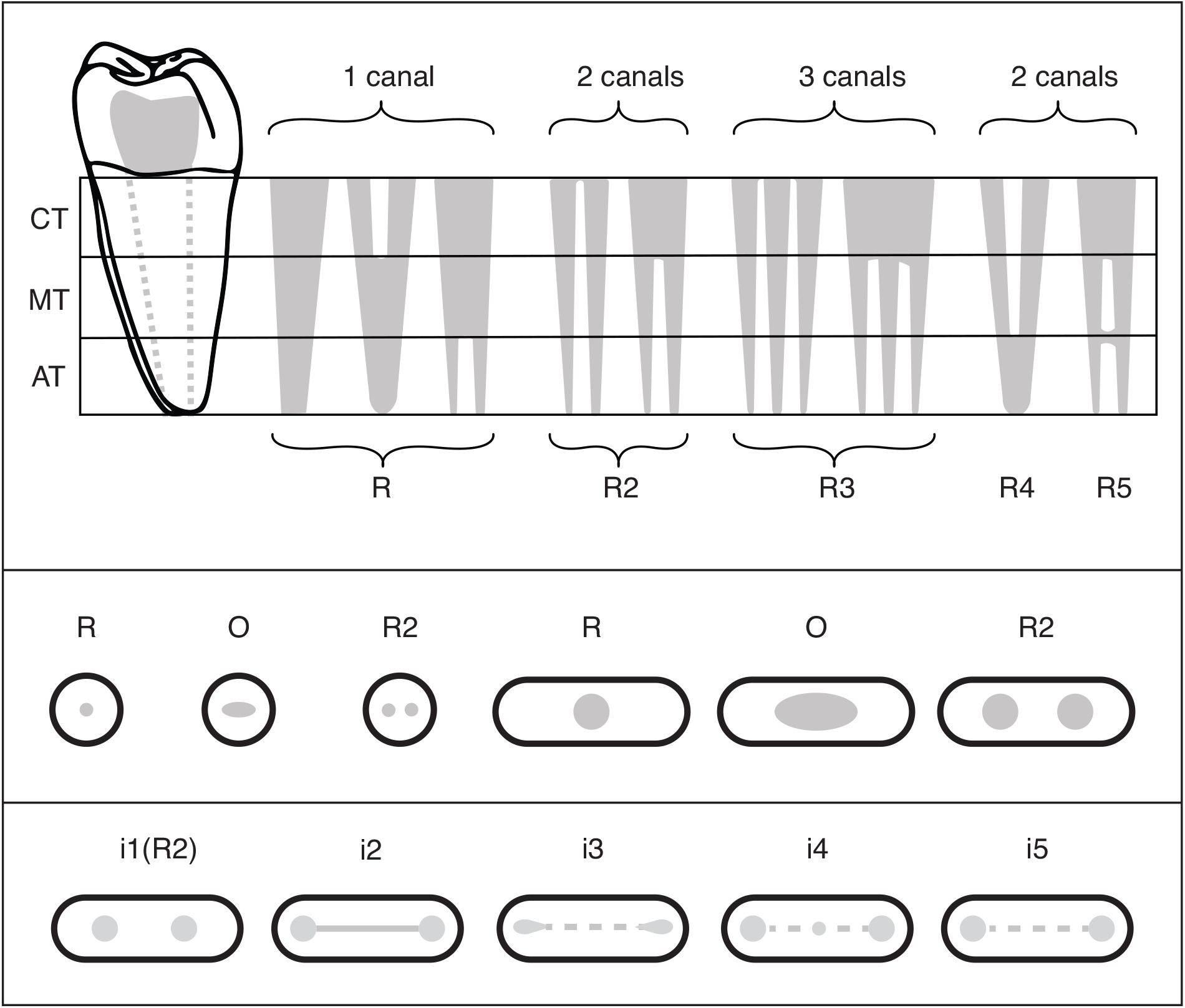
Canals (in grey) have different numbers and morphologies, and frequently join and separate. **Top**: Canal configurations (R – R5) and numbers. Canal structures that bifurcate 33% or more are considered multi-canaled. **Middle:** Root canals are almost always round (R) or oval (O). From left to right: globular (G) and Plate (P) shaped roots in cross section showing a single round (R) canal, a single oval (O) canal, and an R2 (two round canals) **Bottom:** Isthmus canals are characterized by a ribbon of tissue between two round (R) canals. CT = cervical third, MT = middle third, AT = apical third.

### Morphogenetic gradients

Teeth have been observed to exist on a gradient in which adjacent teeth are more similar to one another than non-adjacent teeth (Butler, 1937, 1939, 1963). For example, lateral incisors are more similar to central incisors than they are to canines, while molars are more similar to one another than they are to premolars. Butler (ibid) conceptualized these gradients as morphogenetic fields in which different tooth types are determined by where they develop in the jaws. For Dental Anthropology, Dahlberg (1945b) adapted and extended Butler’s (1937, 1939) morphogenetic fields of mammalian teeth from three (incisor, canine, and molar) to four fields corresponding to four morphological classes of human teeth: incisors, canines, premolars and molars. Dahlberg (1945b) assigned each field a ‘key’ tooth — the most mesial member of each field, with exception of the mandibular central incisor — which he deemed the most developmentally and evolutionary stable tooth in terms of size, numerical variation (e.g., root or cusp number), and/or morphological variation.

Several theories have been proposed to explain patterns of morphological gradients in tooth rows: including morphogenetic field theory (Butler, 1937, 1939, 1956), the clone model (Osborn, 1978), the odontogenetic homeobox code model (McCollum & Sharpe, 2001), cooperative genetic interaction (Mitsiadis & Smith, 2006), and the inhibitory cascade model (Kavanagh et al., 2007). Though consensus has not been reached, Mitsiadis and Smith’s (2006) cooperative genetic interaction model provides a synthesis of the molecular processes underlying patterned morphogenetic fields so far. Briefly, the differential expression of homeobox genes — clusters of regulatory genes that are spatially and temporally expressed during regulatory development (Gehring, 1993) — within the ectomesenchyme (Section 2.1 & Figure 2.1) leads to variation in the number, shape and size of teeth via modulation of signaling molecules. While the majority of this work has focused on development and patterning of tooth crowns, there is expected carry over to tooth roots due to shared developmental process. However, for reasons discussed below (Section 2.5), the genomic pathways of tooth root development are poorly understood.

For Dental Anthropology, Dahlberg (1945b) adapted and extended Butler’s (1937, 1939) morphogenetic fields of mammalian teeth from three (incisor, canine, and molar) to four fields corresponding to four morphological classes of human teeth: incisors, canines, premolars and molars. Dahlberg (ibid) assigned each field a ‘key’ tooth — the most mesial member of each field, with exception of the mandibular central incisor — which he deemed the most developmentally and evolutionary stable tooth in terms of size, numerical variation (e.g., root or cusp number), and/or morphological variation.

### Inter-trait association and independent observations

The inter-relationships of dental traits have been well studied in clinical and anthropological contexts. Tooth dimensions are strongly correlated with one another (Stanley M. Garn et al., 1965, 1968; Harris & Lease, 2005), as are eruption sequences (Ash, 2013; Fleagle, 2013; B. H. Smith, 1991), timing of mineralization (Miller, 2013; Nelson & Ash, 2010; Reid et al., 1998), and agenesis (S M. Garn et al., 1963; Nieminen, 2009). However, non-metric crown and root traits are poorly correlated and usually expressed independently of one another (Corruccini, 1976; Markowski, 1995; Scott et al., 2018). The working assumption is that non-metric crown and root traits are expressed independent of one another, and show little or no interaction with metric crown and root dimensions, or tooth anagenesis (Scott & Irish, 2017).

An exception to the above, nonmetric traits expressed on members of the same morphogenetic field sometimes do show significant correlations (Scott, 1977; Scott & Irish, 2017). While a ‘key tooth’ (i.e., the most mesial) might be the most developmentally and evolutionary stable tooth in a morphogenetic field, it does not always exhibit the full frequency or degree of expression of traits within that field (Butler, 1963). For example, five-cusped M_1_s appear in 85 -100% of all human populations, while M_2_s have a relatively higher frequency of four-cusped molars (Hemphill, 2002; Scott & Turner, 1997). Other traits, such as the protostylid appear with greatest frequency in M_1_s, but with a higher degree of expression in M_2_s and M_3_s (Scott & Turner, 1997).

## Materials and Methods

### Dental formula

Premolars are indicated with P, and molars with M and super- and subscripts differentiate the teeth of the maxilla and mandible respectively. For example, M^2^ indicates the 2nd maxillary molar while M_1_ indicates the 1st mandibular molar. Over the course of evolution of their evolutionary history, apes and old world monkeys have lost the first and second premolars of their evolutionary ancestors (Novacek, 1986; White et al., 2012). Thus, the remaining two premolars are numbered 3 and 4.

### Human samples

Data used in this study were originally collected from CT scans of 945 individuals (Table 2) stored in osteological collections at the Smithsonian National Museum of Natural History, Washington D.C., USA, American Museum of Natural History, New York, USA, and the Duckworth Laboratory at the University of Cambridge, England (Gellis & Foley, 2021; Gellis, 2021). Only adult individuals, based on the eruption, occlusion, and closed root apices of M^3^s/M_3_s (or M^2^s/M_2_s in the case of congenitally absent M^3^s/M_3_s), were used in this study.

**Table 2:**
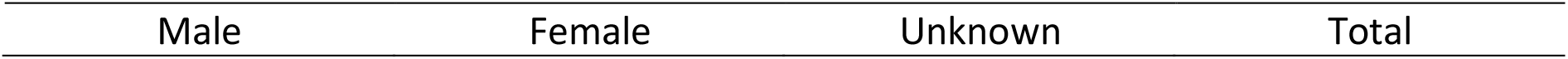
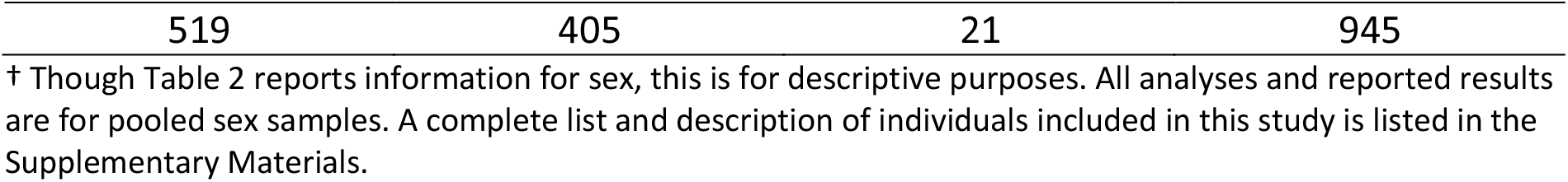
Counts for individuals used in this study†

### Canal count, configuration, and external root morphology

Canal count, configuration, and external root morphology for 5,790 teeth (Table 3) collected from a global sample of 945 individuals were downloaded from the Tooth Root Phenotypic Dataset (Gellis & Foley, 2021; Gellis, 2021) and analyzed with the R Project for Statistical Computing, version 3.6.3 (R Core Team, 2017). Only roots of from the right sides of the maxillary and mandibular dental arcades were analyzed to avoid issues with asymmetry and artificially inflated sample size. All analyses and reported results are for pooled sex samples.

**Table 3:** Tooth, canal, and root counts used in this study†

| Tooth | tooth(n) | root(n) | canals(n) | Tooth | tooth(n) | root(n) | canals(n) |
| --- | --- | --- | --- | --- | --- | --- | --- |
| Maxilla |  |  |  | Mandible |  |  |  |
| P<sup>3</sup> | 515 | 739 | 959 | P<sub>3</sub> | 343 | 345 | 433 |
| P<sup>4</sup> | 467 | 530 | 708 | P<sub>4</sub> | 313 | 313 | 326 |
| M<sup>1</sup> | 697 | 2,060 | 2,430 | M<sub>1</sub> | 410 | 894 | 1,421 |
| M<sup>2</sup> | 596 | 1,561 | 1,910 | M<sub>2</sub> | 385 | 727 | 1,093 |
| M<sup>3</sup> | 362 | 831 | 1,065 | M<sub>3</sub> | 278 | 563 | 747 |
| Total | 2,637 | 5,721 | 7,072 | --- | 1,729 | 2,842 | 4,020 |

† Data from the Tooth Root Phenotypic Dataset (Gellis & Foley, 2021; Gellis, 2021)

### Canal number and configuration

Canal numbers and configurations from the Tooth Root Phenotypic Dataset (Gellis, 2021) are, in part, derived from the Turner Index (Turner, 1981) and congruent with Vertucci’s canal configurations (F. Vertucci et al., 1974; F. J. Vertucci & Gegauff, 1979). Briefly, canals individual canals take the shape of round (R) or oval (O). A single canal extends from the pulp cavity and exits a foramen in the apical third of the root. In the case when two canals bifurcate from a single canal, if the degree of bifurcation is >= 33% of the total length of the canal structure, the canal count is two. The full method and its application are discussed in Gellis and Foley (2021, 2022), but are visualized in Figure 5.

### Inter-trait association and independent observation

For basic descriptives, results include counts of roots, canals, and external root morphologies for all post-canine teeth from the right maxillary and mandibular dental arcades combined by tooth type (premolars or molar). However, to avoid violations of statistical independence of variables, primary statistical analyses were carried out only for ‘key teeth’ (i.e., P^3^, M^1^, P_3_, and M_1_) as discussed in Scott and Turner II (2015).

### Statistical analysis

Multinomial logistic regression (MLR) was used to test and predict relationships between root morphology, and canal configuration and number. MLR uses a logarithmic function to reduce probability values between 0 and 1, with 0 indicating 0% predictive value, and 1 indicating 100% predictive value. Results report probability values between .50 and 1. Multinomial logistic regression was carried out using the “nnet” package (Venables and Ripley, 2002). Because C-shaped (Cs) molars appear primarily in M_2_s, a secondary analysis was carried out as M_2_ should be considered the key tooth for this trait (Turner II et al., 1991).

## Results

Post-canine teeth (Table 3) of 945 individuals from a global sample (Table 2) were used to test if canal number and configuration predicted external root morphology. Identification of the most frequent phenotype (MFP) of root morphology, canal count, and canal configuration are required for multinomial logistic regression. Counts of canal shapes and configurations in single, double, and triple canaled roots were plotted to discern the MFP for descriptive purposes (Figures 6 - 15).

**Figure 6:**
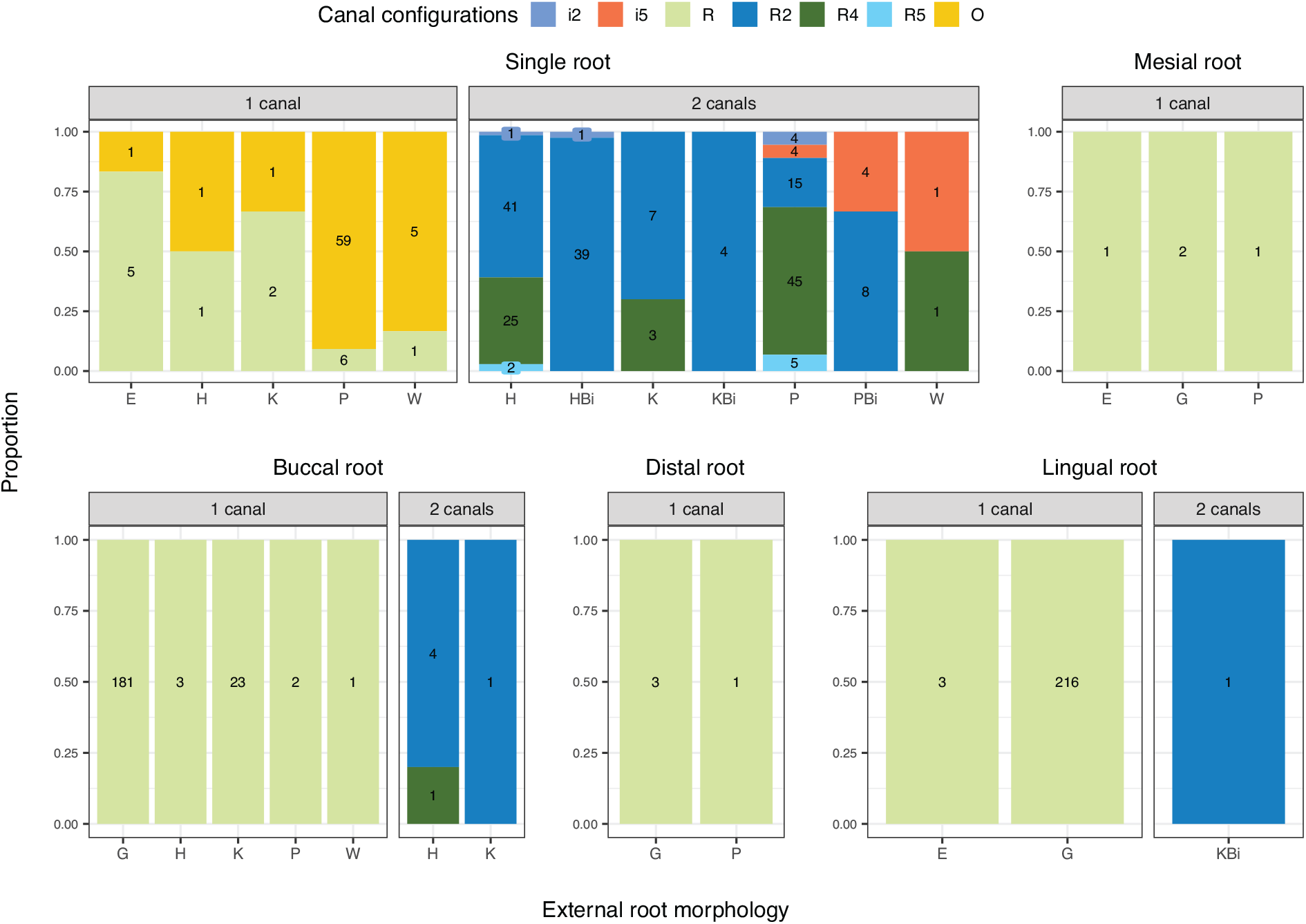
Proportion plots of canal shapes and configurations in roots of P^3^s by canal number. Counts of roots by external morphology inset. Root forms: E = elliptical, G= globular, H = hourglass, K = kidney, P = plate, W = wedge. Bi = root form with apical bifurcation. Counts are calculated over combined root morphologies containing 1 or 2 canals.

### Maxillary Premolars

The MFP of single rooted, single canaled P^3^s is plate shaped (P) with an oval (O) shaped canal (n = 59), while the double canaled MFP is also plate shaped but with an R4 canal configuration (n= 45)(Figure 6). Single rooted premolars also show the most variation in canal configurations and external root morphologies, with multiple forms appearing. Mesial roots appear in low numbers in this study and have no double canaled variants. The MFP of single canaled mesial roots is globular (G) shaped with a round (R) canal (n = 2). The MFP of P^3^ buccal roots is globular with a round canal (n = 181), while the MFP of the double canaled variant is hourglass (H) with and R2 canal configuration (n = 4). Globular shaped roots with a single round canal are the MFP of P^3^ lingual roots (n = 216).

Of the maxillary molars, single rooted P^4^s have the most variation in external morphologies and internal canal configurations(Figure 7). Like P^3^S, the MFP of single canaled P^4^s are plate shaped roots with oval canals (n = 170). The MFP of double canaled variant is also plate shaped, but with an R4 canal configuration (n = 48). The MFP of P^4^ buccal, distal, and lingual roots is globular with a single round canal. This form, and a fused, bifurcated, mesio-lingual (MLFBi) variants are equally represented in P^4^ mesial roots.

**Figure 7:**
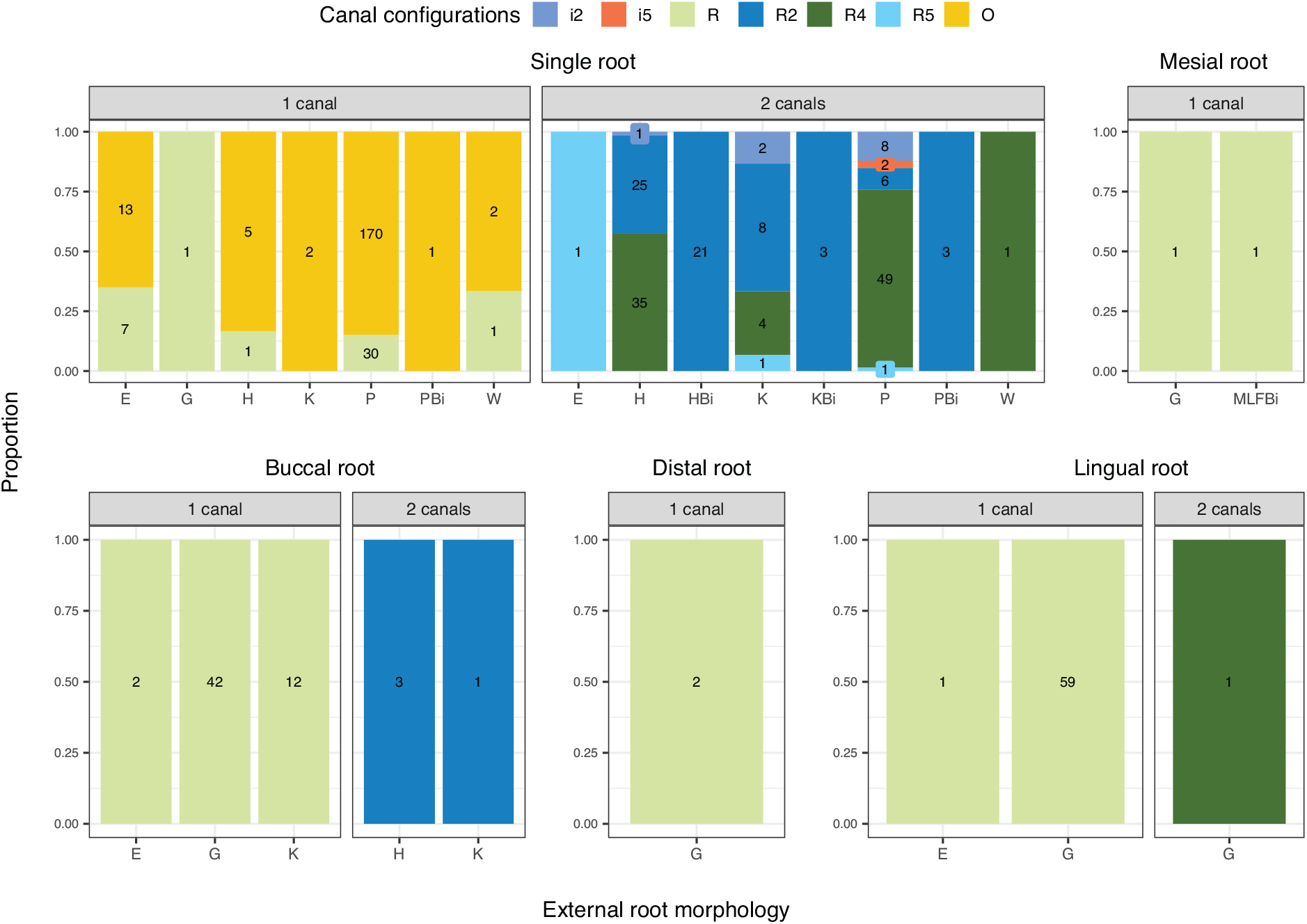
Proportion plots of canal shapes and configurations in roots of P^4^s by canal number. Counts of roots by external morphology inset. Root forms: E = elliptical, G= globular, H = hourglass, K = kidney, P = plate, W = wedge. Bi = root form with apical bifurcation. F = indicates fused roots (e.g., MLF = mesio-lingual fused). Counts are calculated over combined root morphologies containing 1 or 2 canals.

### Maxillary molars

There is a wide range of variation in M^1^ root morphologies and configurations (Figure 8). The MFP of single rooted M^1^s is plate shaped with an R2 canal configuration (n = 2). No single canaled M^1^ variant is present in the sample. M^1^ mesial roots are the most diverse in external morphologies and internal configurations. Wedge (W) shaped mesial roots with a round canal are the single canaled MFP (n = 125), while wedge shaped with an R2 configuration is the MFP of the double canaled variant. There is no MFP for M^1^ buccal roots. Elliptical (E) shaped roots with a single round canal are the MFP of distal (n = 230) and lingual (n = 206) maxillary molar roots.

**Figure 8:**
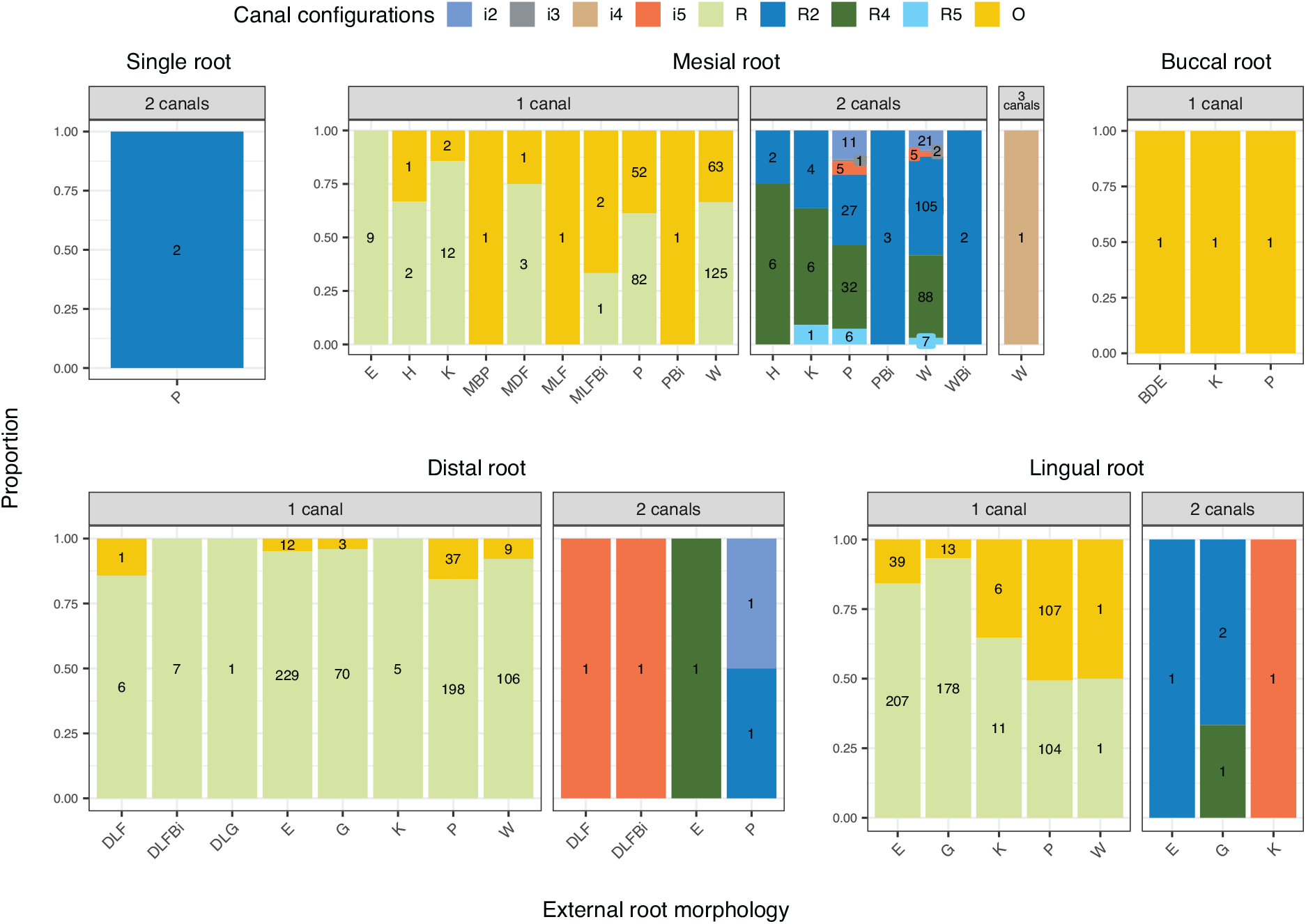
Proportion plots of canal shapes and configurations in roots of M^1^s by canal number. Counts of roots by external morphology inset. Root forms: E = elliptical, G= globular, H = hourglass, K = kidney, P = plate, W = wedge. Bi = root form with apical bifurcation. M = mesial, B = buccal, D = distal, L = lingual. MB = mesio-buccal, ML = mesio-lingual, etc. When appended with an F = indicates fused roots (e.g., MLF = mesio-lingual fused). Counts are calculated over combined root morphologies containing 1, 2, or 3 canals.

**Figure 9:**
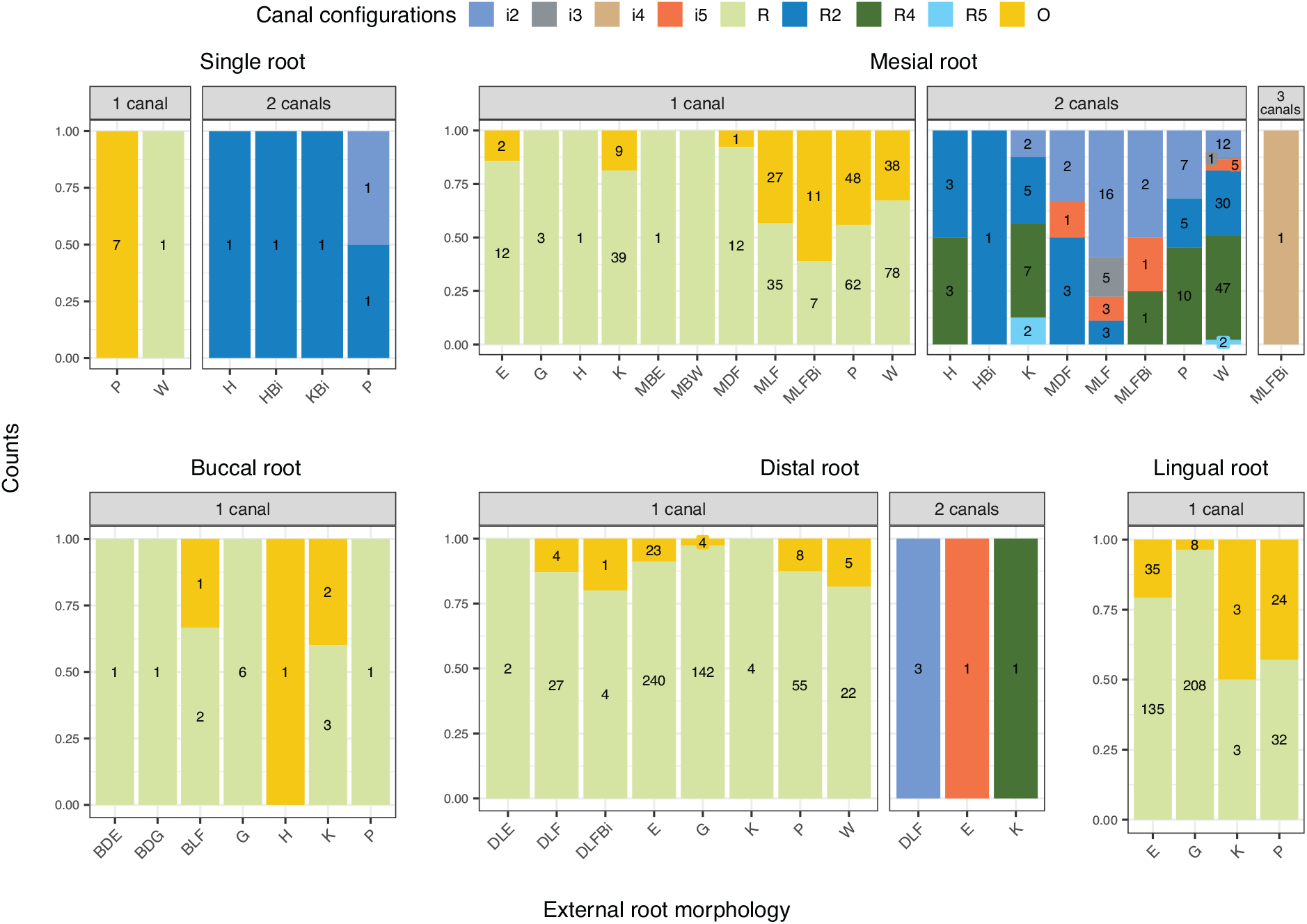
Proportion plots of canal shapes and configurations in roots of M^2^s by canal number. Counts of roots by external morphology inset. Root forms: E = elliptical, G= globular, H = hourglass, K = kidney, P = plate, W = wedge. Bi = root form with apical bifurcation. M = mesial, B = buccal, D = distal, L = lingual. MB = mesio-buccal, ML = mesio-lingual, etc. When appended with an F = indicates fused roots (e.g., MLF = mesio-lingual fused). Counts are calculated over combined root morphologies containing 1 or 2 canals.

Like M^1^s, the mesial root of M^2^s displays a diverse number of external morphologies and internal configurations. Wedge shaped mesial roots with a round canal are the MFP (n = 78) for single canaled variants; wedge shaped with an R4 configuration (n = 47) are the MFP for double canaled variants. The MFP of M^1^ buccal roots is globular with a single round canal (n = 6); no double canaled buccal root variants appear in this sample. An elliptical root with a single round canal is the MFP for M^1^ distal (n = 240) and lingual (n = 208) roots.

The MFP of single rooted M^3^s is plate shaped with a single oval canal for single canaled variants (n = 19) and plate shaped with an R2 configuration (n = 4) for double canaled variants (Figure 10). Like M^1^s and M^2^s, the MFP of M^3^ mesial roots is wedge shaped with a single round canal (n = 58). The MFP of the mesial root double canaled variant is also wedge shaped, but with an R2 canal configuration. The MFP of distal (n = 161) and lingual (n= 150) roots is a globular shaped root with a single round canal. There is no MFP for buccal roots.

**Figure 10:**
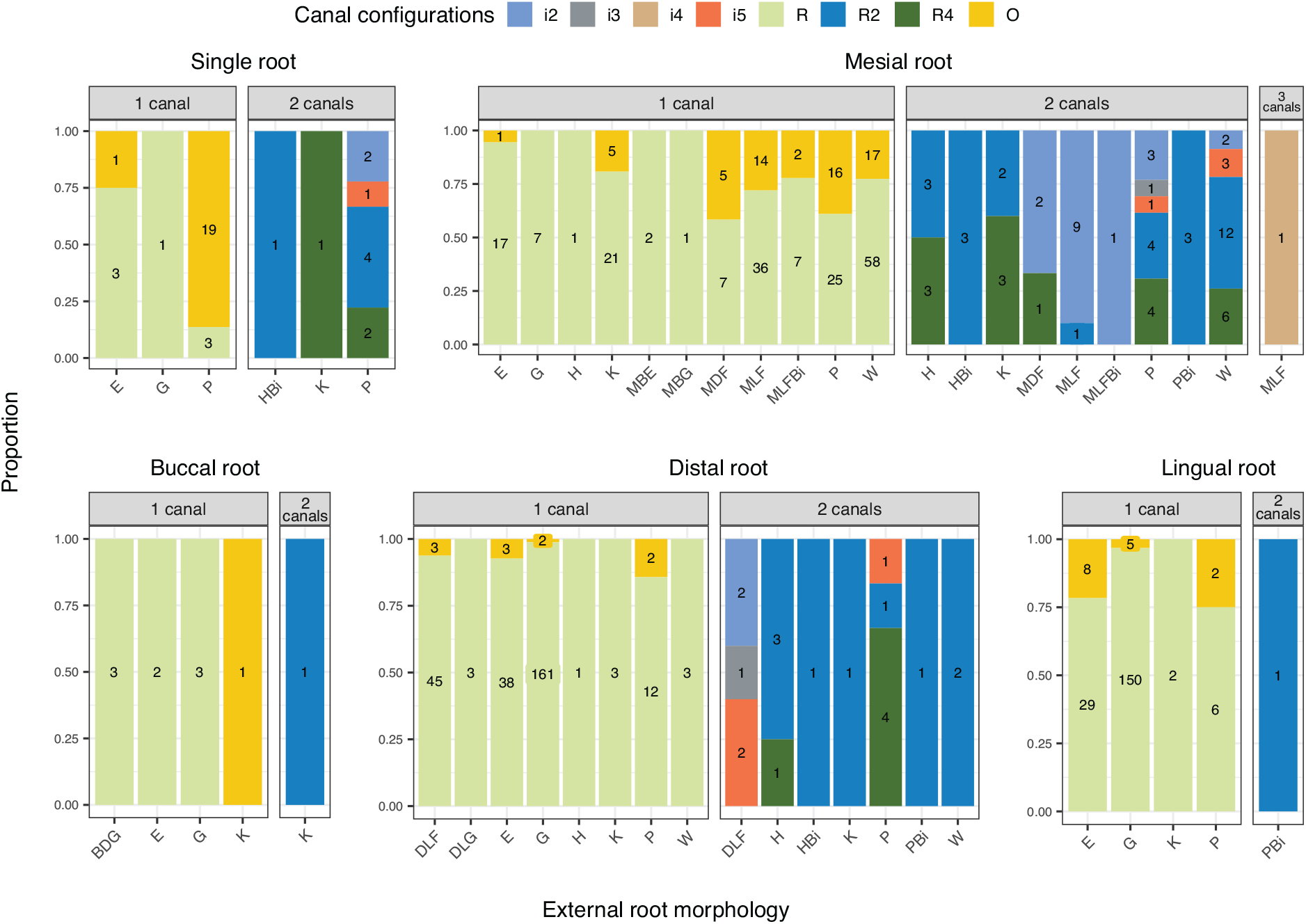
Proportion plots of canal shapes and configurations in roots of M^3^s by canal number. Counts of roots by external morphology inset. Root forms: E = elliptical, G= globular, H = hourglass, K = kidney, P = plate, W = wedge. Bi = root form with apical bifurcation. M = mesial, B = buccal, D = distal, L = lingual. MB = mesio-buccal, ML = mesio-lingual, etc. When appended with an F = indicates fused roots (e.g., MLF = mesio-lingual fused). Counts are calculated over combined root morphologies containing 1 or 2 canals.

### Mandibular Premolars

The MFP of single canaled P_3_s is plate-shaped with an oval canal (n = 110), and Tomes’ (T) root with an i5 isthmus canal configuration for double canaled variants (Figure 11). Double canaled variants mainly contain isthmus canal (i2 – i5) configurations. P_3_s with buccal and lingual roots are minimally represented in this sample.

**Figure 11:**
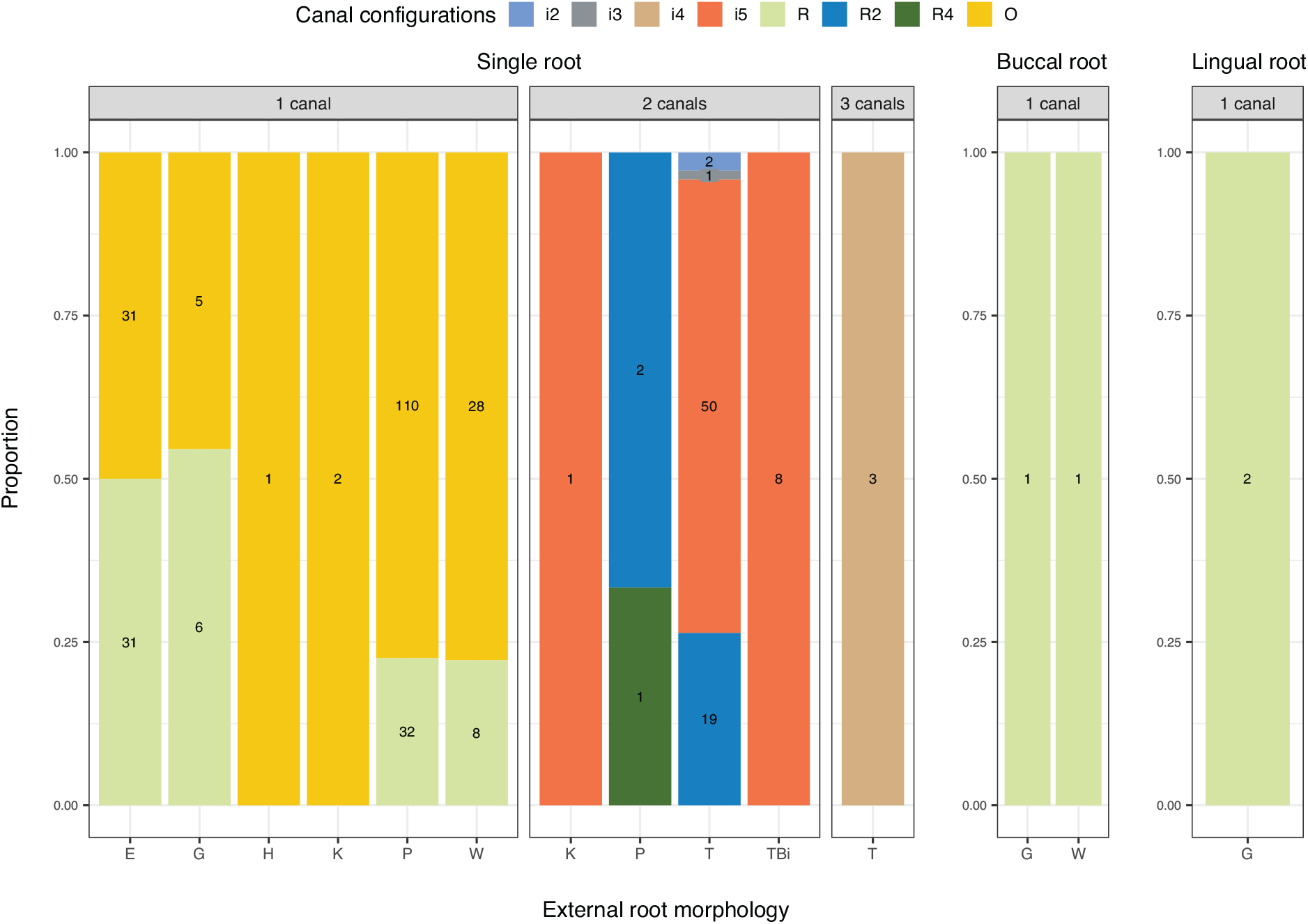
Proportion plots of canal shapes and configurations in roots of P_3_s by canal number. Counts of roots by external morphology inset. Root forms: E = elliptical, G= globular, H = hourglass, K = kidney, P = plate, T + Tomes, W = wedge. Bi = root form with apical bifurcation. M = mesial, B = buccal, D = distal, L = lingual. MB = mesio-buccal, ML = mesio-lingual, etc. When appended with an F = indicates fused roots (e.g., MLF = mesio-lingual fused). Counts are calculated over combined root morphologies containing 1, 2, or 3 canals.

Only single-rooted P_4_s are present in the sample used for this study (Figure 12). The MFP of P_4_ is a single plate shaped root with an oval canal (n = 116). Like P_3_s, Tomes’ root is also present in P_4_s. Like P_3_s, doble canaled P_4_s are primarily represented by Tomes’ rot with an isthmus canal configuration.

**Figure 12:**
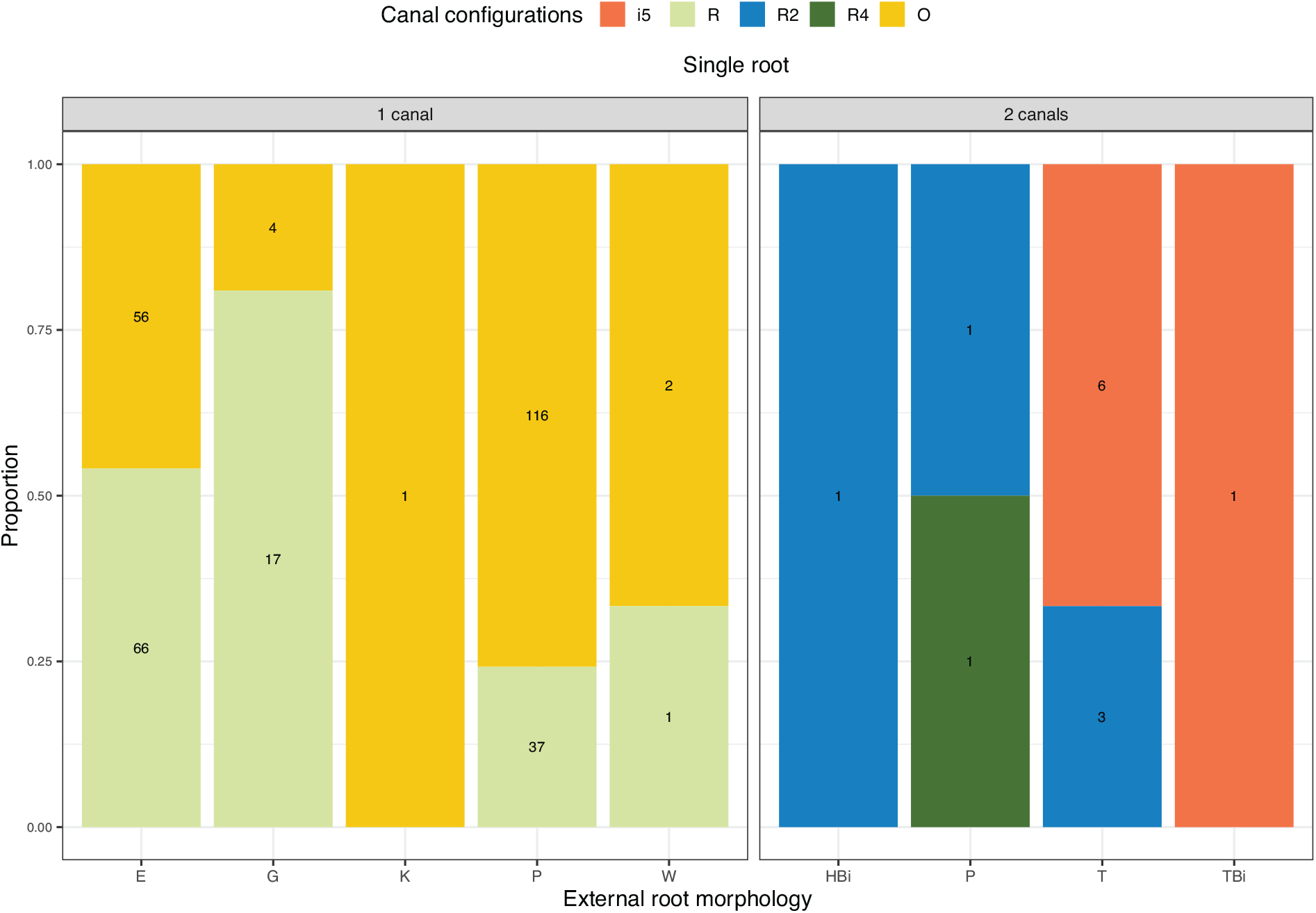
Proportion plots of canal shapes and configurations in roots of P_4_s by canal number. Counts of roots by external morphology inset. Root forms: E = elliptical, G= globular, H = hourglass, K = kidney, P = plate, T = Tomes, W = wedge. Bi = root form with apical bifurcation. M = mesial, B = buccal, D = distal, L = lingual. MB = mesio-buccal, ML = mesio-lingual, etc. When appended with an F = indicates fused roots (e.g., MLF = mesio-lingual fused). Counts are calculated over combined root morphologies containing 1 or 2 canals.

**Figure 13:**
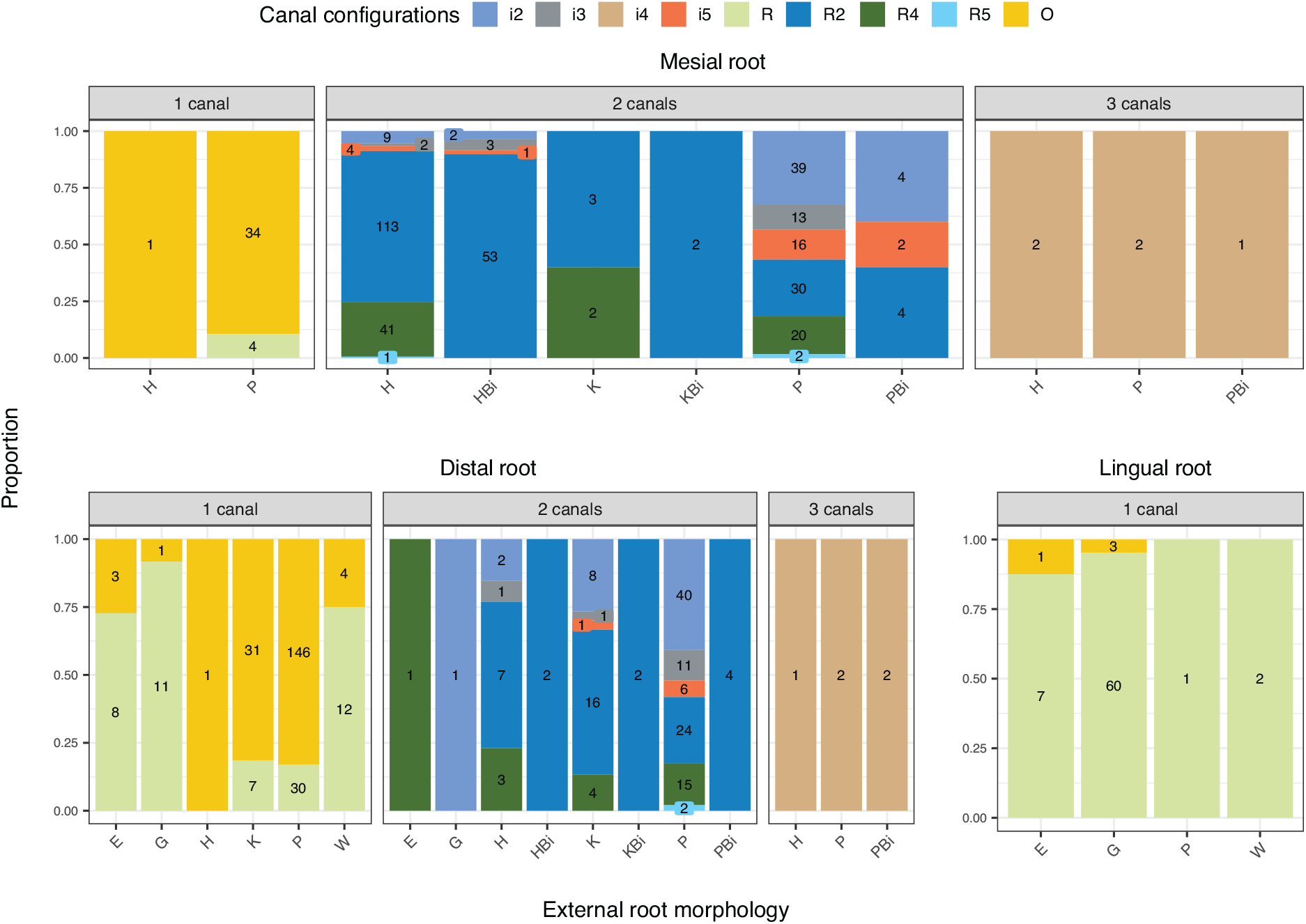
Proportion plots of canal shapes and configurations in roots of M_1_s by canal number. Counts of roots by external morphology inset. Root forms: E = elliptical, G= globular, H = hourglass, K = kidney, P = plate, W = wedge. Bi = root form with apical bifurcation. Counts are calculated over combined root morphologies containing 1, 2, or 3 canals.

### Mandibular Molars

Similar to their maxillary counterparts, mandibular molars have a wide diversity of external morphologies and internal canal configurations. No single rooted M_1_s, or M_1_s with buccal roots appear in this sample. The MFP of single canaled M_1_ mesial roots is plate shaped with an oval canal (n = 34); the MFP of double canaled variants is hourglass shaped with a double-canaled R2 configuration (n = 113). The MFP of single canaled M_1_ distal mandibular molar roots is plate-shaped with an oval canal (n = 146), and plate shaped with an R2 canal configuration (n = 24) for double canaled variants. Globular roots with a single round canal (n = 60) are the MFP for lingual roots. Three canaled variants all contain an i4 isthmus canal configuration.

The MFP of single rooted M_2_s is C-shaped (Cs) with an i3 isthmus canal (Figure 14). The MFP of single canaled M_2_ mesial roots is plate shaped with an oval canal (n = 44), while the MFP for double canaled variants is hourglass with an R2 canal configuration (n = 60). For single canaled M_2_s, the MFP of the distal roots is kidney (K) shaped with an oval canal; double canaled variants are plate shaped with an R2 canal configuration (n = 10). There is no MFP for M_2_ buccal roots.

**Figure 14:**
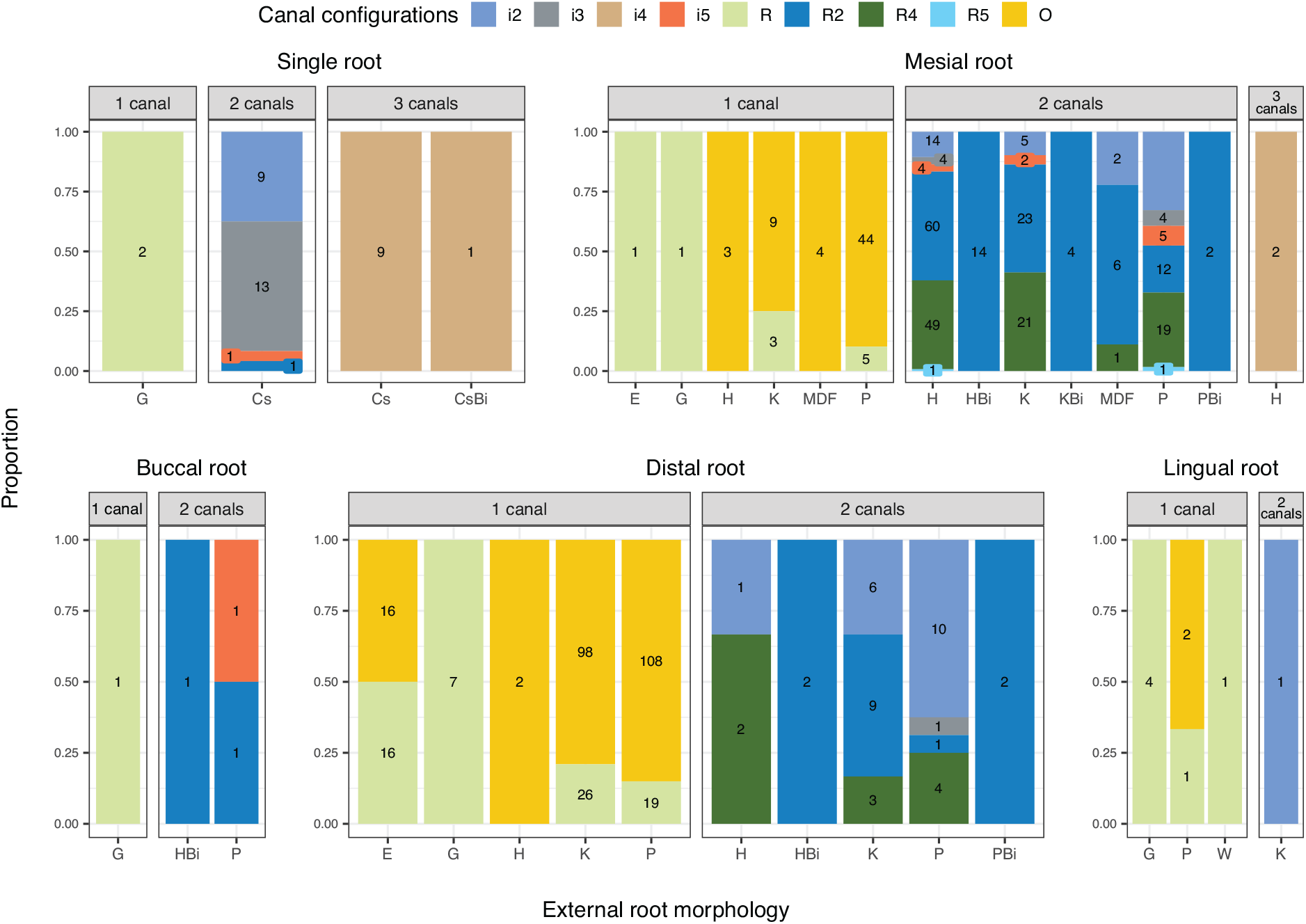
Proportion plots of canal shapes and configurations in roots of M_2_s by canal number. Counts of roots by external morphology inset. Root forms: Cs = C-shaped, E = elliptical, G= globular, H = hourglass, K = kidney, P = plate, W = wedge. Bi = root form with apical bifurcation. F = indicates fused roots (e.g., MLF = mesio-lingual fused). Counts are calculated over combined root morphologies containing 1, 2, or 3 canals.

Though minimally represented in this sample, the MFP of single rooted M_3_s is globular with single round canal for single canaled variants (n = 2) ; and C-shaped with an isthmus i5 configuration (n = 3) for double canaled variants (Figure 15). The MFP of single canaled mesial roots is plate shaped with and oval canal (n = 46). The MFP of the two canaled variant is kidney shaped with an R4 canal configuration. Buccal roots are minimally represented in the sample. The MFP of the single canaled variant is globular with a round canal (n = 9). The MFP of single canaled distal roots is plate shaped with and oval canal (n = 59). =The MFP of the single canaled lingual roots is globular with a round canal (n = 17).

**Figure 15:**
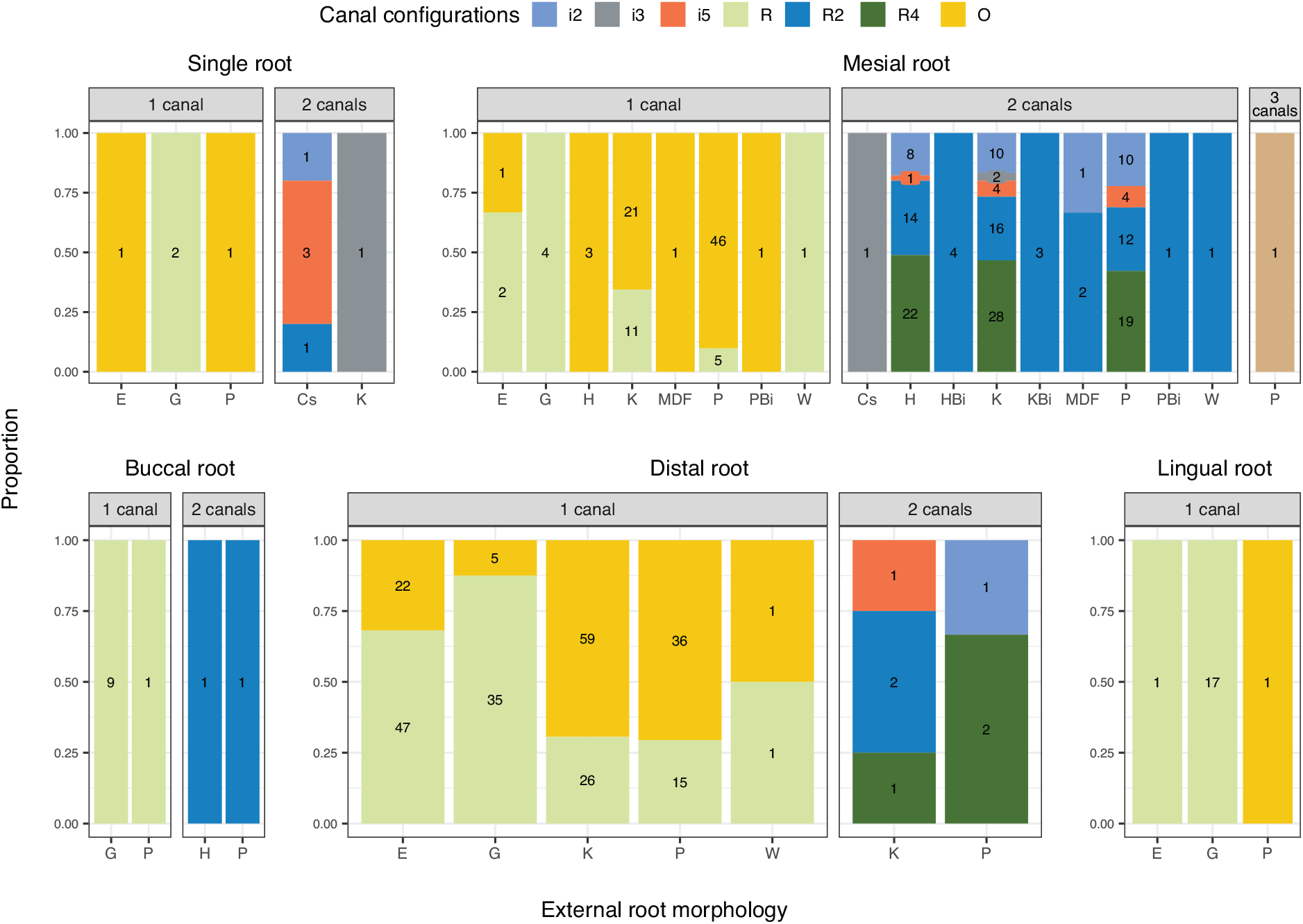
Proportion plots of canal shapes and configurations in roots of M_3_s by canal number. Counts of roots by external morphology inset. Root forms: E = elliptical, G= globular, H = hourglass, K = kidney, P = plate, W = wedge. Bi = root form with apical bifurcation. F = indicates fused roots (e.g., MLF = mesio-lingual fused). Counts are calculated over combined root morphologies containing 1 or 2 canals.

### The most frequent phenotype

Figure 16 visualizes the MFPs of external root morphology and internal canal morphology, count, and configuration in the roots of the maxilla, mandible, and the jaws combined. For some roots there is no complete (i.e., root and canal) MFP found in the sample multiple internal or external forms appear in equal proportions in the sample.

**Figure 16:**
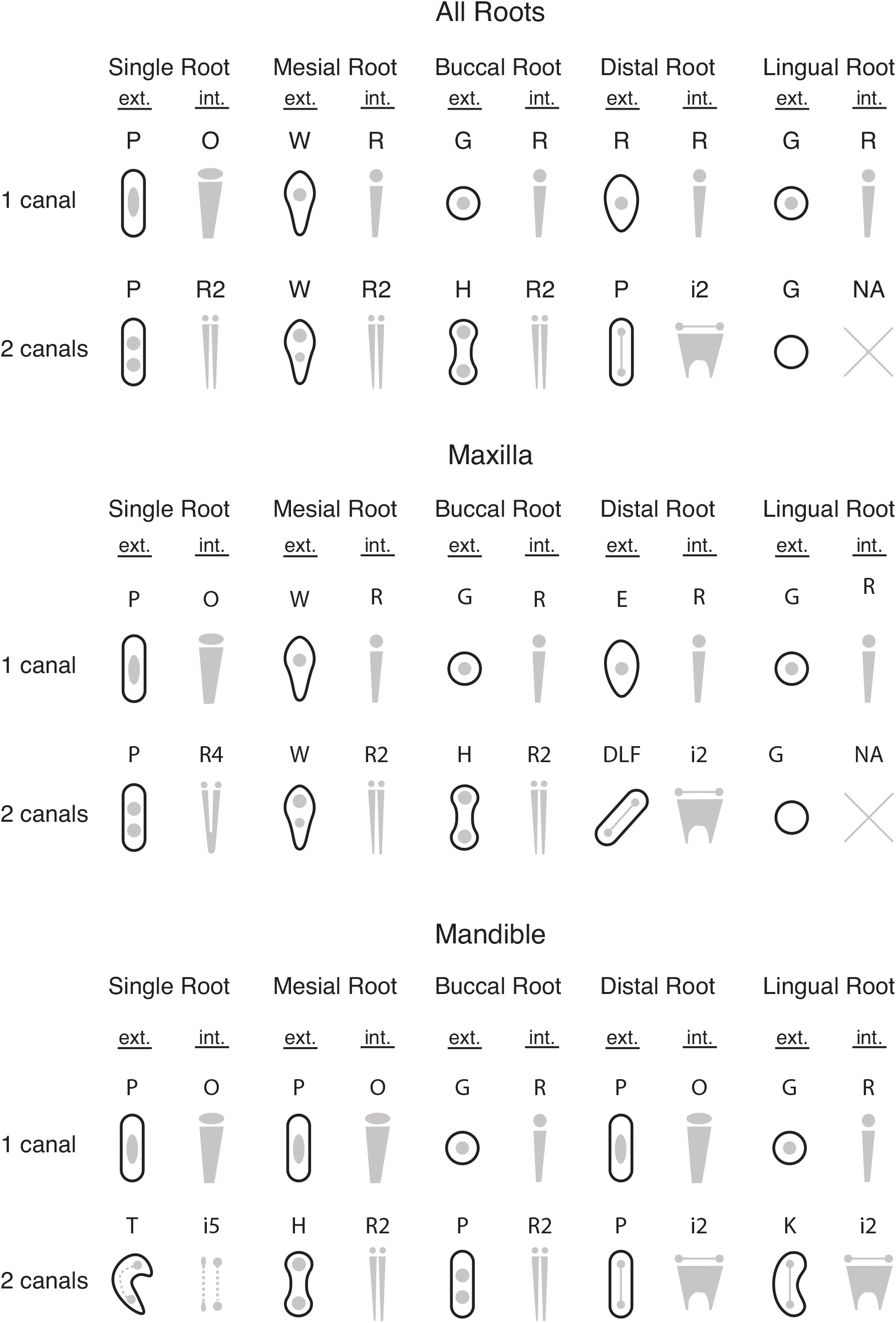
Most frequent phenotype of external root morphology and internal canal count and configuration. Solid grey = canal, ext. = external form in cross section, int. = internal configuration, X = no MFP, as all roots show different external and internal morphologies. Due to low counts, MFPs with 3 canals are not included. Canal forms and descriptions are illustrated in Figure 5. Top: Pooled MFPs from all post-canine teeth of the maxilla and mandible by root. Middle: Pooled MFPs from all post-canine teeth of the maxilla. Bottom: Pooled MFPs from all post-canine teeth of the mandible.

## Predictions of External Root Morphology

### Maxillary premolars and molars

Multinomial logistic regression (MLR) was used to test if canal number and configuration can predict external morphology of P^3^s (Figure 17). Sample size of maxillary P^3^s with mesial and distal roots were too small, and/or with too few levels for inclusion with analysis and are not included. Regardless of canal number, single rooted maxillary premolars are plate shaped. A single oval canal predicts a plate-shaped root (88.06%), and the same morphology is predicted by a double-canaled i2 configuration (66.67%), double-canaled R4 configuration (60.81%), and double-canaled R5 (71.43%) configuration. A single round canal predicts a globular buccal root (86.19%), a double-canaled R2 configuration predicts an hourglass morphology (80.00%), while a double-canaled R4 configuration predicts an hourglass morphology (100.00%). A single round canal predicts globular lingual roots (98.63%). While elliptical and bifurcated kidney root morphologies appear in P^3^ lingual roots in this sample (Figure 6), their numbers are too low (< 4) compared to the globular morphology (n= 216), and thus they are not included in this analysis.

**Figure 17:**
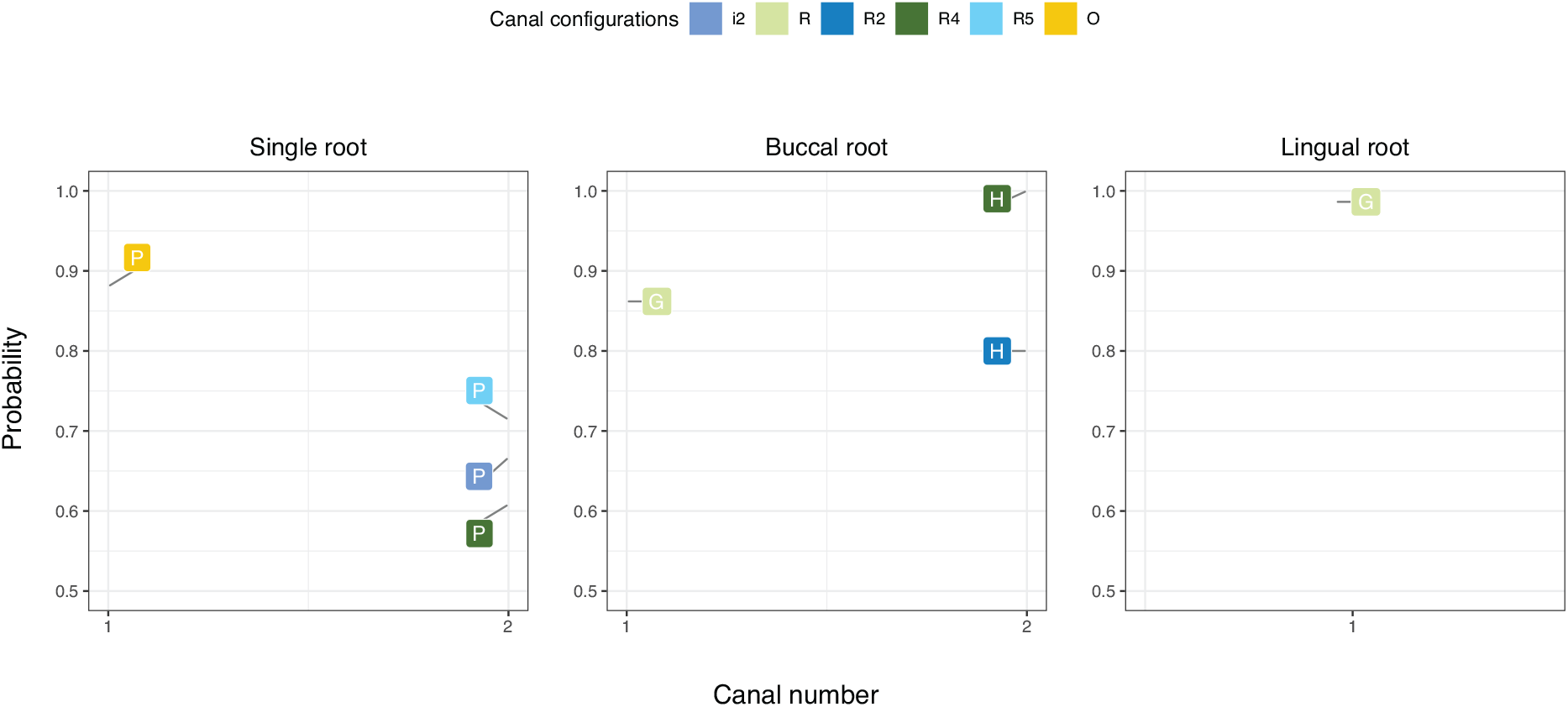
MLR of canal number to canal configuration and root morphology in single, buccal, and lingual roots of P^3^s. Most frequent phenotype for single roots = plate-shaped (P); buccal roots = globular (G); lingual roots = globular (G).

Results of MLR for M^1^s are presented in Figure 18. Sample size of single-rooted M^1^s (n = 2) and M^1^s (n=4) with buccal roots were too small and with too few levels for inclusion in analysis. While the MFP of mesial roots is wedge shaped, there are no single or double-canaled canal configurations that predict this shape with over 73.43% accuracy. The three-canaled i4 configuration predicts wedge shaped mesial roots with 100.00% accuracy. The single-canaled form is predicted by an oval canal (50.81%). A number of isthmus canal variations appear, though it is the double-canaled R2 configuration that has the most predictive power (73.43%). Isthmus canal i2 and R2 canal configurations have the most predictive power for double-canaled plate-shaped distal roots (100.00%); while the double-canaled R4 configuration predicts elliptical shaped roots with 100.00% accuracy. The i5 isthmus canal predicts fused disto-lingual roots (DLFBi) with 50.00% accuracy. Single-canaled plate-shaped lingual roots are predicted by oval canals (59.68%). Double-canaled kidney shaped lingual roots are predicted by i5 isthmus canal configurations (100.00%), while globular variants are predicted by R2 (66.67%) and R4 (100.00%) canal configurations.

**Figure 18:**
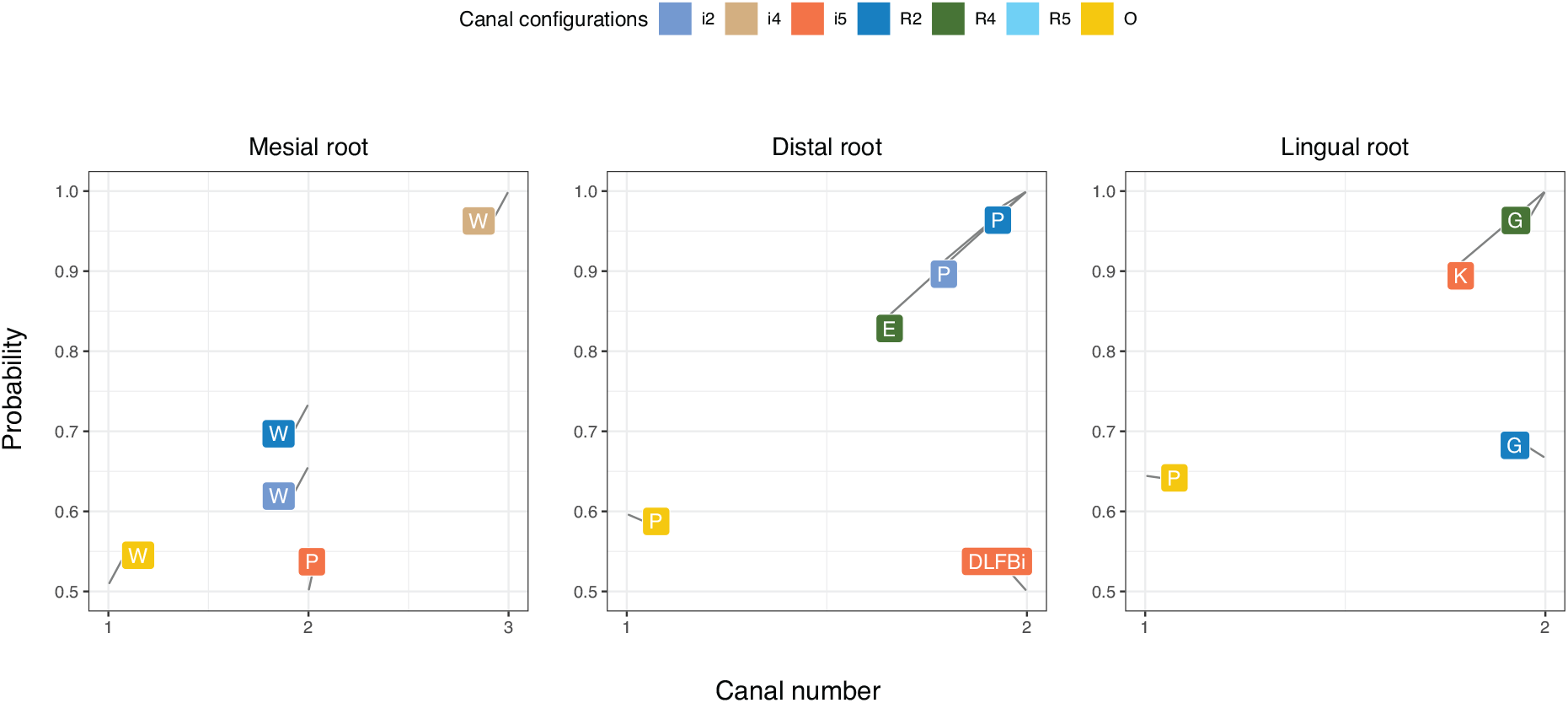
MLR of canal number to canal configuration and root morphology in M^1^s. Most frequent phenotype for mesial roots = wedge (W); distal roots = plate (P); and lingual roots = plate (P).

### Mandibular premolars and molars

All P_3_s used in this study were single rooted. The majority of double-canaled mandibular premolars are Tomes’ roots (Figure 19), while single-canaled premolars are plate-shaped. All Tomes’ shaped premolars have isthmus canal configuration while plate-shaped roots do not. A single oval canal predicts a plate-shaped single rooted mandibular premolar with 62.15% accuracy. Double-canaled mandibular premolars are either plate-shaped with a R4 canal configuration (99.99%) or a Tomes’ root. Isthmus canal i5 configurations predict the Tomes’ root morphology with 84.75.

**Figure 19:**
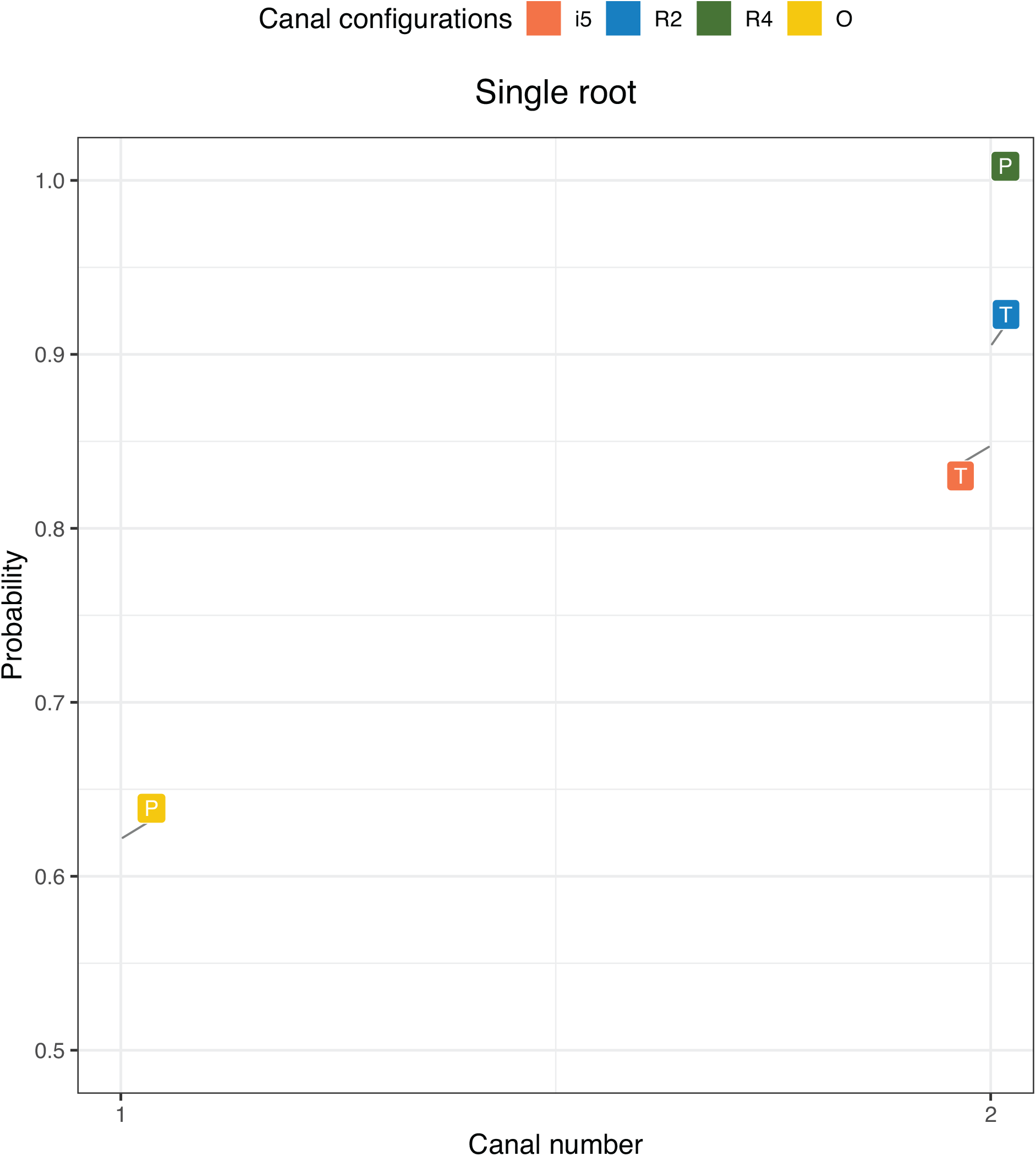
MLR of canal number to canal configuration and root morphology in sin single rooted P_3_s. Most frequent phenotype for single rooted P_3_s = plate (P).

Results of MLR for M_1_s are presented in Figure 20. A single round shaped canal has the greatest predictive power for plate shaped single-canaled mesial mandibular molar roots (100.00%). Amongst a number of canal configurations, both i2 and i3 isthmus canal configurations predict plate shaped double-canaled mesial roots at 72.22%. Distal roots are predominantly plate shaped with double canal variations. An i3 isthmus canal predicts the plate shaped morphology at 84.62%, while double-canaled R4 morphology predicts plate shaped morphology at 65.22%, followed by a number of isthmus canal variants. A single round canals predict globular shaped lingual roots (87.10%), and kidney shaped is predicted by an i2 isthmus canal morphology in Double-canaled variants (99.9%). There were not enough single rooted M_1_s or M_1_s with buccal roots for this study.

**Figure 20:**
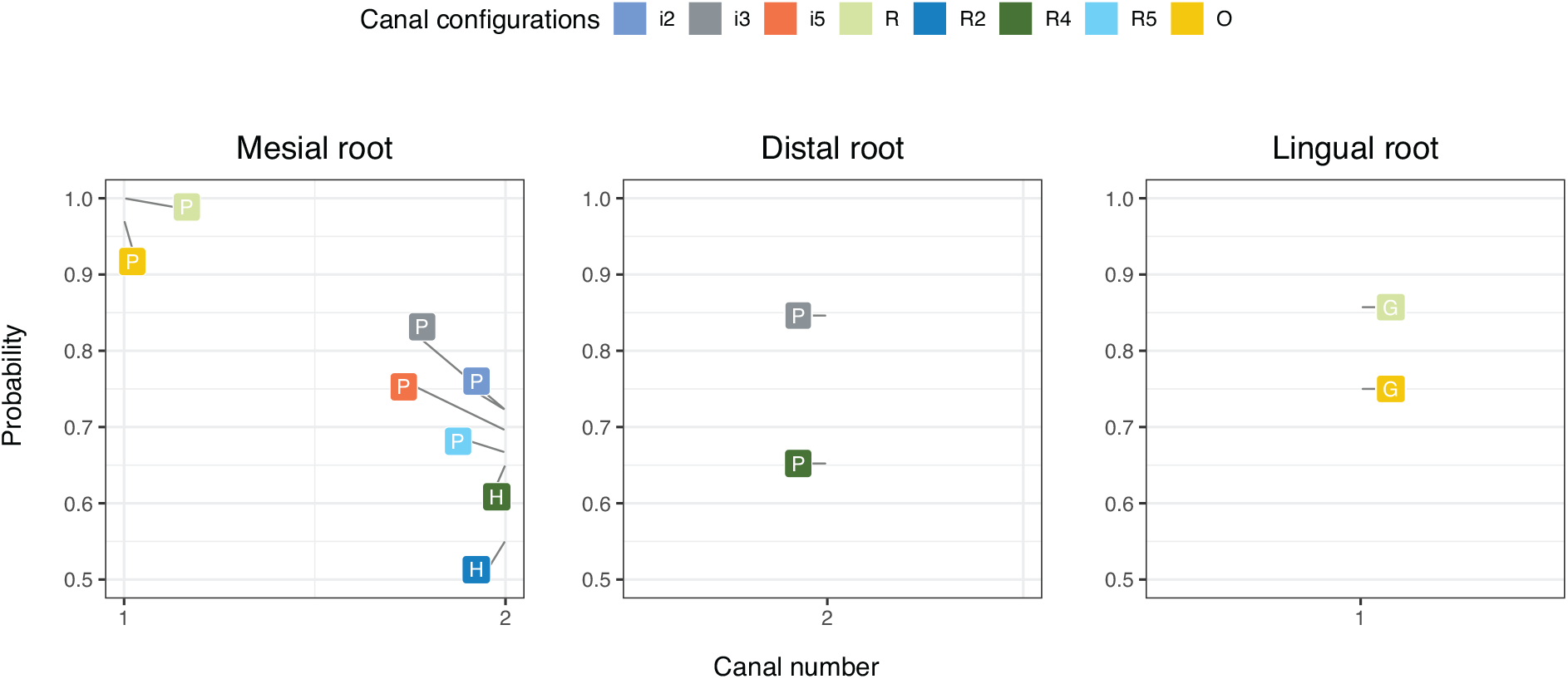
MLR of canal number to canal configuration and root morphology in M_1_ mesial, distal, and lingual roots.

Results of MLR for M_2_s with C-shaped roots and canals are presented in Figure 22. C-shaped molars are characterized by isthmus canals (Fan, Cheung, Fan, Gutmann, & Bian, 2004; Fernandes et al., 2014), and here they (i2, i3, i5) predict C-shaped roots 100.0% of the time. A single round canal predicts a single-rooted G-shaped morphology 99.99% of the time.

**Figure 21:**
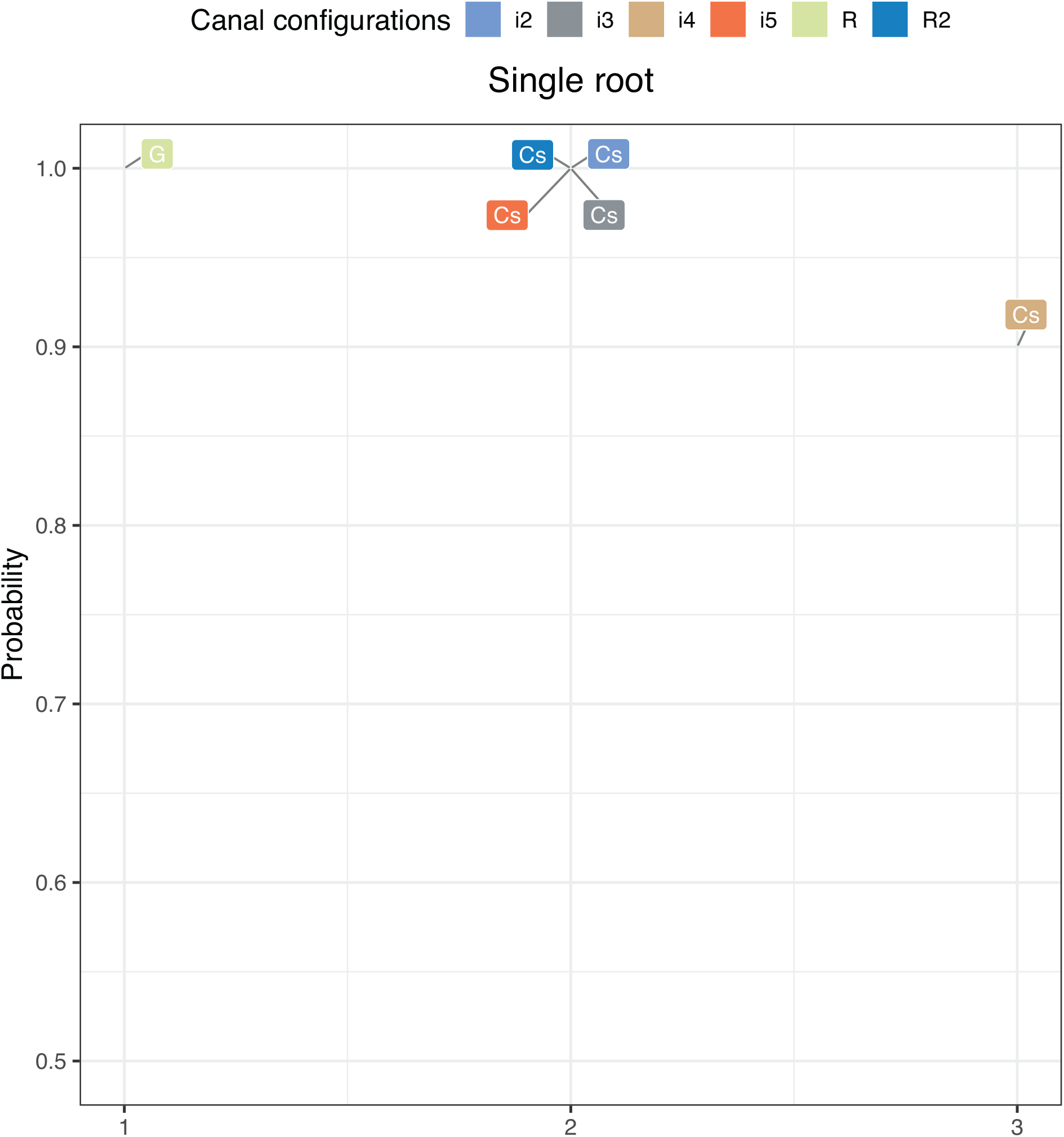
MLR of canal number to canal configuration and root morphology for M_2_ C-shaped roots and canals.

**Figure 22:**
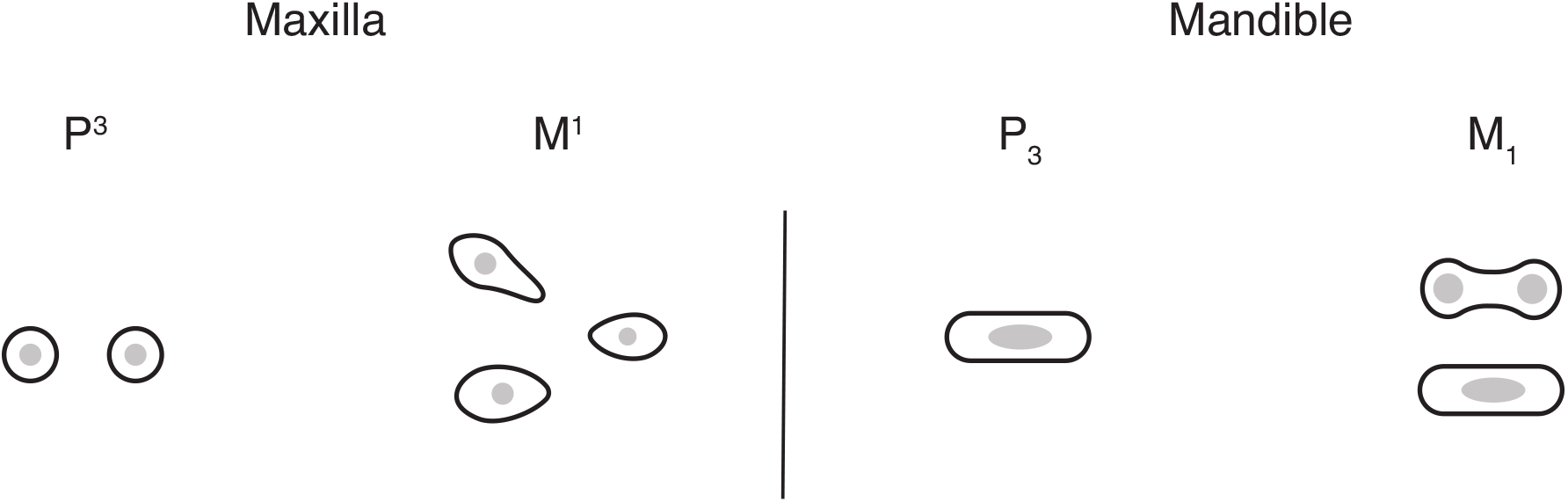
Most frequent phenotypes of external root morphology and internal canal count and configuration for key teeth from the maxilla (P ^3^ and M ^1^) and mandible (P_3_ and M_1_). Black line = external root morphology, solid gray = canal morphology and configuration. Figures are not in anatomical position but are aligned in a way that mimics their placement in the jaws.

Figure 22 shows the MFP for key teeth of the maxilla and mandible. The MFP of P^3^s is two globular roots with single round canals: one in the buccal position, the other in the lingual position. The MFP of M^1^s, is three roots: a wedge-shaped mesial root with a single round canal, and elliptical shaped distal and lingual roots with a single round canal. The MFP of mandibular P_3_s is a single plate shape root with an oval canal. For M_1_s, the MFP is an hourglass mesial root containing two round canals in an R2 configuration, and a plate-shaped distal root with a single oval canal.

## Discussion

This paper set out to (1) discover the relationship between canal number and configuration to external root morphology; (2) investigate if this relationship varies by tooth; and (3) investigate if root canal number and configuration predict external root morphology. It has been shown that there are MFPs for teeth within and between the maxilla and mandible; and that there is a significant associative and predictive power between internal and external tooth root morphologies.

### Predictive power of canal morphology and configuration on external root morphology

Results show that root canal number, morphology, and configuration can predict external root morphology. Results also indicate that certain external morphologies mirror internal morphologies in varying degrees. For example, globular roots are similar in shape to round canals, as are plate-shaped roots to oval canals, as well as C-shaped and Tomes’ roots with isthmus canals. Unsurprisingly, bifurcated forms always appear 2 canals (e.g., R2 configuration). Double canaled forms are stronger predictors of roots with an increased difference in height to width ratio (i.e., plate, hourglass, elliptical, wedge, kidney). This can be interpreted in several ways. The first is that a wider root is the product of multiple canals, and the relationship between canal number and root morphology is a spatial one. Canal formation precedes external root formation (Miller, 2013), and canal number has been shown to predict root number (Gellis & Foley, 2022). However, this interpretation is problematic. For example, single round canals predict plate shaped mesial roots of M_1_s at %100.00 (Figure 18).

Alternatively, while blood vessels do appear in locations that roots will eventually form (Miller, 2013), they may not be the prime determinants of root number and variation. The extension and fusion of opposing IRPs across the cervical foramen create multiple secondary foramina which, in turn, form multiple tooth roots (Kovacs, 1971; Orban & Bhaskar, 1980). It may be that number and orientation of IRP’s is responsible for tooth root dimensions and/or shape alone or in convert with the developing blood vessels. Future studies may be able to elucidate this by comparing location and measure of IRPs to root dimensions and morphology.

Roots function to anchor the teeth to the jaws, and to absorb and transmit the directional forces of mastication. It may simply be that that variations in root morphology are due to masticatory differences. In a study of M^1^ root forms in non-human primates and South-African robust and gracile Australopiths, root shape, size, and orientation were found to correlate with diet, bite force, and chewing pattern (Kupczik et al., 2018). Tooth crown size is already known to correlate to diet in hominins (Moggi-Cecchi & Boccone, 2007) and non-human primates (Kupczik et al., 2009; Spencer, 2003). However, the relationship between dietary strategy and masticatory forces to external root morphology and canal morphology and configuration are poorly understood.

Spatial, developmental, and functional explanations ned not be mutually exclusive. However, based on the on the predictive power of canal number, morphology, and configuration, the first explanation appears to be the strongest. Multiple canaled forms do predict wider and/or bifurcated roots, and these predictions coincide with many of the MFPs described in Figures 16 and 22. One root breaks this pattern - mesial wedge-shaped roots of maxillary molars. This root is predicted by nearly every double-canaled variant and morphology, none of which mirror its external form. This suggests, again, that the relationship between canals and external morphology is a more a spatial one. However, this may be due to certain roots having more ‘evolvability’ than others (discussed below).

### Constrained phenotypes and the most frequent phenotype

Certain forms are in the clear majority. These are plate-shaped roots with single oval shaped canals, and globular shaped roots with single, round shaped canals (Figure 6 -15). At least one of these external morphologies is found in all teeth in the maxilla and mandible with single and double canaled variants. Double canaled roots are more variable in their external morphologies and exhibit a number of canal configurations. This is less the case for the globular variant which rarely has more than one canal. The most common multi-canaled variant is the R2 configuration of two separate canals, followed by the R4 configuration of two distinct canals which are fused at their apices (Figure 5).

In multi-rooted forms, the pattern of external and internal morphologies is dissimilar between the molars of the maxilla and mandible (Figures 8-10 & 13-15). Multi-rooted teeth show significantly more variation in external morphologies than single rooted teeth. For example, mesial roots of multi-rooted M^1^s have 17 external and 19 internal morphologies compared to only 1 external and 1 internal morphology in single rooted M^1^s (Figure 8). In contrast, distal M^1^ roots are varied in their external morphology but constrained in their canal configurations; with most forms containing a single round canal.

While tooth roots possess a great deal of external and internal morphological variation, the diversity of individual roots in this sample is overwhelmingly constrained to several forms (Figures 16 and 22). Phenotypic variation is the raw material upon which selection and drift acts. However, phenotypic variation in organismal development is biased towards certain phenotypes (Arthur, 2004; J. M. Smith et al., 1985; Wilkins, 2007). While canal and root formation are comprised of a series of reciprocal cellular interactions (Jernvall & Thesleff, 2000), clusters of blood vessels entering the developing tooth coincide with the positions where roots will eventually form (Miller, 2013); and there is a clear relationship between canal and root number, in which canal number is either equal to or exceeds root number (Gellis & Foley, 2022). These observations of root development suggest that the structures that help determine root number and position are present early in tooth morphogenesis and play some part in the developmental bias of roots.

Taken in the above context, these results raise several important questions. The first is why are tooth root phenotypes so constrained? The most obvious answer is that, like tooth crowns, roots are under strong genetic control; and disturbances due to genetic alterations may lead to morphological defects or inhibit or cease development. Plate-shaped, oval-canaled, and globular shaped, round canaled roots are the MFP in the study sample. These forms are not only present in Plio-Pleistocene hominins and non-human primates (Hillson, 1996), but in early mammals of the Jurassic period as well (Z.-X. Luo & Wible, 2005; Z. X. Luo et al., 2015); suggesting a deep evolutionary history for these forms. Thus, it would appear that not only are root forms are developmentally and phenotypically constrained, but that these constraints are shared in early mammalian lineages as well.

These constraints may be due to the evolutionary adaptability or, ‘evolvability’ of tooth roots. The capacity to generate novel, heritable phenotypic variation defines a trait’s evolvability (Kirschner & Gerhart, 1998). At the cellular level, regulatory proteins act to promote or inhibit the number of random mutational steps needed to generate novel regulatory mechanisms (Kirschner & Gerhart, 1998). These regulatory processes are relevant to evolutionary processes as they can reduce constraints on change and the accumulation of non-lethal variants. The greater the number and specificity of a protein’s functional requirements, the more resistant they are to change. Additionally, a protein’s structural stability enhances its capacity to evolve by allowing it to accept a wider range of beneficial mutations while retaining its ability to fold to its original structure (Bloom et al., 2006). Together, the number, specificity, and stability of proteins helps explain evolution’s extensive morphological and physiological diversity in light of taxon-wide conservation of core genetic, cellular, and developmental processes.

The concepts of developmental bias, evolvability, and phenotypic constraint, help inform the second question - if tooth roots are so phenotypically constrained, why do certain roots exhibit higher levels of diversity? Based on the above results, the presence of additional canals and canal configurations are responsible for different external root forms. This suggests that multiple canals, which conserve core genetic, cellular, and developmental processes of tooth morphogenesis, may have more ‘evolvability’ than single canaled forms. However, an equally plausible explanation is that canals act as a scaffolding around which the external components of roots differentiate and grow, and ultimately take their final shape from. The two need not be mutually exclusive, as the processes underlying the entirety of root morphogenesis are reciprocal (Jernvall and Thesleff, 2000).

In light of these two questions, what is interesting is that certain roots seem to be more variable than others. The most variable is the mesial root of maxillary molars which exhibits a wide range of external morphologies and canal configurations (Figures 8, 9, & 10). Why this root should exhibit so much variation while others do not is unknown. Additionally, its external morphology does not mirror internal morphology as there is no wedge-shaped canal, and the morphology is retained regardless of canal number or configuration. It has been observed that mandibular premolars are the most variable teeth for humans, fossil hominins, and non-human primates (Wood and Abbott, 1983; Wood et al., 1988; Shields, 2005; Moore et al., 2013, 2015, 2016; Emonet and Kullmer, 2014). However, this study suggests that it is maxillary molars that are the most variable, not only in root number, but in exterior root morphology, and canal number, morphology, and configuration.

### The use of key teeth vs. the total phenotypic set

The use of ‘key teeth’ has long history in dental anthropological studies. From a statistical analysis standpoint this is to avoid issues with how linearly correlated and/or non-independent variables violate statistical assumptions of independence. In modern humans, this approach is problematic as (a) it leaves out a wide range of tooth root and canal morphologies that are either not found in ‘key teeth’ (e.g., M_2_s with C-shaped roots and canals), or have a higher degree of expression than the most mesial member of a morphogenetic field (e.g., fused roots); and (b) many of these traits, such as C-shaped molars, Tomes’ roots, and three rooted M_1_s have an ethnic component, with high frequencies Asian, European, and Sino-American populations respectively (Tomes, 1889; Wang et al., 2012; Ballullaya et al., 2013; Fernandes et al., 2014). These traits are of great importance for population studies based on tooth root phenotypes, and it may be that the inclusion of all teeth rather than just ‘key teeth’ are necessary for generating higher resolution population analyses.

From an adaptive framework, ‘key teeth’ are also problematic as they potentially biased towards adaptive traits at the expense of ‘populations specific’ traits. In humans, M^1^s/M_1_s, the ‘key teeth’ in molar morphogenetic fields, have the highest bite-force magnitudes of all teeth (Ledogar et al., 2016). Results show that M_1_s/M_1_s have the most internal and external morphologies (Figure 8). However, it is unknown if this is due to adaptive pressures of dietary differences within and between the geographic groups used in this study, some other selective pressure. The linear relationship between canal and root number has been shown to significantly vary between populations (Gellis & Foley, 2022). Further work on tooth root trait expression and frequency, and how these correlate with masticatory behaviours will need to be carried out in order to determine how and what traits are adaptive versus which ones are population specific.

## Conclusions

This paper presents a novel investigation into the relationship between canal number, morphology, and configuration to external root morphology. The most frequent phenotypes are described for post canine teeth of the maxilla, mandible, and jaws combined. Results indicate that certain canal morphologies and configurations are strongly associated with and can predict external root morphology. It is unclear why internal and external variation is distributed the way it is, or why the internal and external structures of some roots are more variable than others. Future studies will need to further clarify the underlying developmental mechanisms of tooth root morphogenesis and consider biomechanical and dietary factors as well.

## Acknowledgements

I would like to thank Professor Marta Miraźon-Lahr and Dr. Frances Rivera for permitting use of their CT scans from the Duckworth Collection at the Leverhulme Centre for Human Evolutionary Studies, University of Cambridge; and Dr. Lynn Copes for permitting use of her CT scans, collected for her PhD dissertation, from the Smithsonian National Museum of Natural History and American Museum of Natural History. I would also like to thank the Duckworth Laboratory, University of Cambridge, for permission to study material within its collections. The author has no conflict of interest to declare.

## Data Availability Statement

The data that support the findings of this study are openly available from the Open Science Foundation *Tooth Root Phenotypic Data* set; at https://doi.org/10.17605/OSF.IO/9YNUR

