## Supplementary_materials for "Canal number and configuration are predictors of external root morphology"

### Supplementary Material

**Table A:** Collection information for individuals used in this study

| ID | Collection | Sex | Years BP* | G1† | G2 | G3 | G4 | G5 |
|---|---|---|---|---|---|---|---|---|
| 99_1_102 | AMNH | F | ~1600 | Sino-Americas | North America | American Arctic | Alaska | Ipiutak |
| 99_1_103 | AMNH | M | ~1600 | Sino-Americas | North America | American Arctic | Alaska | Ipiutak |
| 99_1_105 | AMNH | M | ~1600 | Sino-Americas | North America | American Arctic | Alaska | Ipiutak |
| 99_1_161 | AMNH | F | ~1600 | Sino-Americas | North America | American Arctic | Alaska | Ipiutak |
| 99_1_163 | AMNH | F | ~1600 | Sino-Americas | North America | American Arctic | Alaska | Ipiutak |
| 99_1_165 | AMNH | M | ~1600 | Sino-Americas | North America | American Arctic | Alaska | Ipiutak |
| 99_1_166 | AMNH | M | ~1600 | Sino-Americas | North America | American Arctic | Alaska | Ipiutak |
| 99_1_168 | AMNH | F | ~1600 | Sino-Americas | North America | American Arctic | Alaska | Ipiutak |
| 99_1_181 | AMNH | M | ~1600 | Sino-Americas | North America | American Arctic | Alaska | Ipiutak |
| 99_1_192 | AMNH | M | ~1600 | Sino-Americas | North America | American Arctic | Alaska | Ipiutak |
| 99_1_194 | AMNH | M | ~1600 | Sino-Americas | North America | American Arctic | Alaska | Ipiutak |
| 99_1_196 | AMNH | M | ~1600 | Sino-Americas | North America | American Arctic | Alaska | Ipiutak |
| 99_1_197 | AMNH | F | ~1600 | Sino-Americas | North America | American Arctic | Alaska | Ipiutak |
| 99_1_198 | AMNH | NA | ~1600 | Sino-Americas | North America | American Arctic | Alaska | Ipiutak |
| 99_1_201 | AMNH | F | ~1600 | Sino-Americas | North America | American Arctic | Alaska | Ipiutak |
| 99_1_252 | AMNH | M | ~1600 | Sino-Americas | North America | American Arctic | Alaska | Ipiutak |
| 99_1_684 | AMNH | F | ~1600 | Sino-Americas | North America | American Arctic | Alaska | Ipiutak |
| 99_1_72 | AMNH | M | ~1600 | Sino-Americas | North America | American Arctic | Alaska | Ipiutak |
| 99_1_80 | AMNH | M | ~1600 | Sino-Americas | North America | American Arctic | Alaska | Ipiutak |
| 99_1_90 | AMNH | F | ~1600 | Sino-Americas | North America | American Arctic | Alaska | Ipiutak |
| 99_1_92 | AMNH | M | ~1600 | Sino-Americas | North America | American Arctic | Alaska | Ipiutak |
| 99_1_93 | AMNH | M | ~1600 | Sino-Americas | North America | American Arctic | Alaska | Ipiutak |
| 99_1_94 | AMNH | M | ~1600 | Sino-Americas | North America | American Arctic | Alaska | Ipiutak |
| 99_1_95 | AMNH | NA | ~1600 | Sino-Americas | North America | American Arctic | Alaska | Ipiutak |
| 99_1_256 | AMNH | M | Unkw | Sino-Americas | North America | American Arctic | Alaska | NA |
| 99_1_569 | AMNH | F | Unkw | Sino-Americas | North America | American Arctic | Alaska | NA |
| 99_1_575 | AMNH | F | Unkw | Sino-Americas | North America | American Arctic | Alaska | NA |
| 99_1_586 | AMNH | F | Unkw | Sino-Americas | North America | American Arctic | Alaska | NA |
| 99_1_592 | AMNH | F | Unkw | Sino-Americas | North America | American Arctic | Alaska | NA |
| 99_1_606 | AMNH | F | Unkw | Sino-Americas | North America | American Arctic | Alaska | NA |
| 99_1_607 | AMNH | M | Unkw | Sino-Americas | North America | American Arctic | Alaska | NA |
| 99_1_608 | AMNH | F | Unkw | Sino-Americas | North America | American Arctic | Alaska | NA |
| 99_1_61 | AMNH | M | Unkw | Sino-Americas | North America | American Arctic | Alaska | NA |

| | | | | | | | | |
|---|---|---|---|---|---|---|---|---|
| 99_1_619 | AMNH | M | Unkw | Sino-Americas | North America | American Arctic | Alaska | NA |
| 99_1_671 | AMNH | F | ~800 | Sino-Americas | North America | American Arctic | Alaska | Tigara |
| 99_1_672 | AMNH | F | ~800 | Sino-Americas | North America | American Arctic | Alaska | Tigara |
| 99_1_88 | AMNH | F | ~800 | Sino-Americas | North America | American Arctic | Alaska | Tigara |
| 99_1_96A | AMNH | M | ~800 | Sino-Americas | North America | American Arctic | Alaska | Tigara |
| 99_1_224 | AMNH | M | ~800 | Sino-Americas | North America | American Arctic | Alaska | Tigara |
| 99_1_225 | AMNH | M | ~800 | Sino-Americas | North America | American Arctic | Alaska | Tigara |
| 99_1_228 | AMNH | M | ~800 | Sino-Americas | North America | American Arctic | Alaska | Tigara |
| 99_1_233 | AMNH | M | ~800 | Sino-Americas | North America | American Arctic | Alaska | Tigara |
| 99_1_234 | AMNH | F | ~800 | Sino-Americas | North America | American Arctic | Alaska | Tigara |
| 99_1_235 | AMNH | M | ~800 | Sino-Americas | North America | American Arctic | Alaska | Tigara |
| 99_1_237 | AMNH | M | ~800 | Sino-Americas | North America | American Arctic | Alaska | Tigara |
| 99_1_240 | AMNH | F | ~800 | Sino-Americas | North America | American Arctic | Alaska | Tigara |
| 99_1_242 | AMNH | M | ~800 | Sino-Americas | North America | American Arctic | Alaska | Tigara |
| 99_1_246 | AMNH | F | ~800 | Sino-Americas | North America | American Arctic | Alaska | Tigara |
| 99_1_248 | AMNH | F | ~800 | Sino-Americas | North America | American Arctic | Alaska | Tigara |
| 99_1_265 | AMNH | M | ~800 | Sino-Americas | North America | American Arctic | Alaska | Tigara |
| 99_1_268 | AMNH | F | ~800 | Sino-Americas | North America | American Arctic | Alaska | Tigara |
| 99_1_272 | AMNH | M | ~800 | Sino-Americas | North America | American Arctic | Alaska | Tigara |
| 99_1_276 | AMNH | F | ~800 | Sino-Americas | North America | American Arctic | Alaska | Tigara |
| 99_1_277 | AMNH | M | ~800 | Sino-Americas | North America | American Arctic | Alaska | Tigara |
| 99_1_278 | AMNH | M | ~800 | Sino-Americas | North America | American Arctic | Alaska | Tigara |
| 99_1_282 | AMNH | F | ~800 | Sino-Americas | North America | American Arctic | Alaska | Tigara |
| 99_1_393 | AMNH | M | ~800 | Sino-Americas | North America | American Arctic | Alaska | Tigara |
| 99_1_394 | AMNH | F | ~800 | Sino-Americas | North America | American Arctic | Alaska | Tigara |
| 99_1_395 | AMNH | M | ~800 | Sino-Americas | North America | American Arctic | Alaska | Tigara |
| 99_1_396 | AMNH | M | ~800 | Sino-Americas | North America | American Arctic | Alaska | Tigara |
| 99_1_397 | AMNH | M | ~800 | Sino-Americas | North America | American Arctic | Alaska | Tigara |
| 99_1_401 | AMNH | F | ~800 | Sino-Americas | North America | American Arctic | Alaska | Tigara |
| 99_1_406 | AMNH | NA | ~800 | Sino-Americas | North America | American Arctic | Alaska | Tigara |
| 99_1_407 | AMNH | M | ~800 | Sino-Americas | North America | American Arctic | Alaska | Tigara |
| 99_1_408 | AMNH | M | ~800 | Sino-Americas | North America | American Arctic | Alaska | Tigara |
| 99_1_409 | AMNH | F | ~800 | Sino-Americas | North America | American Arctic | Alaska | Tigara |
| 99_1_410 | AMNH | M | ~800 | Sino-Americas | North America | American Arctic | Alaska | Tigara |
| 99_1_412 | AMNH | F | ~800 | Sino-Americas | North America | American Arctic | Alaska | Tigara |
| 99_1_414 | AMNH | F | ~800 | Sino-Americas | North America | American Arctic | Alaska | Tigara |
| 99_1_417 | AMNH | F | ~800 | Sino-Americas | North America | American Arctic | Alaska | Tigara |
| 99_1_421 | AMNH | F | ~800 | Sino-Americas | North America | American Arctic | Alaska | Tigara |

| | | | | | | | | |
|---|---|---|---|---|---|---|---|---|
| 99_1_422 | AMNH | F | ~800 | Sino-Americas | North America | American Arctic | Alaska | Tigara |
| 99_1_423 | AMNH | M | ~800 | Sino-Americas | North America | American Arctic | Alaska | Tigara |
| 99_1_424 | AMNH | F | ~800 | Sino-Americas | North America | American Arctic | Alaska | Tigara |
| 99_1_425 | AMNH | F | ~800 | Sino-Americas | North America | American Arctic | Alaska | Tigara |
| 99_1_426 | AMNH | M | ~800 | Sino-Americas | North America | American Arctic | Alaska | Tigara |
| 99_1_428 | AMNH | M | ~800 | Sino-Americas | North America | American Arctic | Alaska | Tigara |
| 99_1_429 | AMNH | F | ~800 | Sino-Americas | North America | American Arctic | Alaska | Tigara |
| 99_1_431 | AMNH | M | ~800 | Sino-Americas | North America | American Arctic | Alaska | Tigara |
| 99_1_432 | AMNH | M | ~800 | Sino-Americas | North America | American Arctic | Alaska | Tigara |
| 99_1_434 | AMNH | M | ~800 | Sino-Americas | North America | American Arctic | Alaska | Tigara |
| 99_1_438 | AMNH | F | ~800 | Sino-Americas | North America | American Arctic | Alaska | Tigara |
| 99_1_439 | AMNH | M | ~800 | Sino-Americas | North America | American Arctic | Alaska | Tigara |
| 99_1_441 | AMNH | M | ~800 | Sino-Americas | North America | American Arctic | Alaska | Tigara |
| 99_1_442 | AMNH | F | ~800 | Sino-Americas | North America | American Arctic | Alaska | Tigara |
| 99_1_444 | AMNH | F | ~800 | Sino-Americas | North America | American Arctic | Alaska | Tigara |
| 99_1_445 | AMNH | M | ~800 | Sino-Americas | North America | American Arctic | Alaska | Tigara |
| 99_1_446 | AMNH | M | ~800 | Sino-Americas | North America | American Arctic | Alaska | Tigara |
| 99_1_447 | AMNH | M | ~800 | Sino-Americas | North America | American Arctic | Alaska | Tigara |
| 99_1_449 | AMNH | F | ~800 | Sino-Americas | North America | American Arctic | Alaska | Tigara |
| 99_1_450 | AMNH | M | ~800 | Sino-Americas | North America | American Arctic | Alaska | Tigara |
| 99_1_452 | AMNH | M | ~800 | Sino-Americas | North America | American Arctic | Alaska | Tigara |
| 99_1_453 | AMNH | M | ~800 | Sino-Americas | North America | American Arctic | Alaska | Tigara |
| 99_1_454 | AMNH | F | ~800 | Sino-Americas | North America | American Arctic | Alaska | Tigara |
| 99_1_458 | AMNH | F | ~800 | Sino-Americas | North America | American Arctic | Alaska | Tigara |
| 99_1_459 | AMNH | F | ~800 | Sino-Americas | North America | American Arctic | Alaska | Tigara |
| 99_1_460 | AMNH | M | ~800 | Sino-Americas | North America | American Arctic | Alaska | Tigara |
| 99_1_463 | AMNH | F | ~800 | Sino-Americas | North America | American Arctic | Alaska | Tigara |
| 99_1_464 | AMNH | F | ~800 | Sino-Americas | North America | American Arctic | Alaska | Tigara |
| 99_1_465 | AMNH | F | ~800 | Sino-Americas | North America | American Arctic | Alaska | Tigara |
| 99_1_466 | AMNH | F | ~800 | Sino-Americas | North America | American Arctic | Alaska | Tigara |
| 99_1_468 | AMNH | M | ~800 | Sino-Americas | North America | American Arctic | Alaska | Tigara |
| 99_1_472 | AMNH | M | ~800 | Sino-Americas | North America | American Arctic | Alaska | Tigara |
| 99_1_473 | AMNH | F | ~800 | Sino-Americas | North America | American Arctic | Alaska | Tigara |
| 99_1_474 | AMNH | M | ~800 | Sino-Americas | North America | American Arctic | Alaska | Tigara |
| 99_1_475 | AMNH | M | ~800 | Sino-Americas | North America | American Arctic | Alaska | Tigara |
| 99_1_476 | AMNH | M | ~800 | Sino-Americas | North America | American Arctic | Alaska | Tigara |
| 99_1_478 | AMNH | M | ~800 | Sino-Americas | North America | American Arctic | Alaska | Tigara |
| 99_1_480 | AMNH | F | ~800 | Sino-Americas | North America | American Arctic | Alaska | Tigara |

| | | | | | | | | |
|---|---|---|---|---|---|---|---|---|
| 99_1_481 | AMNH | M | ~800 | Sino-Americas | North America | American Arctic | Alaska | Tigara |
| 99_1_482 | AMNH | F | ~800 | Sino-Americas | North America | American Arctic | Alaska | Tigara |
| 99_1_485 | AMNH | F | ~800 | Sino-Americas | North America | American Arctic | Alaska | Tigara |
| 99_1_488 | AMNH | M | ~800 | Sino-Americas | North America | American Arctic | Alaska | Tigara |
| 99_1_489 | AMNH | F | ~800 | Sino-Americas | North America | American Arctic | Alaska | Tigara |
| 99_1_490 | AMNH | M | ~800 | Sino-Americas | North America | American Arctic | Alaska | Tigara |
| 99_1_492 | AMNH | M | ~800 | Sino-Americas | North America | American Arctic | Alaska | Tigara |
| 99_1_493 | AMNH | M | ~800 | Sino-Americas | North America | American Arctic | Alaska | Tigara |
| 99_1_494 | AMNH | F | ~800 | Sino-Americas | North America | American Arctic | Alaska | Tigara |
| 99_1_495 | AMNH | M | ~800 | Sino-Americas | North America | American Arctic | Alaska | Tigara |
| 99_1_496 | AMNH | F | ~800 | Sino-Americas | North America | American Arctic | Alaska | Tigara |
| 99_1_497 | AMNH | M | ~800 | Sino-Americas | North America | American Arctic | Alaska | Tigara |
| 99_1_498 | AMNH | M | ~800 | Sino-Americas | North America | American Arctic | Alaska | Tigara |
| 99_1_499 | AMNH | M | ~800 | Sino-Americas | North America | American Arctic | Alaska | Tigara |
| 99_1_500 | AMNH | F | ~800 | Sino-Americas | North America | American Arctic | Alaska | Tigara |
| 99_1_502 | AMNH | F | ~800 | Sino-Americas | North America | American Arctic | Alaska | Tigara |
| 99_1_504 | AMNH | M | ~800 | Sino-Americas | North America | American Arctic | Alaska | Tigara |
| 99_1_506 | AMNH | M | ~800 | Sino-Americas | North America | American Arctic | Alaska | Tigara |
| 99_1_507 | AMNH | F | ~800 | Sino-Americas | North America | American Arctic | Alaska | Tigara |
| 99_1_508 | AMNH | M | ~800 | Sino-Americas | North America | American Arctic | Alaska | Tigara |
| 99_1_509 | AMNH | M | ~800 | Sino-Americas | North America | American Arctic | Alaska | Tigara |
| 99_1_510 | AMNH | F | ~800 | Sino-Americas | North America | American Arctic | Alaska | Tigara |
| 99_1_511 | AMNH | M | ~800 | Sino-Americas | North America | American Arctic | Alaska | Tigara |
| 99_1_513 | AMNH | F | ~800 | Sino-Americas | North America | American Arctic | Alaska | Tigara |
| 99_1_514 | AMNH | F | ~800 | Sino-Americas | North America | American Arctic | Alaska | Tigara |
| 99_1_515 | AMNH | F | ~800 | Sino-Americas | North America | American Arctic | Alaska | Tigara |
| 99_1_517 | AMNH | F | ~800 | Sino-Americas | North America | American Arctic | Alaska | Tigara |
| 99_1_520 | AMNH | M | ~800 | Sino-Americas | North America | American Arctic | Alaska | Tigara |
| 99_1_521 | AMNH | M | ~800 | Sino-Americas | North America | American Arctic | Alaska | Tigara |
| 99_1_522 | AMNH | F | ~800 | Sino-Americas | North America | American Arctic | Alaska | Tigara |
| 99_1_523 | AMNH | F | ~800 | Sino-Americas | North America | American Arctic | Alaska | Tigara |
| 99_1_524 | AMNH | F | ~800 | Sino-Americas | North America | American Arctic | Alaska | Tigara |
| 99_1_525 | AMNH | M | ~800 | Sino-Americas | North America | American Arctic | Alaska | Tigara |
| 99_1_526 | AMNH | F | ~800 | Sino-Americas | North America | American Arctic | Alaska | Tigara |
| 99_1_527 | AMNH | M | ~800 | Sino-Americas | North America | American Arctic | Alaska | Tigara |
| 99_1_532 | AMNH | F | ~800 | Sino-Americas | North America | American Arctic | Alaska | Tigara |
| 99_1_535 | AMNH | M | ~800 | Sino-Americas | North America | American Arctic | Alaska | Tigara |
| 99_1_536 | AMNH | M | ~800 | Sino-Americas | North America | American Arctic | Alaska | Tigara |

| | | | | | | | | |
|---|---|---|---|---|---|---|---|---|
| 99_1_541 | AMNH | M | ~800 | Sino-Americas | North America | American Arctic | Alaska | Tigara |
| 99_1_542 | AMNH | M | ~800 | Sino-Americas | North America | American Arctic | Alaska | Tigara |
| 99_1_543 | AMNH | F | ~800 | Sino-Americas | North America | American Arctic | Alaska | Tigara |
| 99_1_544 | AMNH | F | ~800 | Sino-Americas | North America | American Arctic | Alaska | Tigara |
| 99_1_546 | AMNH | F | ~800 | Sino-Americas | North America | American Arctic | Alaska | Tigara |
| 99_1_549 | AMNH | M | ~800 | Sino-Americas | North America | American Arctic | Alaska | Tigara |
| 99_1_551 | AMNH | F | ~800 | Sino-Americas | North America | American Arctic | Alaska | Tigara |
| 99_1_552 | AMNH | M | ~800 | Sino-Americas | North America | American Arctic | Alaska | Tigara |
| 99_1_643 | AMNH | F | ~800 | Sino-Americas | North America | American Arctic | Alaska | Tigara |
| 99_1_644 | AMNH | M | ~800 | Sino-Americas | North America | American Arctic | Alaska | Tigara |
| 99_1_666 | AMNH | F | ~800 | Sino-Americas | North America | American Arctic | Alaska | Tigara |
| 99_1_667 | AMNH | F | ~800 | Sino-Americas | North America | American Arctic | Alaska | Tigara |
| 99_1_675 | AMNH | F | ~800 | Sino-Americas | North America | American Arctic | Alaska | Tigara |
| 99_1_69 | AMNH | M | ~800 | Sino-Americas | North America | American Arctic | Alaska | Tigara |
| 226086 | NMNH | F | Unkw | Sahul-Pacific | Oceania | Australia | Victoria | Hexham |
| 226087 | NMNH | F | Unkw | Sahul-Pacific | Oceania | Australia | Victoria | Hexham |
| 226089 | NMNH | M | Unkw | Sahul-Pacific | Oceania | Australia | South Australia | Murray River |
| 226090 | NMNH | F | Unkw | Sahul-Pacific | Oceania | Australia | South Australia | Murray River |
| 329778 | NMNH | F | Unkw | Sahul-Pacific | Oceania | Australia | South Australia | Cape Spencer Aborigine |
| 329779 | NMNH | M | Unkw | Sahul-Pacific | Oceania | Australia | South Australia | Murray River |
| 330604 | NMNH | M | Unkw | Sahul-Pacific | Oceania | Australia | Western Australia | Derby Coast Aborigine |
| 331242 | NMNH | M | Unkw | Sahul-Pacific | Oceania | Australia | Central Australia | Aborigine |
| 331243 | NMNH | M | Unkw | Sahul-Pacific | Oceania | Australia | South Australia | Plympton Aborigine |
| 331247 | NMNH | F | Unkw | Sahul-Pacific | Oceania | Australia | South Australia | Swanport Aborigine |
| 344711 | NMNH | M | Unkw | Sahul-Pacific | Oceania | Australia | Victoria | Loddon River Aborigine |
| 344712 | NMNH | F | Unkw | Sahul-Pacific | Oceania | Australia | Victoria | Mortlake Aborigine |
| 344713 | NMNH | F | Unkw | Sahul-Pacific | Oceania | Australia | Victoria | Murray River Aborigine |
| 344714 | NMNH | F | Unkw | Sahul-Pacific | Oceania | Australia | Victoria | Murray River Aborigine |
| 344715 | NMNH | F | Unkw | Sahul-Pacific | Oceania | Australia | Victoria | Murray River Aborigine |
| 350096 | NMNH | M | Unkw | Sahul-Pacific | Oceania | Australia | South Australia | Aborigine |
| 242687 | NMNH | M | Unkw | Sino-Americas | North America | American Arctic | Greenland | Inuit |
| 242690 | NMNH | F | Unkw | Sino-Americas | North America | American Arctic | Greenland | Inuit |
| 242693 | NMNH | F | Unkw | Sino-Americas | North America | American Arctic | Greenland | Inuit |
| 242694 | NMNH | F | Unkw | Sino-Americas | North America | American Arctic | Greenland | Inuit |
| 242695 | NMNH | M | Unkw | Sino-Americas | North America | American Arctic | Greenland | Inuit |
| 242697 | NMNH | M | Unkw | Sino-Americas | North America | American Arctic | Greenland | Inuit |
| 242698 | NMNH | M | Unkw | Sino-Americas | North America | American Arctic | Greenland | Inuit |
| 242699 | NMNH | F | Unkw | Sino-Americas | North America | American Arctic | Greenland | Inuit |
| 242702 | NMNH | F | Unkw | Sino-Americas | North America | American Arctic | Greenland | Inuit |

| | | | | | | | | |
|---|---|---|---|---|---|---|---|---|
| 242704 | NMNH | F | Unkw | Sino-Americas | North America | American Arctic | Greenland | Inuit |
| 242706 | NMNH | M | Unkw | Sino-Americas | North America | American Arctic | Greenland | Inuit |
| 242707 | NMNH | M | Unkw | Sino-Americas | North America | American Arctic | Greenland | Inuit |
| 242708 | NMNH | F | Unkw | Sino-Americas | North America | American Arctic | Greenland | Inuit |
| 242709 | NMNH | M | Unkw | Sino-Americas | North America | American Arctic | Greenland | Inuit |
| 242710 | NMNH | F | Unkw | Sino-Americas | North America | American Arctic | Greenland | Inuit |
| 242711 | NMNH | F | Unkw | Sino-Americas | North America | American Arctic | Greenland | Inuit |
| 242712 | NMNH | F | Unkw | Sino-Americas | North America | American Arctic | Greenland | Inuit |
| 242714 | NMNH | F | Unkw | Sino-Americas | North America | American Arctic | Greenland | Inuit |
| 242716 | NMNH | M | Unkw | Sino-Americas | North America | American Arctic | Greenland | Inuit |
| 242717 | NMNH | F | Unkw | Sino-Americas | North America | American Arctic | Greenland | Inuit |
| 242718 | NMNH | F | Unkw | Sino-Americas | North America | American Arctic | Greenland | Inuit |
| 242719 | NMNH | F | Unkw | Sino-Americas | North America | American Arctic | Greenland | Inuit |
| 242720 | NMNH | F | Unkw | Sino-Americas | North America | American Arctic | Greenland | Inuit |
| 242721 | NMNH | M | Unkw | Sino-Americas | North America | American Arctic | Greenland | Inuit |
| 242724 | NMNH | F | Unkw | Sino-Americas | North America | American Arctic | Greenland | Inuit |
| 242725 | NMNH | F | Unkw | Sino-Americas | North America | American Arctic | Greenland | Inuit |
| 242726 | NMNH | M | Unkw | Sino-Americas | North America | American Arctic | Greenland | Inuit |
| 242729 | NMNH | M | Unkw | Sino-Americas | North America | American Arctic | Greenland | Inuit |
| 242730 | NMNH | M | Unkw | Sino-Americas | North America | American Arctic | Greenland | Inuit |
| 242731 | NMNH | F | Unkw | Sino-Americas | North America | American Arctic | Greenland | Inuit |
| 242732 | NMNH | M | Unkw | Sino-Americas | North America | American Arctic | Greenland | Inuit |
| 242733 | NMNH | M | Unkw | Sino-Americas | North America | American Arctic | Greenland | Inuit |
| 242734 | NMNH | M | Unkw | Sino-Americas | North America | American Arctic | Greenland | Inuit |
| 242735 | NMNH | F | Unkw | Sino-Americas | North America | American Arctic | Greenland | Inuit |
| 242736 | NMNH | F | Unkw | Sino-Americas | North America | American Arctic | Greenland | Inuit |
| 242738 | NMNH | F | Unkw | Sino-Americas | North America | American Arctic | Greenland | Inuit |
| 242741 | NMNH | F | Unkw | Sino-Americas | North America | American Arctic | Greenland | Inuit |
| 242742 | NMNH | F | Unkw | Sino-Americas | North America | American Arctic | Greenland | Inuit |
| 242743 | NMNH | F | Unkw | Sino-Americas | North America | American Arctic | Greenland | Inuit |
| 242744 | NMNH | M | Unkw | Sino-Americas | North America | American Arctic | Greenland | Inuit |
| 242745 | NMNH | F | Unkw | Sino-Americas | North America | American Arctic | Greenland | Inuit |
| 242746 | NMNH | F | Unkw | Sino-Americas | North America | American Arctic | Greenland | Inuit |
| 242747 | NMNH | M | Unkw | Sino-Americas | North America | American Arctic | Greenland | Inuit |
| 242748 | NMNH | M | Unkw | Sino-Americas | North America | American Arctic | Greenland | Inuit |
| 242757 | NMNH | F | Unkw | Sino-Americas | North America | American Arctic | Greenland | Inuit |
| 242758 | NMNH | F | Unkw | Sino-Americas | North America | American Arctic | Greenland | Inuit |
| 242759 | NMNH | F | Unkw | Sino-Americas | North America | American Arctic | Greenland | Inuit |

| | | | | | | | | |
|---|---|---|---|---|---|---|---|---|
| 242760 | NMNH | M | Unkw | Sino-Americas | North America | American Arctic | Greenland | Inuit |
| 242761 | NMNH | M | Unkw | Sino-Americas | North America | American Arctic | Greenland | Inuit |
| 242831 | NMNH | M | Unkw | Sino-Americas | North America | American Arctic | Greenland | Inuit |
| 242832 | NMNH | M | Unkw | Sino-Americas | North America | American Arctic | Greenland | Inuit |
| 242835 | NMNH | F | Unkw | Sino-Americas | North America | American Arctic | Greenland | Inuit |
| 225044 | NMNH | M | Unkw | Sunda-Pacific | South East Asia | Malay Archipelago | Indonesia | Java |
| 225046 | NMNH | M | Unkw | Sunda-Pacific | South East Asia | Malay Archipelago | Indonesia | Java |
| 225047 | NMNH | M | Unkw | Sunda-Pacific | South East Asia | Malay Archipelago | Indonesia | Java |
| 225053 | NMNH | F | Unkw | Sunda-Pacific | South East Asia | Malay Archipelago | Indonesia | Java |
| 225054 | NMNH | M | Unkw | Sunda-Pacific | South East Asia | Malay Archipelago | Indonesia | Java |
| 225055 | NMNH | F | Unkw | Sunda-Pacific | South East Asia | Malay Archipelago | Indonesia | Java |
| 225399 | NMNH | M | Unkw | Sunda-Pacific | South East Asia | Malay Archipelago | Indonesia | Java |
| 225120 | NMNH | NA | Unkw | Sunda-Pacific | South East Asia | Malay Archipelago | Indonesia | Java |
| 225007 | NMNH | NA | Unkw | Sahul-Pacific | Oceania | Melanesia | Papua New Guinea | New Britain |
| 226096 | NMNH | M | Unkw | Sahul-Pacific | Oceania | Melanesia | Papua New Guinea | New Britain |
| 226099 | NMNH | F | Unkw | Sahul-Pacific | Oceania | Melanesia | Papua New Guinea | New Britain |
| 226101 | NMNH | F | Unkw | Sahul-Pacific | Oceania | Melanesia | Papua New Guinea | New Britain |
| 226102 | NMNH | F | Unkw | Sahul-Pacific | Oceania | Melanesia | Papua New Guinea | New Britain |
| 226103 | NMNH | M | Unkw | Sahul-Pacific | Oceania | Melanesia | Papua New Guinea | New Britain |
| 226105 | NMNH | F | Unkw | Sahul-Pacific | Oceania | Melanesia | Papua New Guinea | New Britain |
| 226107 | NMNH | M | Unkw | Sahul-Pacific | Oceania | Melanesia | Papua New Guinea | New Britain |
| 226108 | NMNH | M | Unkw | Sahul-Pacific | Oceania | Melanesia | Papua New Guinea | New Britain |
| 226109 | NMNH | F | Unkw | Sahul-Pacific | Oceania | Melanesia | Papua New Guinea | New Britain |
| 226110 | NMNH | F | Unkw | Sahul-Pacific | Oceania | Melanesia | Papua New Guinea | New Britain |
| 226111 | NMNH | M | Unkw | Sahul-Pacific | Oceania | Melanesia | Papua New Guinea | New Britain |
| 226112 | NMNH | F | Unkw | Sahul-Pacific | Oceania | Melanesia | Papua New Guinea | New Britain |
| 226113 | NMNH | M | Unkw | Sahul-Pacific | Oceania | Melanesia | Papua New Guinea | New Britain |
| 226114 | NMNH | M | Unkw | Sahul-Pacific | Oceania | Melanesia | Papua New Guinea | New Britain |
| 226115 | NMNH | M | Unkw | Sahul-Pacific | Oceania | Melanesia | Papua New Guinea | New Britain |
| 226116 | NMNH | F | Unkw | Sahul-Pacific | Oceania | Melanesia | Papua New Guinea | New Britain |
| 226117 | NMNH | F | Unkw | Sahul-Pacific | Oceania | Melanesia | Papua New Guinea | New Britain |
| 226118 | NMNH | M | Unkw | Sahul-Pacific | Oceania | Melanesia | Papua New Guinea | New Britain |
| 227464 | NMNH | F | Unkw | Sahul-Pacific | Oceania | Melanesia | Papua New Guinea | New Britain |
| 381080 | NMNH | F | Unkw | Sahul-Pacific | Oceania | Melanesia | Papua New Guinea | New Britain |
| 381082 | NMNH | F | Unkw | Sahul-Pacific | Oceania | Melanesia | Papua New Guinea | New Britain |
| 381083 | NMNH | M | Unkw | Sahul-Pacific | Oceania | Melanesia | Papua New Guinea | New Britain |
| 205333 | NMNH | M | Unkw | Sahul-Pacific | Oceania | Polynesia | New Zealand | NA |
| 225113 | NMNH | F | Unkw | Sahul-Pacific | Oceania | Polynesia | New Zealand | NA |
| 225114 | NMNH | M | Unkw | Sahul-Pacific | Oceania | Polynesia | New Zealand | NA |
| 225115 | NMNH | M | Unkw | Sahul-Pacific | Oceania | Polynesia | New Zealand | NA |

| | | | | | | | | |
|---|---|---|---|---|---|---|---|---|
| 226140 | NMNH | F | Unkw | Sahul-Pacific | Oceania | Polynesia | New Zealand | NA |
| 226142 | NMNH | M | Unkw | Sahul-Pacific | Oceania | Polynesia | New Zealand | NA |
| 226145 | NMNH | M | Unkw | Sahul-Pacific | Oceania | Polynesia | New Zealand | NA |
| 226147 | NMNH | F | Unkw | Sahul-Pacific | Oceania | Polynesia | New Zealand | NA |
| 226150 | NMNH | M | Unkw | Sahul-Pacific | Oceania | Polynesia | New Zealand | NA |
| 226151 | NMNH | F | Unkw | Sahul-Pacific | Oceania | Polynesia | New Zealand | NA |
| 226152 | NMNH | F | Unkw | Sahul-Pacific | Oceania | Polynesia | New Zealand | NA |
| 226153 | NMNH | F | Unkw | Sahul-Pacific | Oceania | Polynesia | New Zealand | NA |
| 226154 | NMNH | F | Unkw | Sahul-Pacific | Oceania | Polynesia | New Zealand | NA |
| 226159 | NMNH | F | Unkw | Sahul-Pacific | Oceania | Polynesia | New Zealand | NA |
| 381086 | NMNH | F | Unkw | Sahul-Pacific | Oceania | Polynesia | New Zealand | NA |
| 381087 | NMNH | F | Unkw | Sahul-Pacific | Oceania | Polynesia | New Zealand | NA |
| 221998 | NMNH | F | Unkw | Sunda-Pacific | South East Asia | Malay Archipelago | Indonesia | Pagi Island |
| 221999 | NMNH | F | Unkw | Sunda-Pacific | South East Asia | Malay Archipelago | Indonesia | Pagi Island |
| 222000 | NMNH | F | Unkw | Sunda-Pacific | South East Asia | Malay Archipelago | Indonesia | Pagi Island |
| 222002 | NMNH | F | Unkw | Sunda-Pacific | South East Asia | Malay Archipelago | Indonesia | Pagi Island |
| 222003 | NMNH | M | Unkw | Sunda-Pacific | South East Asia | Malay Archipelago | Indonesia | Pagi Island |
| 276076 | NMNH | F | Unkw | Sahul-Pacific | Oceania | Melanesia | Papua New Guinea | NA |
| 276077 | NMNH | M | Unkw | Sahul-Pacific | Oceania | Melanesia | Papua New Guinea | NA |
| 276078 | NMNH | F | Unkw | Sahul-Pacific | Oceania | Melanesia | Papua New Guinea | NA |
| 276079 | NMNH | M | Unkw | Sahul-Pacific | Oceania | Melanesia | Papua New Guinea | NA |
| 276080 | NMNH | F | Unkw | Sahul-Pacific | Oceania | Melanesia | Papua New Guinea | NA |
| 225129 | NMNH | M | Unkw | Sunda-Pacific | South East Asia | Malay Archipelago | Philippines | NA |
| 259353 | NMNH | F | Unkw | Sunda-Pacific | South East Asia | Malay Archipelago | Philippines | NA |
| 227456 | NMNH | M | Unkw | Sahul-Pacific | Oceania | Melanesia | Solomon Islands | NA |
| 227457 | NMNH | F | Unkw | Sahul-Pacific | Oceania | Melanesia | Solomon Islands | NA |
| 227458 | NMNH | M | Unkw | Sahul-Pacific | Oceania | Melanesia | Solomon Islands | NA |
| ANI_23 | DW | F | <200 | Sunda-Pacific | South East Asia | Andaman Archipelago | Andaman Island | NA |
| ANI_16 | DW | F | <200 | Sunda-Pacific | South East Asia | Andaman Archipelago | Andaman Island | NA |
| ANI_35 | DW | F | <200 | Sunda-Pacific | South East Asia | Andaman Archipelago | Andaman Island | NA |
| ANI_37 | DW | M | <200 | Sunda-Pacific | South East Asia | Andaman Archipelago | Andaman Island | NA |
| ANI_39_40 | DW | F | <200 | Sunda-Pacific | South East Asia | Andaman Archipelago | Andaman Island | NA |
| ANI_32 | DW | F | <200 | Sunda-Pacific | South East Asia | Andaman Archipelago | Andaman Island | NA |
| ANI_27 | DW | F | <200 | Sunda-Pacific | South East Asia | Andaman Archipelago | Andaman Island | NA |
| ANI_36 | DW | M | <200 | Sunda-Pacific | South East Asia | Andaman Archipelago | Andaman Island | NA |
| ANI_29 | DW | F | <200 | Sunda-Pacific | South East Asia | Andaman Archipelago | Andaman Island | NA |
| ANI_31 | DW | M | <200 | Sunda-Pacific | South East Asia | Andaman Archipelago | Andaman Island | NA |
| ANI_33 | DW | M | <200 | Sunda-Pacific | South East Asia | Andaman Archipelago | Andaman Island | NA |
| ANI_26 | DW | F | <200 | Sunda-Pacific | South East Asia | Andaman Archipelago | Andaman Island | NA |
| ANI_28 | DW | M | <200 | Sunda-Pacific | South East Asia | Andaman Archipelago | Andaman Island | NA |
| ANI_30 | DW | F | <200 | Sunda-Pacific | South East Asia | Andaman Archipelago | Andaman Island | NA |
| ANI_17 | DW | F | <200 | Sunda-Pacific | South East Asia | Andaman Archipelago | Andaman Island | NA |
| ANI_07 | DW | F | <200 | Sunda-Pacific | South East Asia | Andaman Archipelago | Andaman Island | NA |
| ANI_11 | DW | M | <200 | Sunda-Pacific | South East Asia | Andaman Archipelago | Andaman Island | NA |

| | | | | | | | | |
|---|---|---|---|---|---|---|---|---|
| ANI_18 | DW | F | <200 | Sunda-Pacific | South East Asia | Andaman Archipelago | Andaman Island | NA |
| ANI_15 | DW | F | <200 | Sunda-Pacific | South East Asia | Andaman Archipelago | Andaman Island | NA |
| ANI_10 | DW | F | <200 | Sunda-Pacific | South East Asia | Andaman Archipelago | Andaman Island | NA |
| ANI_12 | DW | F | <200 | Sunda-Pacific | South East Asia | Andaman Archipelago | Andaman Island | NA |
| ANI_13 | DW | M | <200 | Sunda-Pacific | South East Asia | Andaman Archipelago | Andaman Island | NA |
| ANI_19 | DW | M | <200 | Sunda-Pacific | South East Asia | Andaman Archipelago | Andaman Island | NA |
| ANI_38 | DW | M | <200 | Sunda-Pacific | South East Asia | Andaman Archipelago | Andaman Island | NA |
| ANI_34 | DW | F | <200 | Sunda-Pacific | South East Asia | Andaman Archipelago | Andaman Island | NA |
| ANI_43 | DW | M | <200 | Sunda-Pacific | South East Asia | Andaman Archipelago | Nicobar Island | NA |
| ANI_42 | DW | M | <200 | Sunda-Pacific | South East Asia | Andaman Archipelago | Nicobar Island | NA |
| ANI_54 | DW | M | <200 | Sunda-Pacific | South East Asia | Andaman Archipelago | Nicobar Island | NA |
| ANI_41 | DW | M | <200 | Sunda-Pacific | South East Asia | Andaman Archipelago | Nicobar Island | NA |
| ANI_44 | DW | F | <200 | Sunda-Pacific | South East Asia | Andaman Archipelago | Nicobar Island | NA |
| ANI_59 | DW | M | <200 | Sunda-Pacific | South East Asia | Andaman Archipelago | Nicobar Island | NA |
| ANI_56 | DW | M | <200 | Sunda-Pacific | South East Asia | Andaman Archipelago | Nicobar Island | NA |
| ANI_50 | DW | M | <200 | Sunda-Pacific | South East Asia | Andaman Archipelago | Nicobar Island | NA |
| ANI_55 | DW | M | <200 | Sunda-Pacific | South East Asia | Andaman Archipelago | Nicobar Island | NA |
| ANI_61 | DW | M | <200 | Sunda-Pacific | South East Asia | Andaman Archipelago | Nicobar Island | NA |
| ANI_57 | DW | M | <200 | Sunda-Pacific | South East Asia | Andaman Archipelago | Nicobar Island | NA |
| ANI_58 | DW | M | <200 | Sunda-Pacific | South East Asia | Andaman Archipelago | Nicobar Island | NA |
| ANI_47 | DW | M | <200 | Sunda-Pacific | South East Asia | Andaman Archipelago | Nicobar Island | NA |
| ANI_49 | DW | M | <200 | Sunda-Pacific | South East Asia | Andaman Archipelago | Nicobar Island | NA |
| ANI_51 | DW | M | <200 | Sunda-Pacific | South East Asia | Andaman Archipelago | Nicobar Island | NA |
| ANI_48 | DW | M | <200 | Sunda-Pacific | South East Asia | Andaman Archipelago | Nicobar Island | NA |
| BU_28 | DW | F | <200 | Sunda-Pacific | South East Asia | Indochinese Peninsula | Myanmar | NA |
| BU_29 | DW | M | <200 | Sunda-Pacific | South East Asia | Indochinese Peninsula | Myanmar | NA |
| BU_21 | DW | M | <200 | Sunda-Pacific | South East Asia | Indochinese Peninsula | Myanmar | NA |
| BU_19 | DW | M | <200 | Sunda-Pacific | South East Asia | Indochinese Peninsula | Myanmar | NA |
| BU_04 | DW | M | <200 | Sunda-Pacific | South East Asia | Indochinese Peninsula | Myanmar | NA |
| BU_32 | DW | M | <200 | Sunda-Pacific | South East Asia | Indochinese Peninsula | Myanmar | NA |
| BU_14 | DW | M | <200 | Sunda-Pacific | South East Asia | Indochinese Peninsula | Myanmar | NA |
| BU_01 | DW | F | <200 | Sunda-Pacific | South East Asia | Indochinese Peninsula | Myanmar | NA |
| BU_31 | DW | M | <200 | Sunda-Pacific | South East Asia | Indochinese Peninsula | Myanmar | NA |
| BU_16 | DW | M | <200 | Sunda-Pacific | South East Asia | Indochinese Peninsula | Myanmar | NA |
| BU_10 | DW | M | <200 | Sunda-Pacific | South East Asia | Indochinese Peninsula | Myanmar | NA |
| MEL_120 | DW | M | <200 | Sahul-Pacific | Oceania | Melanesia | Papua New Guinea | Oriomo River Daudai |
| MEL_219 | DW | F | <200 | Sahul-Pacific | Oceania | Melanesia | Papua New Guinea | Murua Island Muyuw |
| MEL_264 | DW | M | <200 | Sahul-Pacific | Oceania | Melanesia | Papua New Guinea | NA |

| | | | | | | | | |
|---|---|---|---|---|---|---|---|---|
| MEL_104 | DW | F | <200 | Sahul-Pacific | Oceania | Melanesia | Papua New Guinea | Kwaiawata Island Muyuw |
| MEL_197 | DW | M | <200 | Sahul-Pacific | Oceania | Melanesia | Papua New Guinea | Murua Island Muyuw |
| MEL_258 | DW | M | <200 | Sahul-Pacific | Oceania | Melanesia | Papua New Guinea | Kwaiawata Island Muyuw |
| MEL_259 | DW | M | <200 | Sahul-Pacific | Oceania | Melanesia | Papua New Guinea | Kwaiawata Island Muyuw |
| MEL_272 | DW | M | <200 | Sahul-Pacific | Oceania | Melanesia | Papua New Guinea | Kwaiawata Island Muyuw |
| MEL_273 | DW | M | <200 | Sahul-Pacific | Oceania | Melanesia | Papua New Guinea | Kwaiawata Island Muyuw |
| MEL_189 | DW | M | <200 | Sahul-Pacific | Oceania | Melanesia | Papua New Guinea | Murua Island Muyuw |
| SAS_13 | DW | F | <200 | West Eurasia | South Asia | Indian Sub-Continent | North India | Punjab |
| SAS_19 | DW | M | <200 | West Eurasia | South Asia | Indian Sub-Continent | Pakistan | NA |
| SAS_16 | DW | M | <200 | West Eurasia | South Asia | Indian Sub-Continent | Pakistan | Pathan |
| SAS_44 | DW | F | <200 | West Eurasia | South Asia | Indian Sub-Continent | South India | Deccan Berars |
| SAS_45 | DW | M | <200 | West Eurasia | South Asia | Indian Sub-Continent | East India | Patna |
| SAS_08 | DW | F | <200 | West Eurasia | South Asia | Indian Sub-Continent | North India | Naharhmpikya Sinhalese |
| SAS_29 | DW | M | <200 | West Eurasia | South Asia | Indian Sub-Continent | North India | Punjab |
| SAS_23 | DW | F | <200 | West Eurasia | South Asia | Indian Sub-Continent | North India | Punjab |
| SAS_07 | DW | M | <200 | West Eurasia | South Asia | Indian Sub-Continent | North India | NA |
| SAS_04 | DW | F | <200 | West Eurasia | South Asia | Indian Sub-Continent | North India | Veddah |
| SAS_31 | DW | F | <200 | West Eurasia | South Asia | Indian Sub-Continent | North India | Punjab |
| SAS_81 | DW | F | <200 | West Eurasia | South Asia | Indian Sub-Continent | India | NA |
| SAS_57 | DW | F | <200 | West Eurasia | South Asia | Indian Sub-Continent | East India | Hindustan Bihar |
| SAS_10 | DW | M | <200 | West Eurasia | South Asia | Indian Sub-Continent | Sri Lanka | Colombo |
| SAS_17 | DW | F | <200 | West Eurasia | South Asia | Indian Sub-Continent | Pakistan | Pathan |
| SAS_15 | DW | M | <200 | West Eurasia | South Asia | Indian Sub-Continent | Pakistan | Pathan |
| SAS_27 | DW | F | <200 | West Eurasia | South Asia | Indian Sub-Continent | North India | Punjab |
| SAS_20 | DW | F | <200 | West Eurasia | South Asia | Indian Sub-Continent | North India | Punjab |
| SAS_03 | DW | F | <200 | West Eurasia | South Asia | Indian Sub-Continent | North India | Veddah |
| SAS_60 | DW | M | <200 | West Eurasia | South Asia | Indian Sub-Continent | North India | NA |
| SAS_54 | DW | M | <200 | West Eurasia | South Asia | Indian Sub-Continent | North India | Hindustan |
| SAS_71 | DW | F | <200 | West Eurasia | South Asia | Indian Sub-Continent | Bangladesh | Bengal |
| SAS_46 | DW | M | <200 | West Eurasia | South Asia | Indian Sub-Continent | East India | Bihari |
| SAS_61 | DW | M | <200 | West Eurasia | South Asia | Indian Sub-Continent | North India | Pakistan |
| SAS_39 | DW | F | <200 | West Eurasia | South Asia | Indian Sub-Continent | East India | Bengal |
| SAS_35 | DW | M | <200 | West Eurasia | South Asia | Indian Sub-Continent | East India | Bengal |
| SAS_47 | DW | F | <200 | West Eurasia | South Asia | Indian Sub-Continent | East India | Bihari |
| SAS_02 | DW | M | <200 | West Eurasia | South Asia | Indian Sub-Continent | North India | Veddah |
| SAS_51 | DW | M | <200 | West Eurasia | South Asia | Indian Sub-Continent | South India | Coorg |
| SAS_40 | DW | M | <200 | West Eurasia | South Asia | Indian Sub-Continent | East India | Bengal |
| SAS_84 | DW | F | <200 | West Eurasia | South Asia | Indian Sub-Continent | India | NA |

| | | | | | | | | |
|---|---|---|---|---|---|---|---|---|
| SAS_34 | DW | F | <200 | West Eurasia | South Asia | Indian Sub-Continent | East India | Bengal |
| SAS_01 | DW | F | <200 | West Eurasia | South Asia | Indian Sub-Continent | North India | Veddah |
| SAS_09 | DW | M | <200 | West Eurasia | South Asia | Indian Sub-Continent | Sri Lanka | Colombo |
| SAS_69 | DW | M | <200 | West Eurasia | South Asia | Indian Sub-Continent | Bangladesh | Bengal |
| SAS_83 | DW | M | <200 | West Eurasia | South Asia | Indian Sub-Continent | India | Hindu |
| SAS_77 | DW | M | <200 | West Eurasia | South Asia | Indian Sub-Continent | South India | Dravidian |
| SAS_56 | DW | F | <200 | West Eurasia | South Asia | Indian Sub-Continent | West India | Mumbai Parsi |
| SAS_38 | DW | M | <200 | West Eurasia | South Asia | Indian Sub-Continent | East India | Bengal |
| SAS_53 | DW | F | <200 | West Eurasia | South Asia | Indian Sub-Continent | Sri Lanka | Ballam Coffa |
| SAS_78 | DW | M | <200 | West Eurasia | South Asia | Indian Sub-Continent | South India | NA |
| SAS_36 | DW | M | <200 | West Eurasia | South Asia | Indian Sub-Continent | East India | Bengal |
| SAS_42 | DW | M | <200 | West Eurasia | South Asia | Indian Sub-Continent | India Unkw | Ballam Coffa |
| SAS_11 | DW | M | <200 | West Eurasia | South Asia | Indian Sub-Continent | North India | NA |
| SAS_55 | DW | M | <200 | West Eurasia | South Asia | Indian Sub-Continent | Sri Lanka | Eingenadu |
| SAS_67 | DW | F | <200 | West Eurasia | South Asia | Indian Sub-Continent | India Unkw | NA |
| SAS_70 | DW | F | <200 | West Eurasia | South Asia | Indian Sub-Continent | Bangladesh | Bengal |
| SAS_75 | DW | F | <200 | West Eurasia | South Asia | Indian Sub-Continent | India Unkw | NA |
| SAS_79 | DW | F | <200 | West Eurasia | South Asia | Indian Sub-Continent | India | Hindu |
| SAS_37 | DW | F | <200 | West Eurasia | South Asia | Indian Sub-Continent | East India | Bengal Bangladesh |
| SAS_28 | DW | F | <200 | West Eurasia | South Asia | Indian Sub-Continent | North India | Punjab |
| SAS_26 | DW | M | <200 | West Eurasia | South Asia | Indian Sub-Continent | North India | Punjab |
| SAS_30 | DW | M | <200 | West Eurasia | South Asia | Indian Sub-Continent | North India | Punjab |
| SAS_52 | DW | M | <200 | West Eurasia | South Asia | Indian Sub-Continent | South India | Paliyan Tribe |
| SAS_25 | DW | M | <200 | West Eurasia | South Asia | Indian Sub-Continent | North India | Punjab |
| SAS_24 | DW | M | <200 | West Eurasia | South Asia | Indian Sub-Continent | North India | Punjab |
| SAS_21 | DW | F | <200 | West Eurasia | South Asia | Indian Sub-Continent | North India | Punjab |
| SAS_33 | DW | M | <200 | West Eurasia | South Asia | Indian Sub-Continent | East India | Bengal |
| SAS_05 | DW | M | <200 | West Eurasia | South Asia | Indian Sub-Continent | North India | Veddah |
| SAS_62 | DW | M | <200 | West Eurasia | South Asia | Indian Sub-Continent | East India | NA |
| SAS_48 | DW | M | <200 | West Eurasia | South Asia | Indian Sub-Continent | East India | Hindustan Bihar |
| SAS_68 | DW | F | <200 | West Eurasia | South Asia | Indian Sub-Continent | East India | Bengal Bangladesh |
| 5423 | DW | F | <200 | Sub-Saharan Africa | Sub-Saharan Africa | Western Africa | Nigeria | Kagoro |
| 6087 | DW | F | <200 | Sub-Saharan Africa | Sub-Saharan Africa | Western Africa | Nigeria | Kaduna |
| 1734 | DW | M | <200 | Sub-Saharan Africa | Sub-Saharan Africa | Southern Africa | South Africa | Basuto |
| 1743 | DW | F | <200 | Sub-Saharan Africa | Sub-Saharan Africa | Southern Africa | South Africa | Port Elizabeth |
| 5340 | DW | F | <200 | Sub-Saharan Africa | Sub-Saharan Africa | Eastern Africa | Kenya | Akamba |
| AF1082 | DW | F | <200 | Sub-Saharan Africa | Sub-Saharan Africa | Eastern Africa | Kenya | NA |
| 1755 | DW | F | <200 | Sub-Saharan Africa | Sub-Saharan Africa | Southern Africa | South Africa | Wynberg San |

| | | | | | | | | |
|---|---|---|---|---|---|---|---|---|
| 3731 | DW | F | <200 | Sub-Saharan Africa | Sub-Saharan Africa | Southern Africa | South Africa | Knysna Cave |
| 1728 | DW | M | <200 | Sub-Saharan Africa | Sub-Saharan Africa | Western Africa | Guinea | NA |
| 5585 | DW | F | <200 | Sub-Saharan Africa | Sub-Saharan Africa | Southern Africa | Namibia | Walvis Bay |
| 5651 | DW | M | <200 | Sub-Saharan Africa | Sub-Saharan Africa | Western Africa | Nigeria | Muri Province |
| 1735 | DW | M | <200 | Sub-Saharan Africa | Sub-Saharan Africa | Southern Africa | South Africa | Manatee Cradock |
| 5425 | DW | M | <200 | Sub-Saharan Africa | Sub-Saharan Africa | Western Africa | Nigeria | Gannawarri |
| 1747 | DW | M | <200 | Sub-Saharan Africa | Sub-Saharan Africa | Southern Africa | South Africa | Korana |
| 5060 | DW | M | <200 | Sub-Saharan Africa | Sub-Saharan Africa | Central Africa | Congo | Brazaville |
| 4_93 | DW | M | <200 | Sub-Saharan Africa | Sub-Saharan Africa | Western Africa | Nigeria | NA |
| 1733 | DW | M | <200 | Sub-Saharan Africa | Sub-Saharan Africa | Southern Africa | South Africa | Amaponda |
| 6097 | DW | F | <200 | Sub-Saharan Africa | Sub-Saharan Africa | Western Africa | Nigeria | Yola |
| AF_35_0_1 | DW | M | <200 | Sub-Saharan Africa | Sub-Saharan Africa | Southern Africa | South Africa | NA |
| 1731 | DW | M | <200 | Sub-Saharan Africa | Sub-Saharan Africa | Southern Africa | Angola | Luanda |
| 4697 | DW | F | <200 | Sub-Saharan Africa | Sub-Saharan Africa | Western Africa | Nigeria | NA |
| 6110 | DW | F | <200 | Sub-Saharan Africa | Sub-Saharan Africa | Southern Africa | South Africa | Bechuanaland |
| 1732 | DW | M | <200 | Sub-Saharan Africa | Sub-Saharan Africa | Southern Africa | SSA Unkw | NA |
| 6109 | DW | F | <200 | Sub-Saharan Africa | Sub-Saharan Africa | Southern Africa | South Africa | Bechuanaland |
| 5424 | DW | M | <200 | Sub-Saharan Africa | Sub-Saharan Africa | Western Africa | Nigeria | Kagoro |
| 6093 | DW | M | <200 | Sub-Saharan Africa | Sub-Saharan Africa | Eastern Africa | Uganda | Teso |
| 4197 | DW | M | <200 | Sub-Saharan Africa | Sub-Saharan Africa | Southern Africa | South Africa | NA |
| 6089 | DW | F | <200 | Sub-Saharan Africa | Sub-Saharan Africa | Eastern Africa | Nigeria | Kaduna |
| AF_0_1 | DW | F | <200 | Sub-Saharan Africa | Sub-Saharan Africa | Eastern Africa | Mozambique | Makua |
| 1721 | DW | M | <1000 | West Eurasia | North Africa | Northern Africa | Canary Islands | Guanche |
| 1769 | DW | M | <200 | West Eurasia | North Africa | Northern Africa | Egypt | NA |
| 1777 | DW | F | <200 | Sub-Saharan Africa | Sub-Saharan Africa | Central Africa | Congo | Upper Congo River |
| 1751 | DW | M | <200 | Sub-Saharan Africa | Sub-Saharan Africa | Southern Africa | South Africa | Kalahari |
| 5058 | DW | M | <200 | Sub-Saharan Africa | Sub-Saharan Africa | Central Africa | Congo | Bambuti Pygmy |
| 5643 | DW | M | <200 | Sub-Saharan Africa | Sub-Saharan Africa | Western Africa | Ghana | Ashanti |
| 1749 | DW | F | <200 | Sub-Saharan Africa | Sub-Saharan Africa | Southern Africa | South Africa | NA |
| 1729 | DW | F | <200 | Sub-Saharan Africa | Sub-Saharan Africa | Central Africa | Congo | NA |
| 1709 | DW | M | <1000 | West Eurasia | North Africa | Northern Africa | Canary Islands | Guanche |
| 1722 | DW | F | <1000 | West Eurasia | North Africa | Northern Africa | Canary Islands | Guanche |
| 6094 | DW | M | <200 | Sub-Saharan Africa | Sub-Saharan Africa | Eastern Africa | Uganda | Teso |
| 1711 | DW | M | <1000 | West Eurasia | North Africa | Northern Africa | Canary Islands | Guanche |
| 3732 | DW | F | <200 | Sub-Saharan Africa | Sub-Saharan Africa | Southern Africa | South Africa | Knysna Cave |
| 6092 | DW | M | <200 | Sub-Saharan Africa | Sub-Saharan Africa | Eastern Africa | Uganda | NA |
| 1737 | DW | M | <200 | Sub-Saharan Africa | Sub-Saharan Africa | Southern Africa | South Africa | Amaxhosa Great Winterberg |

| | | | | | | | | |
|---|---|---|---|---|---|---|---|---|
| 1727 | DW | F | <200 | Sub-Saharan Africa | Sub-Saharan Africa | Eastern Africa | Tanzania | Makua |
| 1774 | DW | M | <200 | Sub-Saharan Africa | Sub-Saharan Africa | Eastern Africa | Tanzania | NA |
| 4696 | DW | M | <200 | Sub-Saharan Africa | Sub-Saharan Africa | Western Africa | Nigeria | NA |
| Af_31_0_1 | DW | M | <200 | Sub-Saharan Africa | Sub-Saharan Africa | Eastern Africa | Zimbabwe | NA |
| 1730 | DW | M | <200 | Sub-Saharan Africa | Sub-Saharan Africa | Western Africa | Guinea | NA |
| 1739 | DW | M | <200 | Sub-Saharan Africa | Sub-Saharan Africa | Southern Africa | South Africa | Khoikhoi |
| 6096 | DW | M | <200 | Sub-Saharan Africa | Sub-Saharan Africa | Western Africa | Nigeria | Yola |
| 1744 | DW | M | <200 | Sub-Saharan Africa | Sub-Saharan Africa | Southern Africa | South Africa | Knysna Cave |
| 149 | DW | M | <200 | Sub-Saharan Africa | Sub-Saharan Africa | Southern Africa | South Africa | NA |
| Af_44_0_2 | DW | M | <200 | Sub-Saharan Africa | Sub-Saharan Africa | Western Africa | Nigeria | Yoruba Ilorin |
| 1738 | DW | F | <200 | Sub-Saharan Africa | Sub-Saharan Africa | Southern Africa | South Africa | NA |
| Af_20_0_1 | DW | M | <200 | Sub-Saharan Africa | Sub-Saharan Africa | SSA Unkw | SSA Unkw | Bantu Kaoisoudo |
| 6085 | DW | M | <200 | Sub-Saharan Africa | Sub-Saharan Africa | Eastern Africa | Kenya | NA |
| 1778 | DW | F | <200 | Sub-Saharan Africa | Sub-Saharan Africa | Central Africa | Congo | NA |
| AF_30_0_1 | DW | M | <200 | Sub-Saharan Africa | Sub-Saharan Africa | Southern Africa | South Africa | NA |
| 5418 | DW | F | <200 | Sub-Saharan Africa | Sub-Saharan Africa | Eastern Africa | Kenya | Kikuyu |
| AF_44_0_4 | DW | F | <200 | Sub-Saharan Africa | Sub-Saharan Africa | Western Africa | Nigeria | Yoruba Ilorin |
| 1725 | DW | M | 200 | Sub-Saharan Africa | Sub-Saharan Africa | Western Africa | Ghana | Fanti |
| 5428 | DW | F | <200 | Sub-Saharan Africa | Sub-Saharan Africa | Western Africa | Nigeria | Kagoro |
| 5701 | DW | M | <200 | Sub-Saharan Africa | Sub-Saharan Africa | Western Africa | Nigeria | Muri Province |
| AF_15_0_6 | DW | F | <200 | Sub-Saharan Africa | Sub-Saharan Africa | Eastern Africa | Somalia | Jilili |
| AF_15_0_27 | DW | M | <200 | Sub-Saharan Africa | Sub-Saharan Africa | Eastern Africa | Somalia | Darood |
| AF_15_0_30 | DW | F | <200 | Sub-Saharan Africa | Sub-Saharan Africa | Eastern Africa | Somalia | NA |
| AF_15_0_22 | DW | M | <200 | Sub-Saharan Africa | Sub-Saharan Africa | Eastern Africa | Somalia | Darood |
| AF_15_0_34 | DW | M | <200 | Sub-Saharan Africa | Sub-Saharan Africa | Eastern Africa | Somalia | Darood Hawiya |
| AF_15_0_17 | DW | M | <200 | Sub-Saharan Africa | Sub-Saharan Africa | Eastern Africa | Somalia | Darood |
| AF_15_0_50 | DW | M | <200 | Sub-Saharan Africa | Sub-Saharan Africa | Eastern Africa | Somalia | Darood Hawiya |
| AF_15_0_48 | DW | M | <200 | Sub-Saharan Africa | Sub-Saharan Africa | Eastern Africa | Somalia | Darood Hawiya |
| AF_15_0_19 | DW | M | <200 | Sub-Saharan Africa | Sub-Saharan Africa | Eastern Africa | Somalia | Darood |
| AF_15_0_11 | DW | M | <200 | Sub-Saharan Africa | Sub-Saharan Africa | Eastern Africa | Somalia | Tegera Well |
| AF_15_0_23 | DW | M | <200 | Sub-Saharan Africa | Sub-Saharan Africa | Eastern Africa | Somalia | Darood |
| AF_15_0_70 | DW | M | <200 | Sub-Saharan Africa | Sub-Saharan Africa | Eastern Africa | Somalia | Darood Hawiya |
| AF_15_0_62 | DW | M | <200 | Sub-Saharan Africa | Sub-Saharan Africa | Eastern Africa | Somalia | NA |
| AF_15_0_5 | DW | M | <200 | Sub-Saharan Africa | Sub-Saharan Africa | Eastern Africa | Somalia | Didali |
| AF_15_0_55 | DW | M | <200 | Sub-Saharan Africa | Sub-Saharan Africa | Eastern Africa | Somalia | Darood Hawiya |
| AF_15_0_53 | DW | M | <200 | Sub-Saharan Africa | Sub-Saharan Africa | Eastern Africa | Somalia | Darood Hawiya |
| AF_15_0_12 | DW | M | <200 | Sub-Saharan Africa | Sub-Saharan Africa | Eastern Africa | Somalia | Ainaho, Hahr Jalo or Daldshantu tribe |

| | | | | | | | | |
|---|---|---|---|---|---|---|---|---|
| AF_15_0_18 | DW | M | <200 | Sub-Saharan Africa | Sub-Saharan Africa | Eastern Africa | Somalia | Darood |
| AF_15_0_69 | DW | M | <200 | Sub-Saharan Africa | Sub-Saharan Africa | Eastern Africa | Somalia | Darood Hawiya |
| AF_15_0_1 | DW | M | <200 | Sub-Saharan Africa | Sub-Saharan Africa | Eastern Africa | Somalia | Ali Kush |
| AF_15_0_41 | DW | M | <200 | Sub-Saharan Africa | Sub-Saharan Africa | Eastern Africa | Somalia | Darood Hawiya |
| AF_15_0_64 | DW | M | <200 | Sub-Saharan Africa | Sub-Saharan Africa | Eastern Africa | Somalia | Darood Hawiya |
| AF_15_0_32 | DW | M | <200 | Sub-Saharan Africa | Sub-Saharan Africa | Eastern Africa | Somalia | Darood Hawiya |
| AF_15_0_42 | DW | M | <200 | Sub-Saharan Africa | Sub-Saharan Africa | Eastern Africa | Somalia | Darood Hawiya |
| AF_15_0_65 | DW | M | <200 | Sub-Saharan Africa | Sub-Saharan Africa | Eastern Africa | Somalia | Darood Hawiya |
| AF_15_0_02 | DW | M | <200 | Sub-Saharan Africa | Sub-Saharan Africa | Eastern Africa | Somalia | Hariya |
| AF_15_0_31 | DW | M | <200 | Sub-Saharan Africa | Sub-Saharan Africa | Eastern Africa | Somalia | NA |
| AF_15_0_25 | DW | M | <200 | Sub-Saharan Africa | Sub-Saharan Africa | Eastern Africa | Somalia | Darood |
| AF_15_0_16 | DW | M | <200 | Sub-Saharan Africa | Sub-Saharan Africa | Eastern Africa | Somalia | Ainaho, Burao |
| AF_15_0_13 | DW | F | <200 | Sub-Saharan Africa | Sub-Saharan Africa | Eastern Africa | Somalia | Ainaho, Burao |
| AF_15_0_03 | DW | M | <200 | Sub-Saharan Africa | Sub-Saharan Africa | Eastern Africa | Somalia | Hadad |
| AF_15_0_47 | DW | M | <200 | Sub-Saharan Africa | Sub-Saharan Africa | Eastern Africa | Somalia | Darood Hawiya |
| AF_15_0_67 | DW | M | <200 | Sub-Saharan Africa | Sub-Saharan Africa | Eastern Africa | Somalia | Darood Hawiya |
| AF_15_0_33 | DW | M | <200 | Sub-Saharan Africa | Sub-Saharan Africa | Eastern Africa | Somalia | Darood Hawiya |
| AF_23_0_22 | DW | M | <200 | Sub-Saharan Africa | Sub-Saharan Africa | Eastern Africa | Tanzania | Haya |
| AF_23_0_30 | DW | F | <200 | Sub-Saharan Africa | Sub-Saharan Africa | Eastern Africa | Tanzania | Haya |
| AF_23_0_19 | DW | M | <200 | Sub-Saharan Africa | Sub-Saharan Africa | Eastern Africa | Tanzania | Haya |
| AF_23_0_23 | DW | M | <200 | Sub-Saharan Africa | Sub-Saharan Africa | Eastern Africa | Tanzania | Haya |
| AF_23_0_32 | DW | M | <200 | Sub-Saharan Africa | Sub-Saharan Africa | Eastern Africa | Tanzania | Haya |
| AF_23_0_112 | DW | F | <200 | Sub-Saharan Africa | Sub-Saharan Africa | Eastern Africa | Tanzania | Haya |
| AF_23_0_113 | DW | M | <200 | Sub-Saharan Africa | Sub-Saharan Africa | Eastern Africa | Tanzania | Haya |
| AF_23_0_17 | DW | M | <200 | Sub-Saharan Africa | Sub-Saharan Africa | Eastern Africa | Tanzania | Haya |
| AF_23_0_31 | DW | M | <200 | Sub-Saharan Africa | Sub-Saharan Africa | Eastern Africa | Tanzania | Haya |
| AF_23_0_109 | DW | M | <200 | Sub-Saharan Africa | Sub-Saharan Africa | Eastern Africa | Tanzania | Haya |
| AF_23_0_04 | DW | M | <200 | Sub-Saharan Africa | Sub-Saharan Africa | Eastern Africa | Tanzania | Haya |
| AF_23_0_20 | DW | F | <200 | Sub-Saharan Africa | Sub-Saharan Africa | Eastern Africa | Tanzania | Haya |
| AF_23_0_219 | DW | F | <200 | Sub-Saharan Africa | Sub-Saharan Africa | Eastern Africa | Tanzania | Haya |
| AF_23_0_223 | DW | M | <200 | Sub-Saharan Africa | Sub-Saharan Africa | Eastern Africa | Tanzania | Haya |
| AF_23_0_228 | DW | F | <200 | Sub-Saharan Africa | Sub-Saharan Africa | Eastern Africa | Tanzania | Haya |
| AF_23_0_224 | DW | F | <200 | Sub-Saharan Africa | Sub-Saharan Africa | Eastern Africa | Tanzania | Haya |
| AF_23_0_39 | DW | M | <200 | Sub-Saharan Africa | Sub-Saharan Africa | Eastern Africa | Tanzania | Haya |
| AF_23_0_27 | DW | F | <200 | Sub-Saharan Africa | Sub-Saharan Africa | Eastern Africa | Tanzania | Haya |
| AF_23_0_37 | DW | M | <200 | Sub-Saharan Africa | Sub-Saharan Africa | Eastern Africa | Tanzania | Haya |
| AF_23_0_111 | DW | M | <200 | Sub-Saharan Africa | Sub-Saharan Africa | Eastern Africa | Tanzania | Haya |
| AF_23_0_21_ | DW | F | <200 | Sub-Saharan Africa | Sub-Saharan Africa | Eastern Africa | Tanzania | Haya |

| | | | | | | | | |
|---|---|---|---|---|---|---|---|---|
| AF_23_0_209 | DW | F | <200 | Sub-Saharan Africa | Sub-Saharan Africa | Eastern Africa | Tanzania | Haya |
| AF_23_0_35 | DW | M | <200 | Sub-Saharan Africa | Sub-Saharan Africa | Eastern Africa | Tanzania | Haya |
| AF_23_0_118 | DW | M | <200 | Sub-Saharan Africa | Sub-Saharan Africa | Eastern Africa | Tanzania | Haya |
| AF_23_0_115 | DW | M | <200 | Sub-Saharan Africa | Sub-Saharan Africa | Eastern Africa | Tanzania | Haya |
| AF_23_0_225 | DW | F | <200 | Sub-Saharan Africa | Sub-Saharan Africa | Eastern Africa | Tanzania | Haya |
| AF_23_0_119 | DW | M | <200 | Sub-Saharan Africa | Sub-Saharan Africa | Eastern Africa | Tanzania | Haya |
| AF_23_0_110 | DW | M | <200 | Sub-Saharan Africa | Sub-Saharan Africa | Eastern Africa | Tanzania | Haya |
| AF_23_0_44 | DW | F | <200 | Sub-Saharan Africa | Sub-Saharan Africa | Eastern Africa | Tanzania | Haya |
| AF_23_0_38 | DW | M | <200 | Sub-Saharan Africa | Sub-Saharan Africa | Eastern Africa | Tanzania | Haya |
| AF_23_0_213 | DW | M | <200 | Sub-Saharan Africa | Sub-Saharan Africa | Eastern Africa | Tanzania | Haya |
| AF_23_0_36 | DW | M | <200 | Sub-Saharan Africa | Sub-Saharan Africa | Eastern Africa | Tanzania | Haya |
| AF_23_0_218 | DW | M | <200 | Sub-Saharan Africa | Sub-Saharan Africa | Eastern Africa | Tanzania | Haya |
| AF_23_0_25 | DW | F | <200 | Sub-Saharan Africa | Sub-Saharan Africa | Eastern Africa | Tanzania | Haya |
| AF_23_0_42 | DW | F | <200 | Sub-Saharan Africa | Sub-Saharan Africa | Eastern Africa | Tanzania | Haya |
| AF_23_0_114 | DW | M | <200 | Sub-Saharan Africa | Sub-Saharan Africa | Eastern Africa | Tanzania | Haya |
| AF_23_0_200 | DW | M | <200 | Sub-Saharan Africa | Sub-Saharan Africa | Eastern Africa | Tanzania | Haya |
| AF_23_0_18 | DW | M | <200 | Sub-Saharan Africa | Sub-Saharan Africa | Eastern Africa | Tanzania | Haya |
| AF_23_0_202 | DW | M | <200 | Sub-Saharan Africa | Sub-Saharan Africa | Eastern Africa | Tanzania | Haya |
| AF_23_0_116 | DW | M | <200 | Sub-Saharan Africa | Sub-Saharan Africa | Eastern Africa | Tanzania | Haya |
| AF_23_0_227 | DW | F | <200 | Sub-Saharan Africa | Sub-Saharan Africa | Eastern Africa | Tanzania | Haya |
| AF_23_0_226 | DW | M | <200 | Sub-Saharan Africa | Sub-Saharan Africa | Eastern Africa | Tanzania | Haya |
| AF_23_0_34 | DW | F | <200 | Sub-Saharan Africa | Sub-Saharan Africa | Eastern Africa | Tanzania | Haya |
| AF_23_0_8 | DW | F | <200 | Sub-Saharan Africa | Sub-Saharan Africa | Eastern Africa | Tanzania | Haya |
| AF_23_0_16 | DW | M | <200 | Sub-Saharan Africa | Sub-Saharan Africa | Eastern Africa | Tanzania | Haya |
| AF_23_0_2 | DW | M | <200 | Sub-Saharan Africa | Sub-Saharan Africa | Eastern Africa | Tanzania | Haya |
| 1067 | DW | M | <200 | West Eurasia | Europe | Western Europe | France | NA |
| 1065 | DW | M | <200 | West Eurasia | Europe | Central Europe | Switzerland | St Bernard |
| Eu_25_00_2 | DW | F | <200 | West Eurasia | Europe | Central Europe | Switzerland | Graubunden Saint Moritz |
| Eu_25_00_1 | DW | M | <200 | West Eurasia | Europe | Central Europe | Switzerland | Graubunden Saint Moritz |
| 1036 | DW | M | <200 | West Eurasia | Europe | Western Europe | France | NA |
| 1181 | DW | M | <200 | West Eurasia | Europe | Central Europe | Austrian | NA |
| 1178 | DW | M | <200 | West Eurasia | Europe | Northern Europe | Sweden | NA |
| 2235 | DW | M | <200 | West Eurasia | Europe | Western Europe | France | Paris |
| 3000 | DW | M | <200 | West Eurasia | Europe | Central Europe | Hungary | NA |
| 1143 | DW | M | <200 | West Eurasia | Europe | Central Europe | Austrian | Vienna |
| Eu_26_00_2 | DW | M | <200 | West Eurasia | Europe | Central Europe | Germany | Halle |
| 1150 | DW | F | <200 | West Eurasia | Europe | Central Europe | Austrian | NA |
| Eu_31_0_1 | DW | M | <200 | West Eurasia | Europe | Eastern Europe | Ukraine | Crime Sebastopol |

| | | | | | | | | |
|---|---|---|---|---|---|---|---|---|
| 1051 | DW | M | <200 | West Eurasia | Europe | Western Europe | France | Brittany |
| 1042 | DW | M | <200 | West Eurasia | Europe | Western Europe | France | NA |
| Eu_26_00_1 | DW | M | <200 | West Eurasia | Europe | Central Europe | Germany | Halle |
| Eu_31_00_1 | DW | M | <200 | West Eurasia | Europe | Eastern Europe | Russia | Khanty Kondinski |
| 1155 | DW | M | <200 | West Eurasia | Europe | Central Europe | Czechoslovakia | NA |
| Eu_31_00_2 | DW | F | <200 | West Eurasia | Europe | Eastern Europe | Russia | Salekhard |
| 1173 | DW | F | <200 | West Eurasia | Europe | Northern Europe | Finland | Lapland |
| Eu_42_00_1 | DW | F | <200 | West Eurasia | Europe | Southern Europe | Italy | NA |
| Eu_34_4_1 | DW | F | <200 | West Eurasia | Europe | Central Europe | Hungary | Toszeg |
| Eu_24_00_2 | DW | F | <200 | West Eurasia | Europe | Western Europe | France | NA |
| Eu_45_4_1 | DW | M | <200 | West Eurasia | Europe | Southern Europe | Spain | Minorca |
| Eu_44_0_3 | DW | M | <200 | West Eurasia | Europe | Southern Europe | Italy | Sardinia |
| 1114 | DW | M | <200 | West Eurasia | Europe | Southern Europe | Italy | Paestum |
| Eu_42_00_5 | DW | F | <200 | West Eurasia | Europe | Southern Europe | Italy | Lazio |
| Eu_44_00_1 | DW | F | <200 | West Eurasia | Europe | Southern Europe | Italy | Sardinia |
| Eu_24_00_1 | DW | M | <200 | West Eurasia | Europe | Western Europe | France | NA |
| Eu_1_5_67 | DW | M | <200 | West Eurasia | Europe | Western Europe | England | NA |
| 1121 | DW | M | <200 | West Eurasia | Europe | Southern Europe | Italy | Rome |
| Eu_43_00_4 | DW | F | <200 | West Eurasia | Europe | Southern Europe | Malta | Bingemma |
| Eu_1_5_82 | DW | M | <200 | West Eurasia | Europe | Western Europe | England | South Wilshire |
| Eu_45_4_2 | DW | F | <200 | West Eurasia | Europe | Southern Europe | Spain | Minorca |
| Eu_45_4_3 | DW | F | <200 | West Eurasia | Europe | Southern Europe | Spain | Minorca |
| 1118 | DW | F | <200 | West Eurasia | Europe | Southern Europe | Italy | Sardinia |
| Eu_42_00_2 | DW | F | <200 | West Eurasia | Europe | Southern Europe | Italy | NA |
| Eu_43_00_3 | DW | M | <200 | West Eurasia | Europe | Southern Europe | Malta | Tal Horr |
| 5903 | DW | M | <200 | West Eurasia | Europe | Southern Europe | Greece | Thessaly |
| 1120 | DW | M | <200 | West Eurasia | Europe | Southern Europe | Italy | Rome |
| 6044 | DW | M | <200 | West Eurasia | Europe | Southern Europe | Italy | Sicily |
| 1898 | DW | M | <200 | Sino-Americas | South America | Andean | Argentina | Rio Gallegos |
| CA019 | DW | F | <200 | Sino-Americas | North America | Carribbean | Jamaica | NA |
| CA004 | DW | F | <200 | Sino-Americas | Central America | Central America | Guatemala | Gondaiaio |
| CA025 | DW | M | <200 | Sino-Americas | North America | Carribbean | Jamaica | NA |
| CA017 | DW | M | <200 | Sino-Americas | North America | Carribbean | Jamaica | NA |
| NA023 | DW | F | <200 | Sino-Americas | North America | NA Northeast Woodlands | United States | Iroquois |
| SA032 | DW | M | <200 | Sino-Americas | South America | Andean | Peru | NA |
| SA010 | DW | F | <200 | Sino-Americas | South America | Andean | Chile | Valparaiso |
| SA008 | DW | M | <200 | Sino-Americas | South America | Andean | Chile | NA |
| SA021 | DW | M | <200 | Sino-Americas | South America | Andean | Peru | NA |
| SA038 | DW | M | <200 | Sino-Americas | South America | Andean | Peru | Pasamayo |
| SA001 | DW | F | <200 | Sino-Americas | South America | SA Unkw | SA Unkw | NA |
| CA001 | DW | M | <200 | Sino-Americas | South America | Carribbean | Barbados | Arawak |
| NA065 | DW | F | <200 | Sino-Americas | North America | NA Unkw | NA Unkw | NA |
| NA024 | DW | F | <700 | Sino-Americas | North America | NA South West | United States | Ketchipawan |
| CA014 | DW | M | <200 | Sino-Americas | North America | Carribbean | Jamaica | NA |

| | | | | | | | | |
|---|---|---|---|---|---|---|---|---|
| NA015 | DW | F | <700 | Sino-Americas | North America | NA South West | United States | Ketchipawan |
| NA011 | DW | F | <700 | Sino-Americas | North America | NA South West | United States | Ketchipawan |
| SA045 | DW | F | <200 | Sino-Americas | South America | Andean | Peru | NA |
| SA019 | DW | M | <200 | Sino-Americas | South America | Andean | Peru | NA |
| NA52 | DW | M | <700 | Sino-Americas | North America | NA South West | United States | Ketchipawan |
| SA26 | DW | F | <200 | Sino-Americas | South America | Andean | Peru | NA |
| NA72 | DW | M | <200 | Sino-Americas | North America | NA Plains | United States | Sioux |
| SA23 | DW | F | <200 | Sino-Americas | South America | Andean | Peru | NA |
| NA001 | DW | F | <700 | Sino-Americas | North America | NA South West | United States | Ketchipawan |
| NA46 | DW | F | <700 | Sino-Americas | North America | NA South West | United States | Ketchipawan |
| SA39 | DW | M | <200 | Sino-Americas | South America | Andean | Peru | NA |
| SA37 | DW | M | <200 | Sino-Americas | South America | Andean | Peru | NA |
| SA44 | DW | F | <200 | Sino-Americas | South America | Andean | Peru | NA |
| SA17 | DW | M | <200 | Sino-Americas | South America | Andean | Peru | NA |
| SA007 | DW | M | <200 | Sino-Americas | South America | Andean | Chile | NA |
| SA020 | DW | F | <200 | Sino-Americas | South America | Andean | Peru | NA |
| NA002 | DW | F | <700 | Sino-Americas | North America | NA South West | United States | Ketchipawan |
| NA071 | DW | M | <200 | Sino-Americas | North America | NA South West | United States | Apache |
| NA034 | DW | F | <700 | Sino-Americas | North America | NA South West | United States | Ketchipawan |
| SA18 | DW | F | <200 | Sino-Americas | South America | Andean | Peru | NA |
| CA10 | DW | F | <200 | Sino-Americas | North America | Carribbean | Jamaica | NA |
| SA15 | DW | F | <200 | Sino-Americas | South America | Andean | Peru | NA |
| SA006 | DW | M | <200 | Sino-Americas | South America | Andean | Chile | NA |
| NA_12 | DW | F | <700 | Sino-Americas | North America | NA South West | United States | Ketchipawan |
| CA95 | DW | F | <200 | Sino-Americas | North America | Carribbean | Jamaica | NA |
| CA26 | DW | M | <200 | Sino-Americas | North America | Carribbean | Jamaica | NA |
| NA45 | DW | F | <700 | Sino-Americas | North America | NA South West | United States | Ketchipawan |
| SA16 | DW | M | <200 | Sino-Americas | South America | Andean | Peru | NA |
| SA25 | DW | M | <200 | Sino-Americas | South America | Andean | Peru | NA |
| NA68 | DW | F | <200 | Sino-Americas | North America | NA Plains | United States | NA |
| SA_58 | DW | F | <200 | Sino-Americas | South America | Andean | Peru | NA |
| POL_41 | DW | F | <200 | Sahul-Pacific | Oceania | Polynesia | New Zealand | Maori |
| POL_43 | DW | F | <200 | Sahul-Pacific | Oceania | Polynesia | New Zealand | Maori |
| POL_002 | DW | M | <200 | Sahul-Pacific | Oceania | Polynesia | New Zealand | Maori |
| POL_11 | DW | F | <200 | Sahul-Pacific | Oceania | Polynesia | New Zealand | Maori |
| POL_19 | DW | M | <200 | Sahul-Pacific | Oceania | Polynesia | New Zealand | Maori |
| POL_05 | DW | F | <200 | Sahul-Pacific | Oceania | Polynesia | New Zealand | Maori |
| POL_24 | DW | M | <200 | Sahul-Pacific | Oceania | Polynesia | New Zealand | Maori |
| POL_22 | DW | M | <200 | Sahul-Pacific | Oceania | Polynesia | New Zealand | Maori |
| POL_09 | DW | M | <200 | Sahul-Pacific | Oceania | Polynesia | New Zealand | Maori |
| POL_40 | DW | F | <200 | Sahul-Pacific | Oceania | Polynesia | New Zealand | Maori |
| NA_138 | DW | M | <200 | Sino-Americas | North America | American Arctic | American Arctic Unkw | Inuit |
| POL_15 | DW | M | <200 | Sahul-Pacific | Oceania | Polynesia | New Zealand | Maori |
| POL_23 | DW | F | <200 | Sahul-Pacific | Oceania | Polynesia | New Zealand | Maori |
| NA_139 | DW | F | <200 | Sino-Americas | North America | American Arctic | American Arctic Unkw | Inuit |
| POL_12 | DW | M | <200 | Sahul-Pacific | Oceania | Polynesia | New Zealand | Maori |
| NA_163 | DW | F | <200 | Sino-Americas | North America | American Arctic | American Arctic Unkw | Inuit |
| NA_147 | DW | M | <200 | Sino-Americas | North America | American Arctic | American Arctic Unkw | Inuit |
| POL_17 | DW | M | <200 | Sahul-Pacific | Oceania | Polynesia | New Zealand | Maori |
| NA_173 | DW | F | <200 | Sino-Americas | North America | NA Unkw | NA Unkw | NA |
| NA_110 | DW | NA | <200 | Sino-Americas | North America | NA Northwest Coast | Canada | Vancouver Island |
| NA_154 | DW | F | <200 | Sino-Americas | North America | American Arctic | American Arctic Unkw | Inuit |
| POL_45 | DW | M | <200 | Sahul-Pacific | Oceania | Polynesia | New Zealand | Maori |
| NA_149 | DW | M | <200 | Sino-Americas | North America | American Arctic | American Arctic Unkw | Inuit |
| NA_87 | DW | M | <200 | Sino-Americas | North America | NA Northwest Coast | Canada | New Westminster |
| POL_06 | DW | M | <200 | Sahul-Pacific | Oceania | Polynesia | New Zealand | Maori |

| | | | | | | | | |
|---|---|---|---|---|---|---|---|---|
| POL_18 | DW | M | <200 | Sahul-Pacific | Oceania | Polynesia | New Zealand | Maori |
| NA_137 | DW | F | <200 | Sino-Americas | North America | American Arctic | American Arctic Unkw | Inuit |
| POL_13 | DW | M | <200 | Sahul-Pacific | Oceania | Polynesia | New Zealand | Maori |
| NA_145 | DW | F | <200 | Sino-Americas | North America | American Arctic | American Arctic Unkw | Inuit |
| NA_67 | DW | F | <200 | Sino-Americas | North America | NA Northwest Coast | United States | NA |
| NA_133 | DW | F | <200 | Sino-Americas | North America | American Arctic | American Arctic Unkw | Inuit |
| NA_151 | DW | F | <200 | Sino-Americas | North America | American Arctic | American Arctic Unkw | Inuit |
| NA_123 | DW | M | <200 | Sino-Americas | North America | American Arctic | American Arctic Unkw | Inuit |
| NA_124 | DW | F | <200 | Sino-Americas | North America | American Arctic | Greenland | Inuit |
| NA_136 | DW | M | <200 | Sino-Americas | North America | American Arctic | American Arctic Unkw | Inuit |
| NA_140 | DW | M | <200 | Sino-Americas | North America | American Arctic | American Arctic Unkw | Inuit |
| NA_105 | DW | F | <200 | Sino-Americas | North America | NA Northwest Coast | Canada | Vancouver Island |
| NA_62 | DW | M | <200 | Sino-Americas | North America | NA South West | United States | Zuni |
| NA_135 | DW | M | <200 | Sino-Americas | North America | American Arctic | American Arctic Unkw | Inuit |
| NA_68 | DW | F | <200 | Sino-Americas | North America | NA Plains | United States | NA |
| NA_111 | DW | F | <200 | Sino-Americas | North America | NA Northwest Coast | United States | Makah |
| NA_150 | DW | F | <200 | Sino-Americas | North America | American Arctic | American Arctic Unkw | Inuit |
| NA_144 | DW | F | <200 | Sino-Americas | North America | American Arctic | American Arctic Unkw | Inuit |
| NA_121 | DW | F | <200 | Sino-Americas | North America | American Arctic | Greenland | Inuit Eleanoran Bay |
| NA_182 | DW | M | <200 | Sino-Americas | North America | NA Unkw | NA Unkw | Native American |
| NA_82 | DW | M | <200 | Sino-Americas | North America | NA Northeast Woodlands | Canada | Huron |
| POL_44 | DW | M | <200 | Sahul-Pacific | Oceania | Polynesia | New Zealand | Maori |
| NA_95 | DW | M | <200 | Sino-Americas | North America | NA Northwest Coast | Canada | Vancouver Island |
| NA_72 | DW | M | <200 | Sino-Americas | North America | NA Plains | United States | Sioux |
| NA_89 | DW | M | <200 | Sino-Americas | North America | NA Northwest Coast | Canada | New Westminster |
| NA_132 | DW | F | <200 | Sino-Americas | North America | American Arctic | American Arctic Unkw | Inuit |
| NA_75 | DW | F | <200 | Sino-Americas | North America | NA Northeast Woodlands | United States | Iroquois |
| NA_71 | DW | M | <200 | Sino-Americas | North America | NA South West | United States | Apache |
| NA_84 | DW | F | <200 | Sino-Americas | North America | NA Subarctic | Canada | Manitoba |
| NA_153 | DW | M | <200 | Sino-Americas | North America | American Arctic | American Arctic Unkw | Inuit |
| NA_98 | DW | F | <200 | Sino-Americas | North America | NA Northwest Coast | Canada | Vancouver Island |
| NA_164 | DW | F | <200 | Sino-Americas | North America | American Arctic | American Arctic Unkw | Inuit |
| NA_102 | DW | M | <200 | Sino-Americas | North America | NA Northwest Coast | Canada | Vancouver Island |
| NA_97 | DW | F | <200 | Sino-Americas | North America | NA Northwest Coast | Canada | Vancouver Island |
| NA_183 | DW | M | <200 | Sino-Americas | North America | NA Unkw | NA Unkw | Native American |

| | | | | | | | | |
|---|---|---|---|---|---|---|---|---|
| NA_61 | DW | F | <200 | Sino-Americas | North America | NA South West | United States | Zuni |
| NA_83 | DW | M | <200 | Sino-Americas | North America | NA Northeast Woodlands | Canada | Huron |
| NA_104 | DW | M | <200 | Sino-Americas | North America | NA Northwest Coast | Canada | Vancouver Island |
| NA_76 | DW | M | <200 | Sino-Americas | North America | NA Northeast Woodlands | United States | Iroquois |
| NA_101 | DW | F | <200 | Sino-Americas | North America | NA Northwest Coast | Canada | Vancouver Island |
| NA_92 | DW | M | <200 | Sino-Americas | North America | NA Northwest Coast | Canada | Vancouver Island |
| NA_81 | DW | M | <200 | Sino-Americas | North America | NA Northeast Woodlands | Canada | Huron |
| NA_74 | DW | M | <200 | Sino-Americas | North America | NA Northeast Woodlands | Canada | Huron |
| NA_134 | DW | F | <200 | Sino-Americas | North America | American Arctic | American Arctic Unkw | Inuit |
| NU_761 | DW | M | <200 | Sub-Saharan Africa | Sub-Saharan Africa | SSA Unkw | SSA Unkw | NA |
| AUS_001 | DW | M | <200 | Sahul-Pacific | Oceania | Australia | Australia Unkw | NA |
| AUS_016 | DW | M | <200 | Sahul-Pacific | Oceania | Australia | Australia Unkw | Aborigine |
| AUS_017 | DW | F | <200 | Sahul-Pacific | Oceania | Australia | Australia Unkw | NA |
| AUS_020 | DW | M | <200 | Sahul-Pacific | Oceania | Australia | Australia Unkw | NA |
| AUS_022 | DW | F | <200 | Sahul-Pacific | Oceania | Australia | Australia Unkw | NA |
| AUS_023 | DW | M | <200 | Sahul-Pacific | Oceania | Australia | Australia Unkw | NA |
| AUS_024 | DW | M | <200 | Sahul-Pacific | Oceania | Australia | Australia Unkw | Aborigine |
| AUS_025 | DW | M | <200 | Sahul-Pacific | Oceania | Australia | Australia Unkw | Aborigine |
| AUS_027 | DW | F | <200 | Sahul-Pacific | Oceania | Australia | Australia Unkw | NA |
| AUS_028 | DW | F | <200 | Sahul-Pacific | Oceania | Australia | Australia Unkw | NA |
| AUS_029 | DW | M | <200 | Sahul-Pacific | Oceania | Australia | Australia Unkw | NA |
| AUS_030 | DW | M | <200 | Sahul-Pacific | Oceania | Australia | Australia Unkw | Aborigine |
| AUS_032 | DW | M | <200 | Sahul-Pacific | Oceania | Australia | Australia Unkw | NA |
| AUS_037 | DW | M | <200 | Sahul-Pacific | Oceania | Australia | Victoria | NA |
| AUS_046 | DW | F | <200 | Sahul-Pacific | Oceania | Australia | Australia Unkw | Aborigine |
| AUS_047 | DW | F | <200 | Sahul-Pacific | Oceania | Australia | Australia Unkw | NA |
| AUS_048 | DW | F | <200 | Sahul-Pacific | Oceania | Australia | Australia Unkw | NA |
| AUS_049 | DW | F | <200 | Sahul-Pacific | Oceania | Australia | Australia Unkw | NA |
| AUS_050 | DW | F | <200 | Sahul-Pacific | Oceania | Australia | Australia Unkw | NA |
| AUS_051 | DW | M | <200 | Sahul-Pacific | Oceania | Australia | Australia Unkw | Aborigine |
| AUS_053 | DW | M | <200 | Sahul-Pacific | Oceania | Australia | Australia Unkw | Aborigine |
| AUS_054 | DW | F | <200 | Sahul-Pacific | Oceania | Australia | Australia Unkw | NA |
| AUS_055 | DW | M | <200 | Sahul-Pacific | Oceania | Australia | Australia Unkw | NA |
| AUS_056 | DW | F | <200 | Sahul-Pacific | Oceania | Australia | Australia Unkw | NA |
| AUS_058 | DW | M | <200 | Sahul-Pacific | Oceania | Australia | Australia Unkw | NA |
| AUS_059 | DW | F | <200 | Sahul-Pacific | Oceania | Australia | Western Australia | Baiono |
| AUS_062 | DW | M | <200 | Sahul-Pacific | Oceania | Australia | Western Australia | Perth |
| AUS_077 | DW | M | <200 | Sahul-Pacific | Oceania | Australia | Australia Unkw | Aborigine |
| AUS_078 | DW | M | <200 | Sahul-Pacific | Oceania | Australia | Australia Unkw | NA |
| AUS_079 | DW | F | <200 | Sahul-Pacific | Oceania | Australia | South Australia | NA |
| AUS_080 | DW | M | <200 | Sahul-Pacific | Oceania | Australia | South Australia | NA |
| AUS_081 | DW | M | <200 | Sahul-Pacific | Oceania | Australia | South Australia | NA |
| AUS_082 | DW | M | <200 | Sahul-Pacific | Oceania | Australia | South Australia | NA |
| AUS_083 | DW | F | <200 | Sahul-Pacific | Oceania | Australia | Australia Unkw | NA |
| AUS_093 | DW | F | <200 | Sahul-Pacific | Oceania | Australia | South Australia | NA |
| AUS_094 | DW | F | <200 | Sahul-Pacific | Oceania | Australia | South East Australia | NA |
| AUS_095 | DW | F | <200 | Sahul-Pacific | Oceania | Australia | South East Australia | NA |
| AUS_096 | DW | F | <200 | Sahul-Pacific | Oceania | Australia | Victoria | NA |
| AUS_102 | DW | M | <200 | Sahul-Pacific | Oceania | Australia | New South Wales | Aborigine |

| | | | | | | | | |
|---|---|---|---|---|---|---|---|---|
| AUS_104 | DW | M | <200 | Sahul-Pacific | Oceania | Australia | South East Australia | NA |
| AUS_105 | DW | M | <200 | Sahul-Pacific | Oceania | Australia | New South Wales | Aborigine |
| AUS_106 | DW | M | <200 | Sahul-Pacific | Oceania | Australia | Australia Unkw | NA |
| AUS_107 | DW | F | <200 | Sahul-Pacific | Oceania | Australia | New South Wales | Mem Mem |
| AUS_108 | DW | M | <200 | Sahul-Pacific | Oceania | Australia | New South Wales | Berida |
| AUS_109 | DW | M | <200 | Sahul-Pacific | Oceania | Australia | New South Wales | Wollongong |
| AUS_110 | DW | F | <200 | Sahul-Pacific | Oceania | Australia | New South Wales | Newcastle |
| AUS_113 | DW | M | <200 | Sahul-Pacific | Oceania | Australia | New South Wales | Murray River |
| AUS_114_1 | DW | M | <200 | Sahul-Pacific | Oceania | Australia | New South Wales | Murray River |
| AUS_116 | DW | F | <200 | Sahul-Pacific | Oceania | Australia | New South Wales | Murray River |
| AUS_119 | DW | F | <200 | Sahul-Pacific | Oceania | Australia | New South Wales | Murray River |
| AUS_120 | DW | M | <200 | Sahul-Pacific | Oceania | Australia | New South Wales | Murray River |
| AUS_121 | DW | F | <200 | Sahul-Pacific | Oceania | Australia | New South Wales | Murray River |
| AUS_122 | DW | M | <200 | Sahul-Pacific | Oceania | Australia | Queensland | NA |
| AUS_123 | DW | F | <200 | Sahul-Pacific | Oceania | Australia | Queensland | Mackay Aborigine |
| AUS_124 | DW | M | <200 | Sahul-Pacific | Oceania | Australia | Queensland | Aborigine |
| AUS_125 | DW | F | <200 | Sahul-Pacific | Oceania | Australia | Queensland | Croydon Queensland |
| AUS_126 | DW | M | <200 | Sahul-Pacific | Oceania | Australia | Queensland | Croydon Queensland |
| AUS_127 | DW | M | <200 | Sahul-Pacific | Oceania | Australia | Queensland | North Queensland |
| AUS_128 | DW | M | <200 | Sahul-Pacific | Oceania | Australia | Queensland | North Queensland |
| AUS_129 | DW | M | <200 | Sahul-Pacific | Oceania | Australia | Queensland | North Queensland |
| AUS_130 | DW | F | <200 | Sahul-Pacific | Oceania | Australia | Queensland | North Queensland |
| AUS_131 | DW | F | <200 | Sahul-Pacific | Oceania | Australia | Queensland | North Queensland |
| AF_11_5_28 | DW | F | 6400-6000 | West Eurasia | North Africa | North East Africa | Egypt | Badari |
| AF_11_5_21 | DW | F | 6400-6000 | West Eurasia | North Africa | North East Africa | Egypt | Badari |
| AF_11_5_42 | DW | M | 6400-6000 | West Eurasia | North Africa | North East Africa | Egypt | Badari |
| AF_11_5_04 | DW | F | 6400-6000 | West Eurasia | North Africa | North East Africa | Egypt | Badari |
| AF_11_5_17 | DW | F | 6400-6000 | West Eurasia | North Africa | North East Africa | Egypt | Badari |
| AF_11_5_10 | DW | M | 6400-6000 | West Eurasia | North Africa | North East Africa | Egypt | Badari |
| AF_11_5_40 | DW | M | 6400-6000 | West Eurasia | North Africa | North East Africa | Egypt | Badari |
| AF_12_4_25 | DW | M | 2100-1400 | West Eurasia | North Africa | North East Africa | Sudan | Jebel Moya |
| AF_11_5_22 | DW | F | 6400-6000 | West Eurasia | North Africa | North East Africa | Egypt | Badari |
| AF_11_5_25 | DW | F | 6400-6000 | West Eurasia | North Africa | North East Africa | Egypt | Badari |
| AF_12_4_28 | DW | M | 2100-1400 | West Eurasia | North Africa | North East Africa | Sudan | Jebel Moya |
| AF_12_4_18 | DW | M | 2100-1400 | West Eurasia | North Africa | North East Africa | Sudan | Jebel Moya |
| AF_12_4_08 | DW | M | 2100-1400 | West Eurasia | North Africa | North East Africa | Sudan | Jebel Moya |
| AF_11_5_27 | DW | F | 6000-5200 | West Eurasia | North Africa | North East Africa | Egypt | Nagada |
| AF_11_5_59 | DW | M | 6000-5200 | West Eurasia | North Africa | North East Africa | Egypt | Nagada |
| AF_11_5_55 | DW | M | 6000-5200 | West Eurasia | North Africa | North East Africa | Egypt | Nagada |

| | | | | | | | | |
|---|---|---|---|---|---|---|---|---|
| AF_11_5_34 | DW | M | 6000-5200 | West Eurasia | North Africa | North East Africa | Egypt | Nagada |
| AF_11_5_53 | DW | M | 6000-5200 | West Eurasia | North Africa | North East Africa | Egypt | Nagada |
| AF_11_5_02 | DW | F | 6000-5200 | West Eurasia | North Africa | North East Africa | Egypt | Nagada |
| AF_12_4_07 | DW | M | 6000-5200 | West Eurasia | North Africa | North East Africa | Egypt | Nagada |
| AF_11_5_12 | DW | F | 6000-5200 | West Eurasia | North Africa | North East Africa | Egypt | Nagada |
| AF_12_4_15 | DW | M | 2100-1400 | West Eurasia | North Africa | North East Africa | Sudan | Jebel Moya |
| AF_12_4_103 | DW | M | 2100-1400 | West Eurasia | North Africa | North East Africa | Sudan | Jebel Moya |
| AF_11_5_07 | DW | F | 6000-5200 | West Eurasia | North Africa | North East Africa | Egypt | Nagada |
| AF_11_5_44 | DW | M | 6000-5200 | West Eurasia | North Africa | North East Africa | Egypt | Nagada |
| AF_11_5_54 | DW | F | 6000-5200 | West Eurasia | North Africa | North East Africa | Egypt | Nagada |
| AF_11_5_24 | DW | F | 6000-5200 | West Eurasia | North Africa | North East Africa | Egypt | Nagada |
| AF_11_5_32 | DW | M | 6000-5200 | West Eurasia | North Africa | North East Africa | Egypt | Nagada |
| AF_11_5_43 | DW | M | 6000-5200 | West Eurasia | North Africa | North East Africa | Egypt | Nagada |
| AF_11_5_16 | DW | M | 6000-5200 | West Eurasia | North Africa | North East Africa | Egypt | Nagada |
| AF_12_4_91 | DW | M | 2100-1400 | West Eurasia | North Africa | North East Africa | Sudan | Jebel Moya |
| AF_11_5_18 | DW | M | 6000-5200 | West Eurasia | North Africa | North East Africa | Egypt | Nagada |
| AF_12_4_21 | DW | F | 2100-1400 | West Eurasia | North Africa | North East Africa | Sudan | Jebel Moya |
| AF_11_5_41 | DW | F | 6000-5200 | West Eurasia | North Africa | North East Africa | Egypt | Nagada |
| AF_11_5_52 | DW | F | 6000-5200 | West Eurasia | North Africa | North East Africa | Egypt | Nagada |
| AF_12_4_17 | DW | NA | 2100-1400 | West Eurasia | North Africa | North East Africa | Sudan | Jebel Moya |
| AF_12_4_19 | DW | M | 2100-1400 | West Eurasia | North Africa | North East Africa | Sudan | Jebel Moya |
| AF_11_5_58 | DW | F | 6000-5200 | West Eurasia | North Africa | North East Africa | Egypt | Nagada |
| AF_12_4_10 | DW | M | 2100-1400 | West Eurasia | North Africa | North East Africa | Sudan | Jebel Moya |
| AF_11_5_39 | DW | M | 6000-5200 | West Eurasia | North Africa | North East Africa | Egypt | Nagada |
| AF_11_5_46 | DW | F | 6000-5200 | West Eurasia | North Africa | North East Africa | Egypt | Nagada |
| AF_11_5_33 | DW | F | 6000-5200 | West Eurasia | North Africa | North East Africa | Egypt | Nagada |
| AF_11_5_47 | DW | F | 6000-5200 | West Eurasia | North Africa | North East Africa | Egypt | Nagada |
| AF_11_5_35 | DW | F | 6000-5200 | West Eurasia | North Africa | North East Africa | Egypt | Nagada |
| AF_11_5_37 | DW | M | 6000-5200 | West Eurasia | North Africa | North East Africa | Egypt | Nagada |
| AF_11_536 | DW | M | 6000-5200 | West Eurasia | North Africa | North East Africa | Egypt | Nagada |
| AF_11_5_57 | DW | M | 6000-5200 | West Eurasia | North Africa | North East Africa | Egypt | Nagada |
| AF_12_4_22 | DW | F | 2100-1400 | West Eurasia | North Africa | North East Africa | Sudan | Jebel Moya |
| AF_11_5_26 | DW | F | 6000-5200 | West Eurasia | North Africa | North East Africa | Egypt | Nagada |
| AF_11_5_29 | DW | F | 6000-5200 | West Eurasia | North Africa | North East Africa | Egypt | Nagada |
| AF_12_4_26 | DW | M | 2100-1400 | West Eurasia | North Africa | North East Africa | Sudan | Jebel Moya |
| AF_12_4_108 | DW | M | 2100-1400 | West Eurasia | North Africa | North East Africa | Sudan | Jebel Moya |
| AF_11_5_31 | DW | F | 6000-5200 | West Eurasia | North Africa | North East Africa | Egypt | Nagada |
| AF_11_5_56 | DW | F | 6000-5200 | West Eurasia | North Africa | North East Africa | Egypt | Nagada |

| | | | | | | | | |
|---|---|---|---|---|---|---|---|---|
| AF_11_5_38 | DW | F | 6000-5200 | West Eurasia | North Africa | North East Africa | Egypt | Nagada |
| SUD_11 | DW | M | 3750-3500 | West Eurasia | North Africa | North East Africa | Sudan | Kerma |
| SUD_05 | DW | M | 3750-3500 | West Eurasia | North Africa | North East Africa | Sudan | Kerma |
| SUD_13 | DW | M | 3750-3500 | West Eurasia | North Africa | North East Africa | Sudan | NA |
| SUD_5343 | DW | M | 3750-3500 | West Eurasia | North Africa | North East Africa | Sudan | NA |
| SUD_08 | DW | M | 3750-3500 | West Eurasia | North Africa | North East Africa | Sudan | Kerma |
| SUD_14 | DW | M | 3750-3500 | West Eurasia | North Africa | North East Africa | Sudan | Kerma |
| SUD_04 | DW | M | 3750-3500 | West Eurasia | North Africa | North East Africa | Sudan | Kerma |
| SUD_20 | DW | M | 3750-3500 | West Eurasia | North Africa | North East Africa | Sudan | Kerma |
| SUD_10 | DW | M | 3750-3500 | West Eurasia | North Africa | North East Africa | Sudan | Kerma |
| SUD_01 | DW | M | 3750-3500 | West Eurasia | North Africa | North East Africa | Sudan | Kerma |
| SUD_26 | DW | M | 3750-3500 | West Eurasia | North Africa | North East Africa | Sudan | Kerma |
| SUD_38 | DW | M | 3750-3500 | West Eurasia | North Africa | North East Africa | Sudan | Kerma |
| SUD_16 | DW | M | 3750-3500 | West Eurasia | North Africa | North East Africa | Sudan | Kerma |
| SUD_27 | DW | M | 3750-3500 | West Eurasia | North Africa | North East Africa | Sudan | Kerma |
| SUD_15 | DW | M | 3750-3500 | West Eurasia | North Africa | North East Africa | Sudan | Kerma |
| SUD_02 | DW | M | 3750-3500 | West Eurasia | North Africa | North East Africa | Sudan | Kerma |
| SUD_5053 | DW | F | <200 | West Eurasia | North Africa | North East Africa | Sudan | NA |
| SUD_4438 | DW | M | <200 | West Eurasia | North Africa | North East Africa | Sudan | NA |
| MEL_255 | DW | F | <200 | Sahul-Pacific | Oceania | Melanesia | Papua New Guinea | Muyuw Kwaiawata Island |
| AF_11_5_48 | DW | NA | <200 | West Eurasia | North Africa | North East Africa | Egypt | NA |
| 4434 | DW | M | 200 | Sub-Saharan Africa | Sub-Saharan Africa | Western Africa | Ghana | Ashanti |
| 5419 | DW | F | <200 | Sub-Saharan Africa | Sub-Saharan Africa | Western Africa | Nigeria | Kagoro |
| AF_21_0_14 | DW | M | <200 | Sub-Saharan Africa | Sub-Saharan Africa | Eastern Africa | Kenya | Teita |
| AF_21_0_62 | DW | F | <200 | Sub-Saharan Africa | Sub-Saharan Africa | Eastern Africa | Kenya | Teita |
| AF_21_0_102 | DW | F | <200 | Sub-Saharan Africa | Sub-Saharan Africa | Eastern Africa | Kenya | Teita |
| AF_21_0_100 | DW | F | <200 | Sub-Saharan Africa | Sub-Saharan Africa | Eastern Africa | Kenya | Teita |
| AF_21_0_48 | DW | M | <200 | Sub-Saharan Africa | Sub-Saharan Africa | Eastern Africa | Kenya | Teita |
| AF_21_0_108 | DW | F | <200 | Sub-Saharan Africa | Sub-Saharan Africa | Eastern Africa | Kenya | Teita |
| AF_21_0_107 | DW | F | <200 | Sub-Saharan Africa | Sub-Saharan Africa | Eastern Africa | Kenya | Teita |
| AF_21_0_93 | DW | F | <200 | Sub-Saharan Africa | Sub-Saharan Africa | Eastern Africa | Kenya | Teita |
| AF_21_0_95 | DW | F | <200 | Sub-Saharan Africa | Sub-Saharan Africa | Eastern Africa | Kenya | Teita |
| AF_21_0_50 | DW | M | <200 | Sub-Saharan Africa | Sub-Saharan Africa | Eastern Africa | Kenya | Teita |
| AF_21_0_45 | DW | M | <200 | Sub-Saharan Africa | Sub-Saharan Africa | Eastern Africa | Kenya | Teita |
| AF_21_0_68 | DW | M | <200 | Sub-Saharan Africa | Sub-Saharan Africa | Eastern Africa | Kenya | Teita |
| AF_21_0_71 | DW | F | <200 | Sub-Saharan Africa | Sub-Saharan Africa | Eastern Africa | Kenya | Teita |
| AF_21_0_41 | DW | M | <200 | Sub-Saharan Africa | Sub-Saharan Africa | Eastern Africa | Kenya | Teita |

| | | | | | | | | |
|---|---|---|---|---|---|---|---|---|
| AF_21_0_82 | DW | F | <200 | Sub-Saharan Africa | Sub-Saharan Africa | Eastern Africa | Kenya | Teita |
| AF_21_0_94 | DW | F | <200 | Sub-Saharan Africa | Sub-Saharan Africa | Eastern Africa | Kenya | Teita |
| AF_21_0_67 | DW | F | <200 | Sub-Saharan Africa | Sub-Saharan Africa | Eastern Africa | Kenya | Teita |
| AF_21_0_70 | DW | F | <200 | Sub-Saharan Africa | Sub-Saharan Africa | Eastern Africa | Kenya | Teita |
| AF_21_0_60 | DW | M | <200 | Sub-Saharan Africa | Sub-Saharan Africa | Eastern Africa | Kenya | Teita |
| AF_21_0_34 | DW | M | <200 | Sub-Saharan Africa | Sub-Saharan Africa | Eastern Africa | Kenya | Teita |
| AF_21_0_24 | DW | M | <200 | Sub-Saharan Africa | Sub-Saharan Africa | Eastern Africa | Kenya | Teita |
| AF_21_0_9 | DW | M | <200 | Sub-Saharan Africa | Sub-Saharan Africa | Eastern Africa | Kenya | Teita |
| AF_21_0_20 | DW | M | <200 | Sub-Saharan Africa | Sub-Saharan Africa | Eastern Africa | Kenya | Teita |
| AF_21_0_18 | DW | M | <200 | Sub-Saharan Africa | Sub-Saharan Africa | Eastern Africa | Kenya | Teita |
| AF_21_0_43 | DW | M | <200 | Sub-Saharan Africa | Sub-Saharan Africa | Eastern Africa | Kenya | Teita |
| AF_21_0_16 | DW | M | <200 | Sub-Saharan Africa | Sub-Saharan Africa | Eastern Africa | Kenya | Teita |
| AF_21_0_13 | DW | M | <200 | Sub-Saharan Africa | Sub-Saharan Africa | Eastern Africa | Kenya | Teita |
| AF_21_0_40 | DW | M | <200 | Sub-Saharan Africa | Sub-Saharan Africa | Eastern Africa | Kenya | Teita |
| AF_21_0_25 | DW | M | <200 | Sub-Saharan Africa | Sub-Saharan Africa | Eastern Africa | Kenya | Teita |
| AF_21_0_12 | DW | M | <200 | Sub-Saharan Africa | Sub-Saharan Africa | Eastern Africa | Kenya | Teita |
| AF_21_0_29 | DW | M | <200 | Sub-Saharan Africa | Sub-Saharan Africa | Eastern Africa | Kenya | Teita |
| AF_21_0_118 | DW | F | <200 | Sub-Saharan Africa | Sub-Saharan Africa | Eastern Africa | Kenya | Teita |
| AF_21_0_121 | DW | F | <200 | Sub-Saharan Africa | Sub-Saharan Africa | Eastern Africa | Kenya | Teita |
| AF_21_0_133 | DW | F | <200 | Sub-Saharan Africa | Sub-Saharan Africa | Eastern Africa | Kenya | Teita |
| AF_21_0_129 | DW | F | <200 | Sub-Saharan Africa | Sub-Saharan Africa | Eastern Africa | Kenya | Teita |
| AF_21_0_123 | DW | F | <200 | Sub-Saharan Africa | Sub-Saharan Africa | Eastern Africa | Kenya | Teita |
| AF_21_0_126 | DW | F | <200 | Sub-Saharan Africa | Sub-Saharan Africa | Eastern Africa | Kenya | Teita |
| AF_21_0_113 | DW | M | <200 | Sub-Saharan Africa | Sub-Saharan Africa | Eastern Africa | Kenya | Teita |
| 1742 | DW | F | <200 | Sub-Saharan Africa | Sub-Saharan Africa | Southern Africa | South Africa | NA |
| AF_23_0_28 | DW | F | <200 | Sub-Saharan Africa | Sub-Saharan Africa | Eastern Africa | Tanzania | Bukoba |
| AF_15_0_2 | DW | M | <200 | Sub-Saharan Africa | Sub-Saharan Africa | Eastern Africa | Somalia | Hariya |
| AF112 | DW | F | <200 | Sub-Saharan Africa | Sub-Saharan Africa | Southern Africa | South Africa | NA |
| SAS_14 | DW | M | <200 | West Eurasia | South Asia | Indian Sub-Continent | Pakistan | NA |
| 209305 | NMNH | M | <200 | Sunda-Pacific | South East Asia | Malay Archipelago | Philippines | Tagalog Island |
| 209307 | NMNH | F | <200 | Sunda-Pacific | South East Asia | Malay Archipelago | Philippines | Tagalog Island |
| 209310 | NMNH | F | <200 | Sunda-Pacific | South East Asia | Malay Archipelago | Philippines | Tagalog Island |
| 226155 | NMNH | F | <200 | Sahul-Pacific | Oceania | Polynesia | New Zealand | NA |
| 226156 | NMNH | F | <200 | Sahul-Pacific | Oceania | Polynesia | New Zealand | NA |
| 226158 | NMNH | M | <200 | Sahul-Pacific | Oceania | Polynesia | New Zealand | NA |
| 222001 | NMNH | M | <200 | Sunda-Pacific | South East Asia | Malay Archipelago | Indonesia | Pagi Island |
| 381079 | NMNH | F | <200 | Sahul-Pacific | Oceania | Melanesia | Papua New Guinea | New Britain |
| 99_1_109 | NMNH | M | ~1600 | Sino-Americas | North America | American Arctic | Alaska | Ipiutak |
| 99_1_199 | NMNH | F | ~1600 | Sino-Americas | North America | American Arctic | Alaska | Ipiutak |

| | | | | | | | | |
|---|---|---|---|---|---|---|---|---|
| 99_1_209 | NMNH | M | ~1600 | Sino-Americas | North America | American Arctic | Alaska | Ipiutak |
| 99_1_211 | NMNH | F | ~1600 | Sino-Americas | North America | American Arctic | Alaska | Ipiutak |
| 99_1_174 | NMNH | F | ~1600 | Sino-Americas | North America | American Arctic | Alaska | NA |
| 99_1_568 | NMNH | M | ~1600 | Sino-Americas | North America | American Arctic | Alaska | NA |
| 99_1_594 | NMNH | M | ~1600 | Sino-Americas | North America | American Arctic | Alaska | NA |
| 99_1_593 | NMNH | F | ~1600 | Sino-Americas | North America | American Arctic | Alaska | NA |
| 99_1_64 | NMNH | F | ~800 | Sino-Americas | North America | American Arctic | Alaska | Tigara |
| 99_1_642 | NMNH | M | ~800 | Sino-Americas | North America | American Arctic | Alaska | Tigara |
| 99_1_649 | NMNH | M | ~800 | Sino-Americas | North America | American Arctic | Alaska | Tigara |
| 99_1_222 | NMNH | F | ~800 | Sino-Americas | North America | American Arctic | Alaska | Tigara |
| 99_1_231 | NMNH | M | ~800 | Sino-Americas | North America | American Arctic | Alaska | Tigara |
| 99_1_231A | NMNH | F | ~800 | Sino-Americas | North America | American Arctic | Alaska | Tigara |
| 99_1_232 | NMNH | F | ~800 | Sino-Americas | North America | American Arctic | Alaska | Tigara |
| 99_1_260 | NMNH | F | ~800 | Sino-Americas | North America | American Arctic | Alaska | Tigara |
| 99_1_262 | NMNH | F | ~800 | Sino-Americas | North America | American Arctic | Alaska | Tigara |
| 99_1_402 | NMNH | M | ~800 | Sino-Americas | North America | American Arctic | Alaska | Tigara |
| 99_1_403 | NMNH | M | ~800 | Sino-Americas | North America | American Arctic | Alaska | Tigara |
| 99_1_404 | NMNH | M | ~800 | Sino-Americas | North America | American Arctic | Alaska | Tigara |
| 99_1_415 | NMNH | F | ~800 | Sino-Americas | North America | American Arctic | Alaska | Tigara |
| 99_1_420 | NMNH | F | ~800 | Sino-Americas | North America | American Arctic | Alaska | Tigara |
| 99_1_435 | NMNH | M | ~800 | Sino-Americas | North America | American Arctic | Alaska | Tigara |
| 99_1_462 | NMNH | M | ~800 | Sino-Americas | North America | American Arctic | Alaska | Tigara |
| 99_1_503 | NMNH | F | ~800 | Sino-Americas | North America | American Arctic | Alaska | Tigara |
| 226098 | NMNH | NA | <200 | Sahul-Pacific | Oceania | Melanesia | Papua New Guinea | New Britain |
| 227454 | NMNH | NA | <200 | Sahul-Pacific | Oceania | Melanesia | Papua New Guinea | New Britain |
| 227459 | NMNH | NA | <200 | Sahul-Pacific | Oceania | Melanesia | Papua New Guinea | New Britain |
| 227465 | NMNH | NA | <200 | Sahul-Pacific | Oceania | Melanesia | Papua New Guinea | New Britain |
| 242755 | NMNH | NA | <200 | Sino-Americas | North America | American Arctic | Canada | Baffin Island |
| 242834 | NMNH | NA | <200 | Sino-Americas | North America | American Arctic | Canada | Baffin Island |
| 259354 | NMNH | NA | <200 | Sunda-Pacific | South East Asia | Malay Archipelago | Philippines | NA |
| 342024 | NMNH | NA | Unkw | Sino-Americas | North America | American Arctic | Greenland | Inuit |
| 379058 | NMNH | NA | <200 | Sunda-Pacific | South East Asia | Malay Archipelago | Philippines | NA |
| 380430 | NMNH | F | <200 | Sunda-Pacific | South East Asia | Malay Archipelago | NA | Malaysian |
| 380447 | NMNH | M | <200 | Sunda-Pacific | South East Asia | Malay Archipelago | NA | NA |
| 380448 | NMNH | M | <200 | Sunda-Pacific | South East Asia | Malay Archipelago | NA | Malaysian |
| 380450 | NMNH | M | Unkw | Sahul-Pacific | Oceania | Australia | Northern Territory | Crocodile Island |

**ID** = collection identification number, **M** =Male, **F** = female, ***BP** = before present, **DW** = Duckworth, **AMNH** = American Museum of Natural History, **NMNH** = Smithsonian National Museum of Natural History, **†G1** = Major Human Subdivisions, **G2** = Continental Group, **G3** = Continental Region, **G4** = Country/State, **G5** = Locality/Tribe, **Unkw** = unknown
